# A multi-measure approach for assessing the performance of fMRI preprocessing strategies in resting-state functional connectivity

**DOI:** 10.1101/837609

**Authors:** Michalis Kassinopoulos, Georgios D. Mitsis

## Abstract

It is well established that head motion and physiological processes (e.g. cardiac and breathing activity) should be taken into consideration when analyzing and interpreting results in fMRI studies. However, even though recent studies aimed to evaluate the performance of different preprocessing pipelines there is still no consensus on the *optimal* strategy. This is partly due to the fact that the quality control (QC) metrics used to evaluate differences in performance across pipelines have often yielded contradictory results. Furthermore, preprocessing techniques based on physiological recordings or data decomposition techniques (e.g. aCompCor) have not been comprehensively examined. Here, to address the aforementioned issues, we propose a framework that summarizes the scores from eight previously proposed and novel QC metrics to a reduced set of two QC metrics that reflect the signal-to-noise ratio and the reduction in motion artifacts and biases in the preprocessed fMRI data. Using this framework, we evaluate the performance of three commonly used practices on the quality of data: 1) Removal of nuisance regressors from fMRI data, 2) discarding motion-contaminated volumes (i.e., scrubbing) before regression, and 3) low-pass filtering the data and the nuisance regressors before their removal. Using resting-state fMRI data from the Human Connectome Project, we show that the scores of the examined QC metrics improve the most when the global signal (GS) and about 17% of principal components from white matter (WM) are removed from the data. Finally, we observe a small further improvement with low-pass filtering at 0.20 Hz and milder variants of WM denoising, but not with scrubbing.

## 1. Introduction

Functional connectivity (FC) using resting-state functional magnetic resonance imaging (fMRI) has attracted much attention since Biswal and colleagues first demonstrated that, during rest, the blood-oxygen-level-dependent (BOLD) signals in distinct areas of the somatomotor network are temporally correlated (Biswal et al., 1995). Strategies for studying resting-state FC have advanced in the last two decades, allowing the identification of large-scale functional networks, termed resting-state networks (RSNs; Fox et al., 2005; Smith et al., 2013b) that are also known to activate (or deactivate in the case of the default mode network) on a range of tasks (Raichle et al., 2001; Smith et al., 2009). The whole-brain functional connectivity (FC) is similar across subjects but at the same time has a subject-specific component as well. (Finn et al., 2015). FC estimates have been shown to predict behavioral measures in individuals (Smith et al., 2015), while differences in FC have been reported in patients with a range of cerebrovascular and mental disorders compared to healthy subjects (Demirtaş et al., 2016; Leonardi et al., 2013; Woodward and Cascio, 2015). These findings suggest that FC has the potential to improve our understanding regarding the functional organization over development, aging, and diseases states, as well as assist in the development of new biomarkers.

However, a main problem in fMRI is that significant variance on the BOLD signal is driven by head motion, which has been shown to have severe consequences on FC studies (Power et al., 2015; Satterthwaite et al., 2019). Motion artifacts tend to be more similar in nearby regions compared to distant regions (Power et al., 2012; Satterthwaite et al., 2012; van Dijk et al., 2012). As a result, correlations between regions that are close to each other (short-distance correlations) tend to be inflated by motion more compared to distant regions (long-distance correlations; see for example Fig. 5 in Satterthwaite et al., 2013). Consequently, in studies comparing differences in FC between populations that exhibit different levels of motion, motion artifacts can cause artificial differences in FC between the examined populations. Therefore, this should be carefully considered in the preprocessing and analysis of the data. This phenomenon is particularly problematic for studies of development, aging and disease as children, elderly and patients tend to move more during the scan than young or control subjects (Power et al., 2015).

Importantly, confounds in fMRI arise also from physiological noise (Caballero-Gaudes and Reynolds, 2017; Liu, 2016; Murphy et al., 2013). Cardiac pulsatility in large vessels caused by cardiac-related pressure changes generates small movements in and around large vessels. In turn, these movements introduce fast pseudo-periodic fluctuations (~1 Hz) in the BOLD signal (Dagli et al., 1999), particularly in areas around the brainstem as well as areas in the superior sagittal sinus and lateral sulcus. Similarly, breathing motion introduces high-frequency artifacts (~0.3 Hz) particularly at the edges of the brain. Slow-frequency fluctuations in heart rate and breathing pattern (<0.1 Hz) are also typically observed during rest which have a direct effect on the levels of oxygenated hemoglobin in the brain (Birn et al., 2006; Chang et al., 2009; Kassinopoulos and Mitsis, 2021, 2019; Shmueli et al., 2007). As a result, they affect widespread regions in the gray matter (GM). Group-level statistical maps generated in our previous work with areas affected by the aforementioned physiological processes are available on https://neurovault.org/collections/5654/ (Fig. 12 in Kassinopoulos and Mitsis, 2019). Finally, widespread regions in GM are also prone to artifacts induced by slow spontaneous fluctuations in levels of arterial carbon dioxide (Prokopiou et al., 2019; Wise et al., 2004) and blood pressure (Whittaker et al., 2019). fMRI fluctuations induced by changes in physiological signals are of particular concern as there is accumulating evidence that physiological processes can considerably affect FC estimates if not taken into account during the preprocessing stage (Birn, 2012; Birn et al., 2008a; Chang and Glover, 2009; Chen et al., 2020; Tong et al., 2019).

Several noise correction techniques (NCTs) have been proposed to correct for head motion artifacts and physiological noise that can be classified as model-based or model-free techniques. In the case of head motion, model-based techniques are based on the motion parameters (MPs) estimated from volume realignment performed in the initial steps of preprocessing. Three translational and three rotational displacement parameters are estimated from volume realignment that describe the rigid-body movement of head in space, yielding in total 6 MPs. The most common practice used in FC studies to account for motion is to remove the 6 MPs from the data through linear regression (Power et al., 2015). Sometimes the derivatives of the 6 MPs or even the squared terms of these 12 time series are also removed from the data (Satterthwaite et al., 2013). Another practice employed in recent studies, termed scrubbing, is to identify volumes contaminated by strong motion artifacts and discard them from the data or replace them with values from the adjacent volumes using interpolation (Lemieux et al., 2007; Power et al., 2015).

With regards to physiological noise, model-based techniques utilize physiological recordings collected during the fMRI scan. Typically, cardiac and breathing activity are recorded through a pulse oximeter and a respiratory bellow respectively, and are used to model their associated artifacts. The RETROICOR model proposed by Glover et al. (2000) employs the physiological recordings to generate nuisance regressors of sinusoidal signals that are in phase with the cardiac and breathing cycle. Subsequently, the extracted nuisance regressors are removed from the data through linear regression to account for artifacts related to cardiac pulsatility and breathing motion. Similarly, the convolution models proposed by Birn et al. (2008) and Chang et al. (2009) employ the physiological signals to extract heart rate and respiratory measures which are subsequently convolved with the so-called cardiac and respiration response functions. The outputs of these convolutions are used as nuisance regressors to account for the effects of heart rate and breathing (Birn et al., 2008b, 2006; Chang et al., 2009; Kassinopoulos and Mitsis, 2019).

An alternative option for noise correction in fMRI are model-free techniques that, in contrast to model-based techniques, do not require external physiological recordings and, have the theoretical benefit to be independent of a pre-established model. Some model-free techniques are based on principal component analysis (PCA) or independent component analysis (ICA), which decompose the fMRI data into a number of components (Behzadi et al., 2007; Perlbarg et al., 2007; Pruim et al., 2015b; Salimi-Khorshidi et al., 2014). Following this, components associated to noise are identified based on criteria such as their spatial pattern or frequency content and are subsequently removed from the data. Furthermore, low-pass filtering at about 0.08 Hz is commonly used to remove high-frequency noise as RSNs are known to exhibit slow oscillations below 0.1 Hz (Damoiseaux et al., 2006). Finally, the mean time series across voxels in the whole brain, referred to as global signal (GS), as well as mean time series from voxels in the three tissue compartments, GM, white matter (WM) and cerebrospinal fluid (CSF), are sometimes considered as nuisance regressors to account for global artifacts (Power et al., 2017).

Even though combining a variety of NCTs and removing a large set of nuisance regressors from the data may effectively suppress the effects of motion and physiological noise, this approach also leads to a reduction in the degrees of freedom in the data and potentially a loss in the signal of interest. Due to this, the set of regressors chosen for a particular dataset needs to be considered in conjunction with the degrees of freedom in the data, which in turn depend on the duration of the data and sampling frequency. Importantly, preprocessing steps that often precede the removal of nuisance regressors, such as temporal filtering and scrubbing, can also diminish the effective degrees of freedom in the data and increase the likelihood for spurious connectivity (Bright et al., 2017; Yan et al., 2013), which further complicates the task of choosing the *optimal* preprocessing pipeline.

Recent studies have attempted to compare the performance of a variety of NCTs as well as preprocessing pipelines that consist of a combination of techniques mentioned earlier (Birn et al., 2014; Burgess et al., 2016; Ciric et al., 2017; Parkes et al., 2018). A number of quality control (QC) metrics reflecting properties such as the identifiability of RSNs or the mitigation of motion effects were used in these studies. However, a common finding in these studies is that the scores obtained from the QC metrics for the examined NCTs or pipelines often yielded contradictory results. For example, pipelines yielding the highest score in terms of RSN identifiability were found to be less successful in reducing motion artifacts (Ciric et al., 2017). Moreover, even though there is strong evidence that model-free techniques based on PCA or ICA are able to reduce artifacts due to head motion or physiological noise (Behzadi et al., 2007; Muschelli et al., 2014; Pruim et al., 2015a; Salimi-Khorshidi et al., 2014), it is still an open question whether combining them with model-based techniques can result in superior performance.

In this work, we propose a framework for summarizing the scores of different signal and motion-related QC metrics by weighting these metrics according to their sensitivity. We define the sensitivity of a metric as the consistency of its scores across subgroups of subjects with similar levels of motion. Subsequently, using this multi-measure approach for assessing fMRI data quality, we compare the performance of several NCTs on multi-session resting-state fMRI data provided by the Human Connectome Project (Van Essen et al., 2013). The comparison of NCTs is done in a series of stages that allows us to better understand the role of each technique (e.g. low-pass filtering) in noise correction. With respect to model-free approaches, we examine FIX (“FMRIB’s ICA-based X-noisefier”; Salimi-Khorshidi et al., 2014) as well as variants of aCompCor (Behzadi et al., 2007). FIX consists of whole-brain ICA decomposition followed by removal of noisy components identified using a multi-level classifier (Salimi-Khorshidi et al., 2014). Besides evaluating the performance of the original aCompCor approach, we examine whether removing more components than previously suggested, either from WM or CSF, is beneficial. Finally, for the variant of aCompCor that exhibits the best improvement in QC scores, we investigate the additional benefit of: 1) removing nuisance regressors derived from the MPs and physiological recordings, 2) excluding motion-contaminated volumes from the analysis, and 3) performing low-pass filtering before the removal of regressors.

## 2. Methodology

Unless stated otherwise, the preprocessing and analysis described below were performed in Matlab (R2018b; Mathworks, Natick MA).

### 2.1 Human Connectome Project (HCP) Dataset

We used resting-state scans from the HCP S1200 release (Glasser et al., 2016; Van Essen et al., 2013). The HCP dataset includes, among others, resting-state (eyes-open and fixation on a cross-hair) data from healthy young individuals (age range: 22-35 years) acquired on two different days. On each day, two 15-minute scans were collected. We refer to the two scans collected on days 1 and 2 as R1a/R1b and R2a/R2b, respectively. fMRI acquisition was performed with a multiband factor of 8, spatial resolution of 2 mm isotropic voxels, and a repetition time TR of 0.72 s (Glasser et al., 2013).

The minimal preprocessing pipeline for the resting-state HCP dataset is described in Glasser et al. (2013). In brief, the pipeline includes gradient-nonlinearity-induced distortion correction, motion correction, EPI image distortion correction, non-linear registration to MNI space and mild high-pass (2000 s) temporal filtering. The motion parameters are included in the database for further correction of motion artifacts. Apart from the minimally preprocessed data, the HCP provides a cleaned version of the data whereby time series corresponding to ICA components that FIX classified as noisy as well as 24 motion-related regressors (i.e., the 6 MPs estimated during volume realignment along with their temporal derivatives and the squared terms of these 12 time series) were regressed out of the data (Smith et al., 2013a). The cleaned fMRI data are referred to later as FIX-denoised data.

Both minimally-preprocessed and FIX-denoised data are available in volumetric MNI152 and grayordinate space. The grayordinate space combines cortical surface time series and subcortical volume time series from GM, and has more accurate spatial correspondence across subjects than volumetrically aligned data (Glasser et al., 2013; Glasser et al., 2016), particularly when the fMRI data have high spatial resolution such as in HCP.

In the present work, we examined minimally-preprocessed and FIX-denoised data in both volumetric and grayordinate (MSMall registration; Glasser et al., 2016) space from 390 subjects with good quality physiological signals (cardiac and breathing waveforms) in all four scans, as assessed by visual inspection. The cardiac and breathing signals were collected with a photoplethysmograph and respiratory bellow respectively.

### 2.2 Parcellation of the fMRI data

The following three fMRI-based atlases were considered in this study:

a. The Gordon atlas (Gordon et al., 2016): This atlas is composed of 333 cortical regions with 286 parcels belonging to one of twelve large-scale networks and the rest being unassigned. Only the 286 parcels that are assigned to networks were considered in the present study.
b. The Seitzman atlas (Seitzman et al., 2020): This atlas consists of 239 cortical, 34 subcortical and 27 cerebellar volumetric parcels. Among the 300 parcels, 285 parcels are assigned to one of thirteen large-scale networks and only these ones were considered here.
c. The MIST atlas (Urchs et al., 2017): The MIST atlas is available in several resolutions ranging from 7 to 444 parcels. In this study, we considered the MIST_444 parcellation that consists of 444 regions from the whole brain that are assigned to the 7 networks of the MIST_7 parcellation.

All three atlases were defined on resting-state fMRI data and all (MIST) or the majority of (Gordon and Seitzman) their parcels were assigned to large-scale networks. The association of the parcels to networks is required for three of the quality control (QC) metrics described later (i.e., *FCC*, *FD*-*FCC* and *ICCC*). Therefore, as mentioned earlier, parcels that do not belong to a specific network were excluded from the study.

Before the parcellation, in the case of the Seitzman atlas, we performed spatial smoothing on the fMRI data with a Gaussian smoothing kernel of 5 mm full width half maximum (FWHM). Spatial smoothing is commonly done on fMRI data to suppress spatially random noise and enhance the signal-to-noise ratio (SNR). However, when mapping the fMRI data to a parcellation with relatively large parcels such as the parcels in the Gordon and MIST atlases, spatial smoothing is implicitly done. Therefore, we chose to omit spatial smoothing for these two atlases. We performed spatial smoothing before conducting the mapping to the Seitzman atlas because this atlas consists of small spherical ROIs of 8 mm diameter (Seitzman et al., 2020) and, thus, if spatial smoothing is not performed, the parcel time series extracted from these ROIs may suffer from low SNR.

To speed up the preprocessing step, the regression of the nuisance regressors for each pipeline was performed in a parcel-rather than voxel-wise manner. In other words, the minimally-preprocessed and FIX-denoised fMRI data were first mapped to parcel time series by averaging the time series of all voxels or vertices within a parcel and, subsequently, the resulted parcel time series were corrected for noise using the techniques described later. The mapping of the fMRI data to the Gordon parcel space was done using the fMRI data in the grayordinate form, while the mapping to the Seitzman and MIST parcel space was done using the volumetric form of the fMRI data. Finally, to mitigate the influence of spurious low-frequency fluctuations on estimations of correlations, all parcel time series were high-pass filtered at 0.008 Hz, complying with the rule of thumb proposed in Leonardi & Van De Ville (2015) of a high-pass cut-off frequency *f*_*c*_ ≥ 1/(*window length*) = 1/(15 *min*).

### 2.3 Tissue-based regressors

The T1-weighted (T1w) images of each subject are provided in the HCP database in both the native and MNI152 space. To extract the tissue-based regressors used by several pipelines examined here, we initially performed tissue segmentation on the T1w images in the MNI152 space using FAST (FMRIB’s Automated Segmentation Tool) in FSL 5.0.9, generating probabilistic maps for the GM, WM and CSF compartments (Zhang et al., 2001). Subsequently, the GM, WM and CSF binary masks were defined as follows: voxels with a probability above 0.25 of belonging to GM were assigned to GM, while the same was done for WM and CSF for probability values of 0.9 and 0.8 respectively. The choice of the threshold values was made based on visual inspection while overlaying the binary maps on the T1w images. Finally, based on these maps, the global signal was defined as the mean fMRI time series across all voxels within GM. In addition, PCA regressors were obtained separately for voxels in WM and CSF. The GS and PCA regressors were derived from both the minimally-preprocessed and FIX-denoised fMRI data in the volumetric space.

### 2.4 Model-based regressors related to motion and physiological fluctuations

An important goal of this study was to quantify the effect of model-based NCTs with respect to the quality of the fMRI data for atlas-based FC analysis and how they contribute to fMRI denoising when combined with tissue-based regressors. Therefore, for each scan the following four sets of model-based regressors were considered:

a. **Motion parameters (MPs; 24 regressors):** The 6 MPs derived from the volume realignment during the minimal preprocessing, as well as their temporal derivatives, are provided in the HCP database. In addition to the 12 aforementioned time series (12 MPs), we derived their squared terms, yielding in total 24 motion parameters (24 MPs).
b. **Cardiac regressors (6 regressors):** The cardiac regressors were modelled using a 3^rd^ order RETROICOR model (Glover et al., 2000) based on the cardiac signal of each scan. These regressors aimed at accounting for the effect of cardiac pulsatility on the fMRI time series.
c. **Breathing regressors (6 regressors):** The breathing regressors were modelled using a 3^rd^ order RETROICOR model based on the breathing signal of each scan (Glover et al., 2000). These regressors aimed at accounting for the effect of breathing motion.
d. **Systemic low frequency oscillations (SLFOs; 1 regressor):** The SLFOs refer to non-neuronal physiological BOLD fluctuations induced by changes in heart rate and breathing patterns, which were modelled following the framework proposed in our previous work (Kassinopoulos and Mitsis, 2019). The heart rate and respiratory flow extracted from the cardiac and breathing signals of each scan were convolved with scan-specific cardiac and respiratory response functions and the outputs of these convolutions were linearly combined to obtain the corresponding SLFOs. To estimate the scan-specific physiological response functions and determine the linear combination of heart rate- and respiratory flow- related components needed to model the SLFOs, numerical optimization techniques that maximize the fit of the model output (i.e., the SLFOs-related time series) to the GS were employed. The GS was used as a fitting target as it is strongly driven by fluctuations in heart rate and breathing patterns. For more information on this method we refer the reader to Kassinopoulos and Mitsis (2019). The codes for the estimation of SLFOs can be found on https://github.com/mkassinopoulos/PRF_estimation.

Even though we selected subjects with good quality of physiological recordings, it was still important to preprocess both the cardiac and breathing signals to ensure the extraction of accurate heart rate and respiratory flow traces. To this end, we applied temporal filtering and correction for outliers as described in Kassinopoulos and Mitsis (2019). Moreover, as the effect of HR and breathing pattern variations on the fMRI BOLD signal is considered to last about half a minute (Kassinopoulos and Mitsis, 2019) physiological recordings for at least half a minute before the beginning of the fMRI acquisition would be required to account for these effects. However, due to the lack of data at this period, the first 40 image volumes were disregarded from the fMRI data, while the corresponding physiological data were retained so that the SLFOs could be modelled from the beginning of the considered fMRI scan.

### 2.5 Framewise data quality indices

A common index of quality in fMRI data is the framewise displacement (*FD*) introduced by Power et al. (2012). This index is defined as the sum of absolute values of the first derivatives of the 6 motion (realignment) parameters, after converting the rotational parameters to translational displacements on a sphere of radius 50 mm. *FD* is essentially a time series that reflects the extent of motion during the scan. In this work, *FD* was used for six QC metrics that are described in Section 2.6 to quantify the degree of motion artifacts and biases in the preprocessed data. In addition, it was used to identify volumes associated with relatively large *FD* values that are presumably corrupted by motion, and examine the effect of scrubbing, whereby these flagged volumes are discarded before any further analysis (Section 2.8.4).

Another widely used framewise index of data quality is *DVARS* (Derivative of rms VARiance over voxelS) which measures how much the intensity of an fMRI volume varies at each timepoint compared to the previous point (Power et al., 2012). *DVARS* is defined as the spatial root mean square of the voxel time series after they are temporally differentiated. While *DVARS* is obtained from the fMRI time series and is not directly related to head movement, it demonstrates similar trends with *FD* (Power et al., 2012; Suppl. Fig. 10-Suppl. Fig. 11). Similarly to *FD*, *DVARS* was used in this study for two QC metrics related to the effect of motion, as well as to flag volumes corrupted by motion.

### 2.6 Quality control (QC) metrics

Nine QC metrics that are described below were used to evaluate the data quality with respect to the identifiability of large-scale networks and the presence of motion-related artifacts and biases. *FCC*, *FD*-*FCC*, *ICCC*, *FD*-*FDDVARS* and *FD*-*MFC* are novel metrics proposed in the present study whereas the rest of the four metrics were proposed in previous studies. Pearson (full) correlation was calculated between the time series of each pair of parcels resulting in an FC matrix per scan, pipeline and atlas. Apart from *FDDVARS* and *FD*-*FDDVARS*, all other metrics are based on the FC matrix. Note that throughout the text we refer to a pair of parcels as *edge.* Apart from *MICC* and *ICCC*, for all other metrics, only the first of the four scans was considered from each subject. This was done to facilitate the comparison of the scores of those metrics with scores that would have been obtained in conventional datasets which often consist of a single resting-state fMRI scan per subject of 10-15 minutes duration. *MICC* and *ICCC* were calculated using all four scans from each subject, as by definition these two metrics require repeated measures (scans).

#### Functional connectivity contrast (FCC)

A main property of the three parcellations used in this study is that each parcel is assigned to a specific large-scale network, which implicitly suggests that on average a pair of parcels belonging to the same network, also called within-network edge (WNE), should exhibit a higher (towards positive) correlation value compared to a pair of parcels from different networks (between-networks edge; BNE). Based on this property, we assumed that if a pipeline improves the signal-to-noise ratio (SNR) in the data, it should also lead to a larger difference between correlation values corresponding to WNEs and BNEs. To quantify the extent to which WNEs had higher correlation values than BNEs after applying a preprocessing pipeline on the data, we used the metric FC contrast (*FCC*). FCC was defined as the *Z*-statistic of the Wilcoxon rank-sum test related to the null hypothesis that WNEs and BNEs in the FC matrix are samples from continuous distributions with equal medians (note that the real values rather than the absolute values were used in the calculation of the *Z*-statistic). In other words, higher values of *FCC* correspond to higher (towards positive) correlation values for WNEs as compared to BNEs. Furthermore, for the optimal pipeline found in this work, we quantified the identifiability of each of the networks separately using the same metric but considering only WNEs belonging to the network of interest rather than WNEs from all networks when comparing WNEs with BNEs.

#### FD-FCC

While it is desired to improve the *FCC* score for the data of each scan, at the same time it is desired that low-motion and high-motion scans demonstrate similar *FCC* scores. Therefore, *FD*-*FCC* was defined as the correlation between the mean *FD* and *FCC* across scans and was used in this work to assess potential biases due to different levels of motion between scans.

#### Median of Intraclass correlation values (MICC)

Intraclass correlation (*ICC*) is a widely used metric in statistics to assess how reproducible measurements of the same quantity are across different observers or instruments (Shrout and Fleiss, 1979). Similar to previous studies evaluating the performance of preprocessing strategies in fMRI, we have used *ICC* to assess test-retest reliability across the four sessions of each subject in whole-brain FC estimates (Birn et al., 2014; Parkes et al., 2018; Shirer et al., 2015). For a pair of parcels *i* and *j*, *ICC*_*i*,*j*_ was defined as

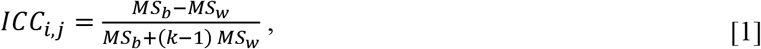

where *k* is the number of scans per subject (4), *MS*_*b*_ is the between-subject mean square of correlation values between parcels *i* and *j*, and *MS*_*w*_ is the within-subject mean square of correlation values for the same pair of parcels. The *MICC* score assigned to each pipeline for the assessment of data quality was defined as the median of *ICC*_*i*,*j*_ across all edges.

#### ICC contrast (ICCC)

A common finding from previous studies is that *MICC* values tend to drop when a preprocessing pipeline is applied (Birn et al., 2014; Parkes et al., 2018; Shirer et al., 2015). This finding suggests that artifacts in fMRI data have high subject-specificity and, thus, when the artifacts are reduced after the preprocessing, *MICC* decreases as well. In addition, Birn et al. (2014) have observed that the 200 most significant connections exhibited higher mean *ICC* after preprocessing as compared to the mean *ICC* across all (16100) connections. Similarly, we also observed that WNEs that exhibited higher connectivity strength than BNEs also exhibited higher ICC values. Based on this observation, in the present study we propose a new metric termed *ICC* contrast (*ICCC*), which quantifies the difference in *ICC*_*i*,*j*_ values between WNEs and BNEs. The assumption behind *ICCC* is that only WNEs should demonstrate high subject specificity. Similarly to *FCC*, *ICCC* was defined as the *Z*-statistic of the Wilcoxon rank-sum test related to the null hypothesis that WNEs and BNEs in the *ICC* matrix are samples from continuous distributions with equal medians. In other words, a high value of *ICCC* suggests that WNEs have higher *ICC*_*i*,*j*_ values compared to BNEs.

#### FDDVARS

To assess the presence of motion artifacts in the parcel time series after preprocessing, we calculated the Pearson correlation coefficient between *FD* and *DVARS* (Muschelli et al., 2014). Even though *FD* was estimated only once based on the motion (realignment) parameters, the framewise data quality index *DVARS* was estimated for each pipeline separately using the parcel (instead of voxel) time series after the preprocessing, as described in Section 2.5 (note that the parcel-wise *DVARS* considered here was computed across all parcels over the whole-brain and should not be confused with the parcel-wise *DVARS* considered in Muschelli et al. (2014) that was computed across voxels within a parcel). Even though the parcel-wise and voxel-wise *DVARS* values were similar (mean correlation of 0.56±0.21), they were not identical (Suppl. Fig. 10-Suppl. Fig. 11). The parcel time series, defined as the average time series of voxels assigned to each parcel, exhibited lower levels of noise compared to voxel time series and, as a consequence, the parcel *FDDVARS* values were also found to be lower (Suppl. Fig. 10-Suppl. Fig. 11). And as it is common practice to conduct the FC analysis at the parcel-level, we performed the assessment of data quality using the parcel *FDDVARS* values. The *FDDVARS* score assigned to each pipeline was obtained by initially calculating the correlation value between *FD* and *DVARS* for each scan and then averaging the correlations across all scans.

#### FD-FDDVARS

While the *FDDVARS* score reflects the extent to which motion artifacts are present in the data of a given scan, a low mean value of *FDDVARS* within a group of scans does not necessarily imply that the motion-related biases across scans in the FC estimates are also low. High-motion scans are contaminated by more severe motion artifacts compared to low-motion scans, which has been shown to systematically bias the estimated FC matrices (Power et al., 2015). Even though a preprocessing strategy may reduce the motion artifacts in both high- and low-motion scans, if in the preprocessed data there are still differences in the levels of motion artifacts between the two groups, this would result in a systematic bias for the FC estimates of these groups. To assess the presence of motion-related biases, we used the QC metric *FD*-*FDDVARS*, which was defined as the correlation between the mean *FD* and *FDDVARS* across scans.

#### *FDFC*_median_

For a pair of parcels *i* and *j*, *FDFC*_*i*,*j*_ was defined as the correlation between the Pearson correlation coefficient of this pair (i.e., *FC*_*i*,*j*_) and the mean *FD* across scans. To assess the quality of data with respect to motion-related biases in FC, each pipeline was assigned an *FDFC*_median_ score that corresponded to the median absolute *FDFC*_*i*,*j*_ across all edges (Parkes et al., 2018; Power et al., 2012).

#### *FDFC*_dist_

Earlier studies on the effect of motion in fMRI have shown that, in the case of raw data, the correlation values for short-distance edges are inflated to a greater extent due to motion compared to long-distance edges (Power et al., 2012; Satterthwaite et al., 2013). An interpretation for this distance-dependent effect is that when motion occurs, nearby voxels are typically affected by similar displacements and, thus, present similar spin-history artifacts as well. And even though distant voxels may also exhibit similar displacements that can lead to widespread artifacts, these artifacts are not as pronounced as in the case of nearby voxels (Power et al., 2015). As these distance-dependent motion artifacts are generally considered undesirable, the quality control metric *FDFC*_dist_ is commonly used to quantify the degree of this dependence and assess the capability of a preprocessing strategy to mitigate it (Ciric et al., 2018, 2017; Muschelli et al., 2014; Parkes et al., 2018; Power et al., 2015, 2012; Satterthwaite et al., 2019, 2012). *FDFC*_dist_ is defined as the correlation between the *FDFC*_*i*,*j*_ value, as defined for the previous QC metric (*FDFC*_median_) and the Euclidean distance between parcels *i* and *j*, across all edges (Ciric et al., 2017; Parkes et al., 2018).

#### FD – Mean FC (FD-MFC)

The metric *FDFC*_median_ inherently assumes that the Pearson correlation of an edge is affected by motion in the same way across subjects. However, considering that differences in brain anatomy across subjects exist (Bijsterbosch et al., 2018), we can not necessarily assume that motion affects edge-wise FC estimates in the exact same way across subjects. Therefore, to assess the effect of motion on FC estimates, without the aforementioned assumption, we propose the *FD*-*MFC* metric which is defined as follows: First, the mean of all Pearson correlations in the FC matrix (*MFC*) is estimated for each scan separately (considering only unique pairs of parcels) and, subsequently, the correlation between the mean *FD* and *MFC* across all scans (subjects) is obtained.

### 2.7 Normalization of QC metrics

The nine QC metrics described in Session 2.6 can be categorized into signal-related and motion-related metrics. The signal-related metrics (*FCC*, *MICC* and *ICCC*) reflect the SNR of the data and are expected to yield low scores for data that have not been preprocessed, as high levels of artifacts are likely to obscure the signal of interest. They are also expected to yield low scores for data to which a very aggressive pipeline is applied and the signal of interest is lost. On the other hand, relatively high scores of signal-related metrics would be expected for data whereby a good pipeline is applied and artifacts are reduced while the signal of interest is preserved. Motion-related metrics are expected to yield high absolute scores for data that have not been preprocessed, indicating the presence of motion artifacts or biases, whereas for gradually more aggressive pipelines, these scores are expected to approach zero, reflecting the reduction of motion artifacts and biases.

As the goal of a preprocessing strategy is to remove artifacts while also preserving the signal of interest, ideally a pipeline that yields high scores in signal-related QC metrics and low scores in motion-related metrics would be preferred. However, due to the fact that each QC metric is based on different assumptions and some metrics are based on Pearson correlation while other ones are based on the Wilcoxon rank-sum test, each metric illustrates a different trend across pipelines and yields a different range of scores (see for example Fig. 2) making the selection of the optimal pipeline difficult. Therefore, to overcome this potential drawback, we followed the following steps:

1. First, we randomly split the 390 subjects to 10 groups of 39 subjects, ensuring that the groups were characterized by similar distributions of mean *FD* values.
2. Then, the QC scores were estimated for each of the 10 groups separately. Apart from *MICC* and *ICCC*, for all other metrics, only the first of the four scans were considered from each subject to avoid estimating correlations with repeated measures. *MICC* and *ICCC* were calculated using all four scans from each subject, as by definition these two *ICC*-based metrics require repeated measures (scans). *FCC* and *FDDVARS* that are calculated on a scan basis, rather than within a group of scans, were averaged across subjects within each group. In this way, the quality of the data for a given atlas, pipeline and group of subjects was assigned with one score for each of the nine QC metrics.
3. Subsequently, all motion-related metrics were normalized to 1 – *abs*(*x*), where x is the score of each metric, so that, similarly to signal-related metrics, a high positive score is assigned to good quality data.
4. In the next step, the scores were expressed as *Z*-scores based on the relation 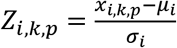 where *x*_*i*,*k*,*p*_ and *z*_*i*,*k*,*p*_ are respectively the original and *Z*-score values obtained for QC *i*, group of subjects *k* and pipeline *p* and, *μ*_*i*_ and *σ*_*i*_ are respectively the mean and standard deviations of the scores from all groups of subjects related to the QC *i* obtained from the raw data. For example, considering the case where the *FCC* scores obtained from the raw data for the ten groups of subjects have a mean and standard deviation of 47.3 and 1.6 respectively, if the *FCC* score for a given pipeline (e.g. FIX) and group of subjects is 56.0, the corresponding *Z*-score for that particular pipeline and group of subjects would be 5.4 [(56.0-47.3)/1.6=5.4].
5. Subsequently, the *Z*-scores of *FCC* and *ICCC* were averaged to yield the summarized score *QC*_signal_ and the *Z*-scores of the 6 motion-related metrics *FD*-*FCC*, *FDDVARS*, *FD*-*FDDVARS*, *FDFC*_median_, *FDFC*_dist_ and *FD*-*MFC* were averaged to yield the summarized score *QC*_motion_. The *MICC* was not included in the estimation of the *QC*_signal_ score as it was proven to reflect subject-specificity due to noise rather than signal of interest.
6. Finally, the two latter summarized scores, *QC*_signal_ and *QC*_motion_, were averaged to obtain the score for the combined quality control metric *CQC*.

The normalization described here allowed us to express the scores of the QC metrics as *Z*-scores that reflect the relative improvement in standard deviations with respect to the raw data. High *Z*-scores can be interpreted as the associated QC metric exhibiting high sensitivity. If the QC metric for a given pipeline exhibits a high *Z*-score, it is very likely that in a new dataset the score for the same QC metric will be better when the data are preprocessed with the same pipeline compared to the raw data. After the normalization of the QC metrics, all metrics were summarized into two indices, the *QC*_signal_ and the *QC*_motion_, which, in turn, were averaged to obtain the final *CQC* score. While we present the results for *QC*_signal_ and *QC*_motion_ apart from *CQC*, the choice for the *optimal* pipeline is based on the third one.

### 2.8 Noise correction techniques (NCTs)

We assessed the performance of a large number of preprocessing pipelines using eight QC metrics that quantify the improvement in network identifiability and reduction of motion-related artifacts and biases. The pipelines consisted of commonly used preprocessing strategies, namely scrubbing, temporal low-pass filtering and removal of nuisance regressors through linear regression. To better understand the effect of each of the aforementioned strategies, five different groups of pipelines were examined that are described in the following sections. The QC metrics used to evaluate the performance of each pipeline are described in Section 2.6.

#### 2.8.1 Optimizing aCompCor

In aCompCor, PCA regressors are obtained from the WM and CSF tissues and the first five components ordered by the variance explained in the WM and CSF voxel time series are used as nuisance regressors. This practice implicitly suggests that the PCA regressors that explain most of the variance in WM and CSF are also the ones with the stronger association to model-based nuisance regressors. To examine whether the latter is the case, we estimated the fraction of variance in WM and CSF PCA regressors explained by each of the following sets of regressors: 1) 24 MPs, 2) breathing regressors, 3) cardiac regressors, 4) SLFOs, and 5) all the aforementioned regressors combined. In addition, we estimated the fraction of variance in the model-based regressors explained by an increasing number of WM or CSF regressors (between 1 and 100). The estimated explained variances corresponding to each model-based regressor were averaged across regressors associated with the same source of noise.

After confirming the hypothesis stated above, a main objective of this study was to examine the performance of WM and CSF denoising independently and determine the number of regressors that improves the quality of the fMRI data the most. To this end, for both noise ROIs, we considered the removal of the most significant PCA regressors with or without including the GS as an additional nuisance regressor for a varying number of PCA regressors between 1 and 600 (to keep the computational time low we varied the number using a logarithmic base of ten as follows: 1, 2, … ,10, 20, … , 100, 200, … , 600). Note that each scan consisted of 1160 fMRI volumes, therefore 600 components correspond to about half of the available PCA regressors. Regarding the tissue-based regressors used in aCompCor, we refer to a set of regressors from WM and CSF as 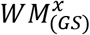 and 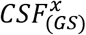, respectively, where *x* indicates the number of PCA regressors considered from each of the two tissue compartments and the presence of the *GS* as superscript denotes the inclusion of the GS in the set of nuisance regressors. For example, the set of regressors 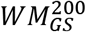 refers to the set consisting of the *GS* and the first 200 PCA regressors from WM. Note that the set 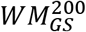 demonstrated the highest improvement in QC scores and, thus, the subsequent analyses in this work investigate the possibility of further improvement using additional strategies in the preprocessing along with the regression of this set.

#### 2.8.2 Evaluation of data-driven NCTs

Typically, fMRI studies consider only data-driven regressors for the preprocessing of the data that can be a combination of motion, tissue-based regressors or whole-brain component regressors (e.g., FIX). However, it is still not clear which preprocessing strategy is the best for whole-brain FC as the number and type of regressors included in the preprocessing stage vary across studies. In the present study, a selection of pipelines used in the literature were evaluated using the QC metrics described in Section 2.6 to allow a comparison between them (Table 1). However, as the focus of this analysis was to examine the effect of the regressors *per se* rather than the entire pipeline, steps such as scrubbing (i.e., removal of motion-contaminated volumes) or temporal low-pass filtering were omitted. In addition, several pipelines were considered in this analysis that consisted of a small number of regressors (e.g. pipelines 1-5), even though, typically, more aggressive pipelines are found in the literature. These pipelines were considered in order to better understand possible differences in QC scores yielded by more aggressive pipelines (e.g. pipelines 7 and 8).

**Table 1.** Preprocessing pipelines based on data-driven approaches

| Pipeline | Sets of nuisance regressors considered in the pipeline | (Related to) pipeline used in: |
| --- | --- | --- |
| 1 | 6 motion parameters (6 MPs) | — |
| 2 | 12 motion parameters (12 MPs; i.e., 6 MPs plus their derivatives) | — |
| 3 | 24 motion parameters (24 MPs; i.e., 12 MPs plus their squared terms) | — |
| 4 | Global signal (GS) | — |
| 5 | 12 MPs, GS | — |
| 6 | $WM^5$, $CSF^5$ (i.e., first 5 PCA regressors from white matter (WM) and first 5 PCA regressors from cerebrospinal fluid (CSF)) | Behzadi et al., 2007 |
| 7 | 12 MPs, mean WM time-series ($WM_{mean}$) and mean CSF time-series ($CSF_{mean}$) | Urchs et al., 2017 |
| 8 | 12 MPs, $WM_{mean}$, $CSF_{mean}$, GS | Finn et al., 2015 |
| 9 | 24 MPs, [GS, $WM_{mean}$, $CSF_{mean}$, plus their derivatives] | Laumann et al., 2017 |
| 10 | 24 MPs, [GS, $WM_{mean}$, $CSF_{mean}$, plus their derivatives and the squared terms of the 6 aforementioned time-series] | Ciric et al., 2017; Xia et al., 2018 |
| 11 | *FIX* (i.e., the *FIX*-denoised data as provided from HCP) | Bijsterbosch et al., 2017; Smith et al., 2015; Zhang et al., 2018 |
| 12 | $FIX_{GS}$ (i.e., *FIX* followed by GS regression) | Burgess et al., 2016 |
| 13 | *FIX*, GS, $WM^5$, $CSF^5$ | Siegel et al., 2017 |
| 14 | PCA regressors needed to explain 50% of variance in WM and CSF | Muschelli et al., 2014 |
| 15 | GS, WM regressors needed to explain 30% of variance | — |
| 16 | GS, WM regressors needed to explain 35% of variance | — |
| 17 | GS, WM regressors needed to explain 40% of variance | — |
| 18 | GS, WM regressors needed to explain 45% of variance | — |
| 19 | GS, WM regressors needed to explain 50% of variance | — |
| 20 | $WM_{GS}^{200}$ (i.e., the GS and the first 200 PCA regressors from WM) | — |

Regarding the pipelines that involve FIX (i.e., pipelines 11-13), even though the HCP database provides the results from MELODIC-ICA and, thus, we could have removed noisy ICA components and further nuisance regressors from the minimally-preprocessed fMRI data in one step, we chose to remove the additional nuisance regressors from the FIX-denoised data found in the HCP database to be consistent with the approach taken in previous studies (Burgess et al., 2016; Siegel et al., 2017). For pipelines 12 and 13, we used the GS and WM/CSF regressors derived from the FIX-denoised data. Note that, as mentioned earlier, the FIX denoising performed in HCP included the removal of the 24 MPs.

#### 2.8.3 Evaluation of model-based (motion and physiological) NCTs

Even though the motion parameters are indirectly derived from the data through the process of volume realignment, they do not purely correspond to motion-induced fMRI artifacts, but rather rigid-body displacements (Patriat et al., 2017). Therefore, treating them as nuisance regressors during preprocessing inherently imposes some assumptions about the effect of motion on the fMRI time series, which may not be valid. Similarly, the efficiency of physiological regressors that are obtained from concurrent physiological recordings (e.g. cardiac and breathing signals) depends on the validity of the employed physiological response function models, as well as the quality of the recordings. Thus, an important question that needs to be addressed is whether the aforementioned model-based regressors contribute to the denoising of the fMRI data, and particularly when combined with tissue-based regressors that do not have the limitations of the model-based approaches. To this end, using the QC metrics, we evaluated 64 combinations of pipelines that employ sets of model-based and tissue-based regressors. Specifically, we considered as model-based regressors the 24 MPs, the cardiac and breathing regressors and the SLFOs regressor, while as tissue-based regressors we considered the GS and 200 PCA regressors from WM (*WM*_200_).

#### 2.8.4 Scrubbing

In the analyses preceding scrubbing, it was found that the set of nuisance regressors 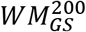 yielded the highest QC scores. Therefore, the next question that we aimed to address is whether scrubbing can provide any further improvement in the QC scores for this specific set of regressors and at what threshold. This analysis was done both using the *FD* and the *DVARS* to determine the motion-contaminated volumes that would be discarded. In the case of *FD*, we repeated the analysis for the values of threshold *FD*_thr_ 0.15, 0.20, 0.25, 0.30, 0.50, 0.80 and 1.00 mm, and in the case of *DVARS* we considered the values of threshold *DVARS*_thr_ 0.5, 1, 1.5, 2, 5, 10 and 20 median absolute deviations (MAD).

#### 2.8.5 Low-pass filtering (LPF)

We also investigated whether low-pass filtering the data and removing nuisance regressors would yield higher QC scores compared to no filtering. Low-pass filtering, however, apart from removing high-frequency noise, it also leads to a substantially decreased number of degrees of freedom (Bright et al., 2017). The degrees of freedom *DOF* can be roughly estimated by the relation: 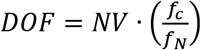 where *NV* is the number of volumes, *f*_*c*_ is the cut-off frequency and *f*_*N*_ the Nyquist frequency. For instance, in our dataset, low-pass filtering at 0.08 Hz (*NV* = 1160, 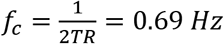 = 0.69 *Hz*) reduces the degrees of freedom from 1160 to ~134. This property suggests that removing a large set of nuisance regressors (e.g. 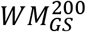), in addition to applying a strong low-pass filter (e.g. a low-pass filter with 0.08 Hz cut-off frequency), can remove all the degrees of freedom available in the data. Given this, we examined the effect of low-pass filtering for both mild and aggressive variants of WM denoising, considering only cut-off frequencies that yield data with a minimum of 30 degrees of freedom. Specifically, we examined the sets 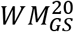 and 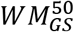 with low-pass filtering at 0.05, 0.08, 0.2, 0.3, 0.4, 0.5 and 0.6 Hz, the set 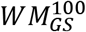 with values of cut-off frequency between 0.08 Hz and 0.6 Hz, as well as the set 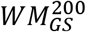 with values between 0.2 and 0.6 Hz.

## 3. Results

Here we present results mainly for the Gordon atlas since the results between the three examined atlases did not yield significant differences. The results obtained using the Seitzman and MIST atlases can be found in the Supplementary Material.

### 3.1 Optimizing aCompCor

Fig. 1 shows the estimated fraction of variance in WM and CSF PCA regressors explained by various sets of model-based regressors (left panel), as well as the estimated fraction of variance explained in the sets of model-based regressors by an increasing number of WM and CSF regressors (right panel). The WM and CSF regressors were ordered with respect to decreasing tissue signal variance explained (see Suppl. Fig. 1 for the variance explained in the WM and CSF compartments by the corresponding WM and CSF regressors). In the left panel, it can be seen that the model-based regressors explained larger fraction of variance in the first few WM and CSF regressors than in subsequent regressors. Moreover, the high-motion scans demonstrated different trends compared to low-motion scans. For example, in the left panel we can see that high-motion scans exhibited stronger association between the first PCA regressors and the 24 MPs, while low-motion scans exhibited stronger association between the first PCA regressors and regressors related to cardiac pulsatility. Looking at WM vs CSF regressors, we observed several slight differences such as that the first few WM regressors explained better the 24 MPs compared to the CSF regressors, whereas the opposite was observed when looking at the cardiac pulsatility. However, when considering a large number of regressors (e.g. 100) both WM and CSF regressors explained a significant fraction of variance for all four sets of model-based regressors, with mean correlation values above 0.5. This suggests that both WM and CSF regressors can account (to some extent) for BOLD fluctuations due to head and breathing motion as well as cardiac pulsatility and SLFOs.

**Fig. 1.**
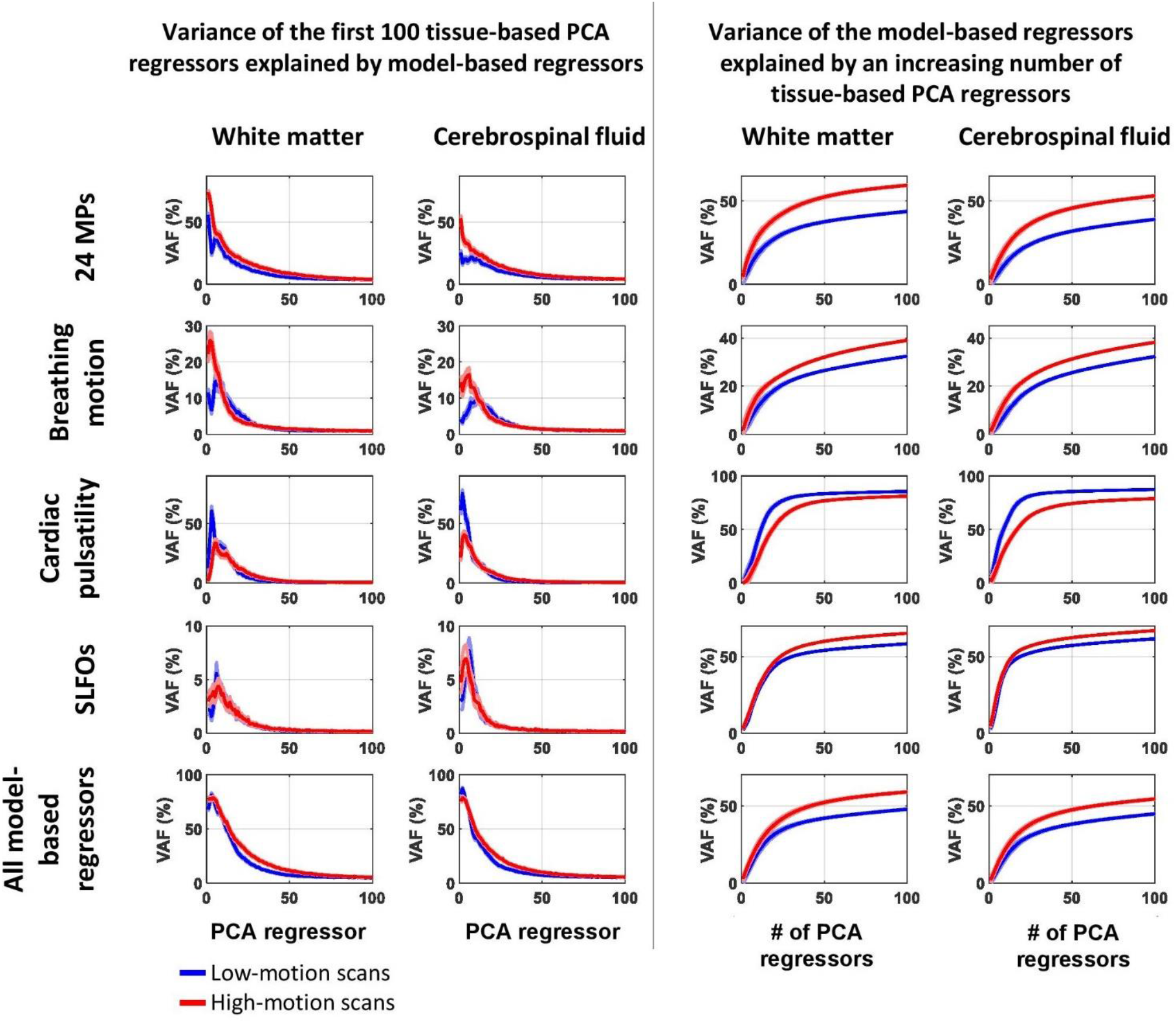
Relation between model-based regressors and PCA regressors obtained from WM and CSF. Left panel (first two columns): Estimated variance explained in each of the first 100 WM and CSF PCA regressors by the set of model-based regressors stated on the left of each row (for instance, it shows the variance in the 50th principal component of WM explained by the 24 MPs). Right panel (last two columns): Mean estimated variance explained in the regressors stated on the left of each row by an increasing number of WM or CSF regressors (for instance, it shows the variance explained on average in the 24 MPs by the first 50 principal components of WM). To examine the dependence of the curves on the degree of motion in each scan, two groups of scans were considered, referred to as low- and high-motion scans, corresponding to the lower and upper quartile of the distribution of mean *FD* values respectively. The blue and red curves correspond to the variance accounted for (VAF) averaged across low-motion and high-motion scans, respectively, whereas the shaded areas denote the standard error. For all four sources of noise, we observe that the first few PCA regressors demonstrated stronger association to the model-based regressors compared to components found later in the order, justifying the practice of using the most significant PCA components in aCompCor.

Different trends were observed between the nine QC metrics when varying the number of components (Fig. 2, Suppl. Fig. 2). Similar to previous studies, the signal-related metric median intraclass correlation (*MICC*) yielded high scores in the raw data whereas when preprocessing was performed, the *MICC* was decreasing as more WM or CSF regressors were removed (Fig. 2c; Birn et al., 2014; Parkes et al., 2018). This trend is possibly due to that noise in fMRI is characterized by high subject specificity and, hence, removing the noise when using more aggressive pipelines leads to reduction in subject specificity (Birn et al., 2014; Parkes et al., 2018). As the *MICC* metric did not seem to reflect the preservation of signal in the data, we excluded it from the rest of the analysis.

**Fig. 2.**
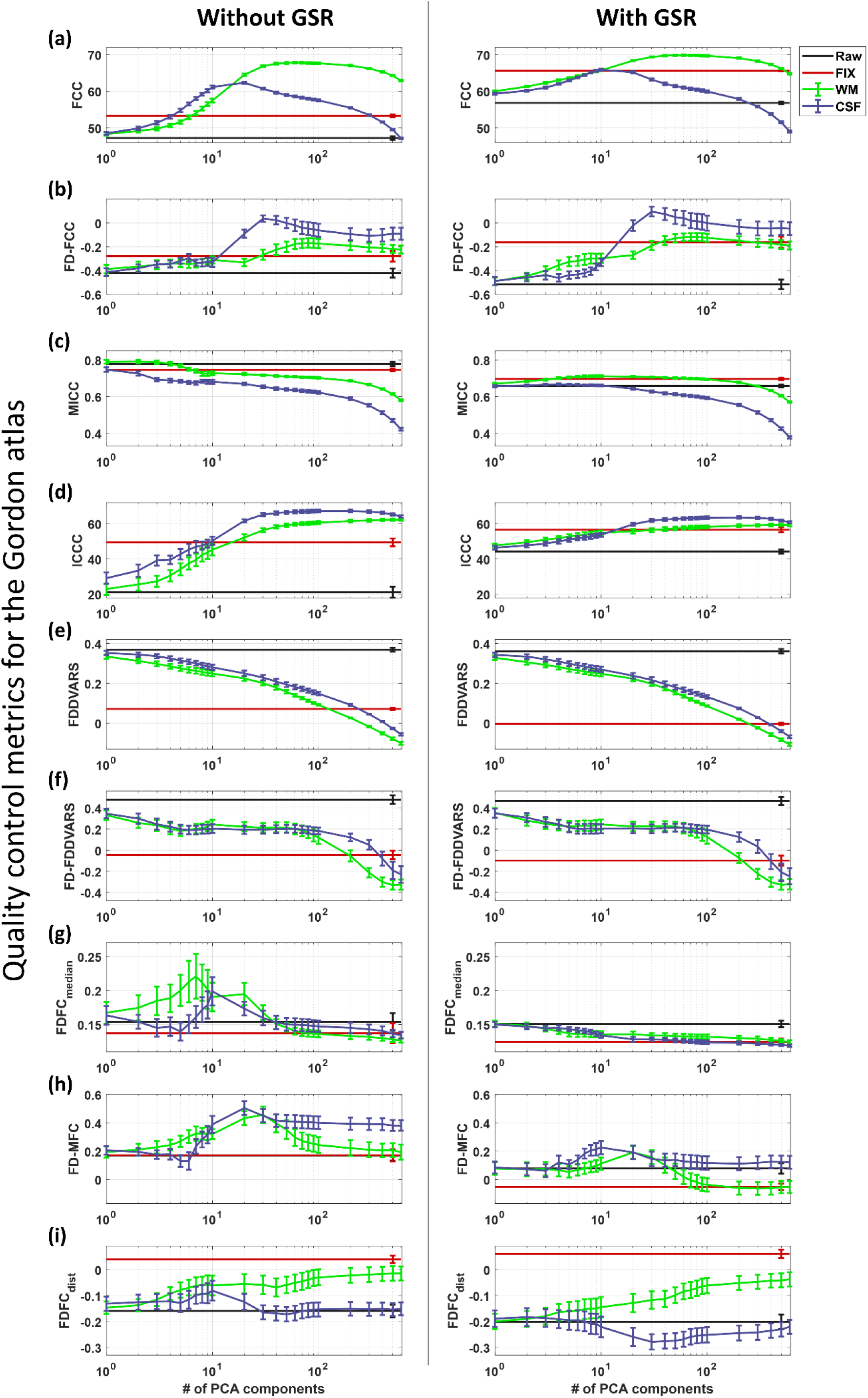
Quality control (QC) scores obtained by aCompCor using the data in the Gordon parcel space. The green and purple curves show the dependence of the QC scores on the number of WM and CSF PCA regressors that were removed from the data respectively. The black and red lines indicate the QC scores for the raw (i.e., minimally-preprocessed data) and FIX-denoised data. The middle points in the curves and lines correspond to the QC scores averaged across the 10 groups of subjects and the error bars indicate the standard error across the 10 groups. To ease visualization, the error bars for the raw and FIX-denoised data are shown in the column corresponding to 500 components. The two columns correspond to the global signal being regressed out (right) or not (left) whereas the rows (a)-(i) correspond to the nine QC metrics described in Session 2.6.

Fig. 3 shows the metric *QC*_signal_ which was defined as the mean *Z*-score of the two signal-related metrics, *FCC* and *ICCC*, as well as the metric *QC*_motion_, defined as the mean *Z*-score of the six motion-related metrics, *FD*-*FCC*, *FDDVARS*, *FD*-*FDDVARS*, *FDFC*_median_, *FDFC*_dist_ and *FD*-*MFC*. Fig. 3 also shows the scores for the combined QC metric CQC, defined as the average score between *QC*_signal_ and *QC*_motion_. Note that due to their definition, *QC*_signal_ reflects the enhancement of SNR in the data, whereas *QC*_motion_ reflects the reduction in motion artifacts and biases. Even though WM and CSF denoising yielded similar performance in terms of mitigating motion effects, WM denoising achieved a considerably higher SNR compared to CSF denoising (Fig. 3). Furthermore, including the GS to the nuisance regressors significantly improved the scores for both *QC*_signal_ and *QC*_motion_, particularly for low numbers of PCA components. Due to these observations, the subsequent results and discussion are focused on the performance of WM denoising, and, unless explicitly stated otherwise, it is assumed that the GS is also included in the set of regressors.

**Fig. 3.**
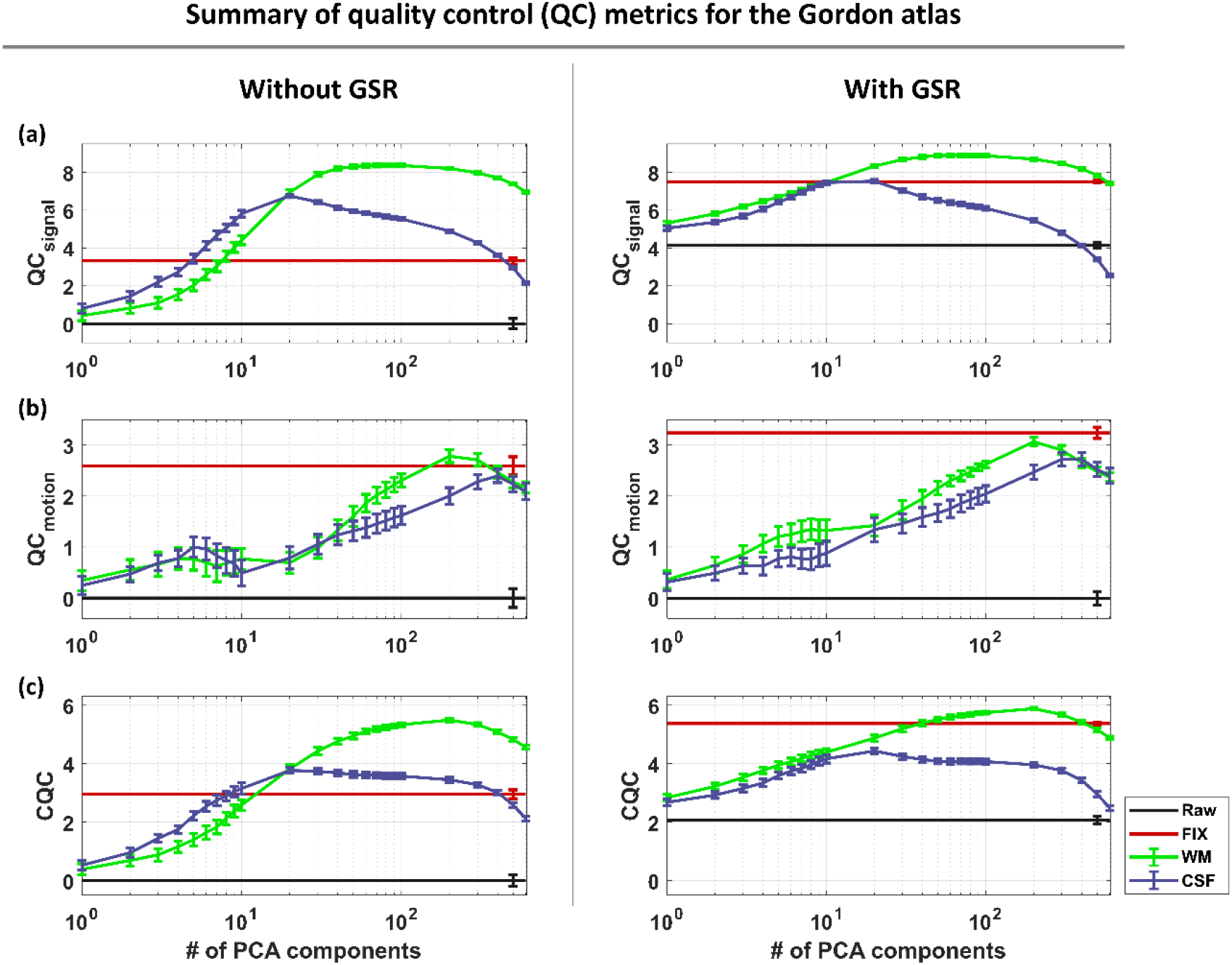
Summarized quality control (QC) scores obtained by aCompCor using the data in the Gordon parcel space. The *Z*-scores of the two signal-related metrics *FCC* and *ICCC*, and six motor-related metrics *FD*-*FCC*, *FDDVARS*, *FD*-*FDDVARS*, *FDFC*_median_, *FDFC*_dist_ and *FD*-*MFC* were averaged to yield the summarized scores *QC*_signal_ (a) and *QC*_motion_ (b), respectively. Subsequently, the two latter summarized scores were averaged to obtain the combined QC metric (*CQC*). We observe that about 50 to 100 PCA regressors from WM were needed in order to achieve high *QC*_signal_ scores, while 200 components from WM yielded the highest score in *QC*_motion_. Including the GS in the set of regressors led to slightly higher scores for both summarized metrics. With respect to the CQC metric, the set of regressors 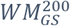 yielded the highest score (5.9) with the *FIX*_*GS*_ demonstrating the second highest score (5.4). While CSF denoising yielded comparable scores to WM denoising with respect to reduction of motion artifacts and biases (*QC*_motion_), it also led to loss of signal of interest based on the low scores of *QC*_signal_.

Overall, we observed that *QC*_signal_ was high for the sets of regressors 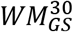 to 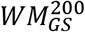 with a maximum score of 8.9 for 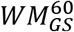 (Fig. 3). In contrast, *QC*_motion_ illustrated a sharp peak for the more aggressive set of regressors 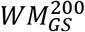 and, as a consequence, the optimal set of regressors according to CQC was the latter one (i.e., 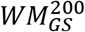). Note that the fMRI scans considered in this study consisted of 1160 volumes, therefore the 200 WM regressors used in the preprocessing correspond to ~17% of the total WM regressors. To allow generalization of our results to conventional datasets that typically have a longer TR between 2-4 s, we repeated the analysis for the case of the Gordon atlas after downsampling the fMRI data by a factor of four, resulting in an effective TR of 2.88 s. For this subset of fMRI data that consisted of 290 volumes per scan, the best improvement, as assessed with the summarized metric CQC, was observed with the set of regressors 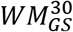 (Suppl. Fig. 7).

The analysis of optimizing aCompCor was repeated in the Seitzman and MIST parcel space and yielded similar trends for a varying number of PCA regressors (Suppl. Fig. 3-Suppl. Fig. 6). Similar to the data in the Gordon space, the set 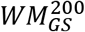 was found to be the best choice for the data in the Seitzman atlas space, whereas the set 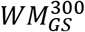 seemed to perform slightly better for the data in the MIST atlas space. In the following analyses, for both three atlases, we considered 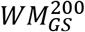 when comparing the performance with other preprocessing strategies (e.g. scrubbing and low-pass filtering).

#### Signal-related QC metrics

##### Functional connectivity contrast (FCC)

The metric *FCC* proposed in this work for assessing the identifiability of large-scale networks exhibited unimodal curves for both WM and CSF, both with and without GSR (Fig. 2a). However, WM denoising achieved higher scores with a maximum score of 69.9 for 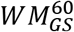. Fig. 4 shows the FC matrix for the raw (minimally-preprocessed) data and for data that have been preprocessed with different pipelines for a scan that demonstrated considerable improvement in identifiability of the networks when regressing out the set 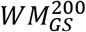. It is evident that the raw data were very noisy, preventing the identification of the networks (*FCC*=23.0) but when denoising was performed with 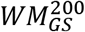, all 12 networks were clearly identified (*FCC*=69.4). Interestingly, when GSR was applied without any other nuisance regressor or NCT to the raw data, it did not have a strong effect on the contrast but when it was applied after FIX denoising it led to a significant increase of *FCC* score from 40.1 to 65.4. Overall, GSR improved the FCC score for both FIX and WM denoising (Fig. 2a).

**Fig. 4.**
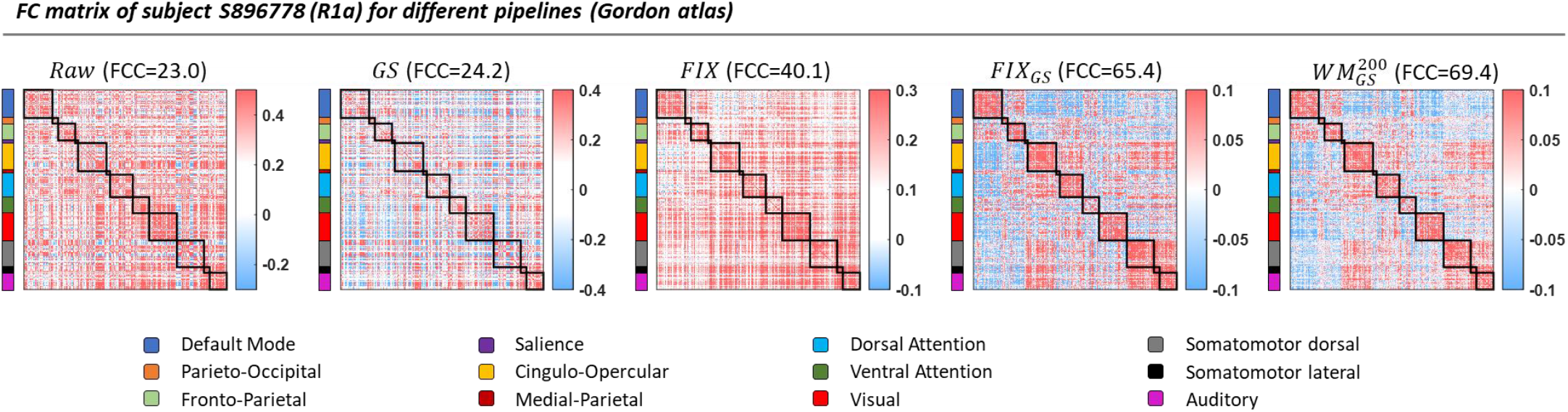
FC matrix of subject S896778 (R1a) for different pipelines (Gordon atlas). While the networks could not be distinguished by visually inspecting the FC matrix of the raw data, they were easily identified after regressing out the set 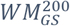 or after FIX denoising, especially when FIX was combined with GSR. Similar conclusions were drawn based on the *FCC* metric, which quantifies the identifiability of the networks (reported on the top of each matrix).

Fig. 5 shows the FC matrices averaged across all 1560 scans (group-level FC matrices) obtained from raw and four preprocessed fMRI datasets (i.e., data preprocessed with different pipelines). The FCC estimated from the group-level FC matrices were substantially higher compared to the *FCC* estimated on a scan basis for the same pipelines (Fig. 2a). Note that for the raw data, the *FCC* score that was estimated first on a scan-basis and, then, averaged across all scans was 47.3 (Fig. 2a), whereas the *FCC* score estimated from the group-level FC matrix (i.e. the FC matrix was first averaged across all scans) had a higher value of 67.2 (Fig. 5). In addition, the *FCC* score obtained from the group-level FC matrix (67.2) was at similar levels with the highest *FCC* score achieved on a scan-specific basis across all pipelines (i.e., when preprocessed with 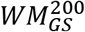).

**Fig. 5.**
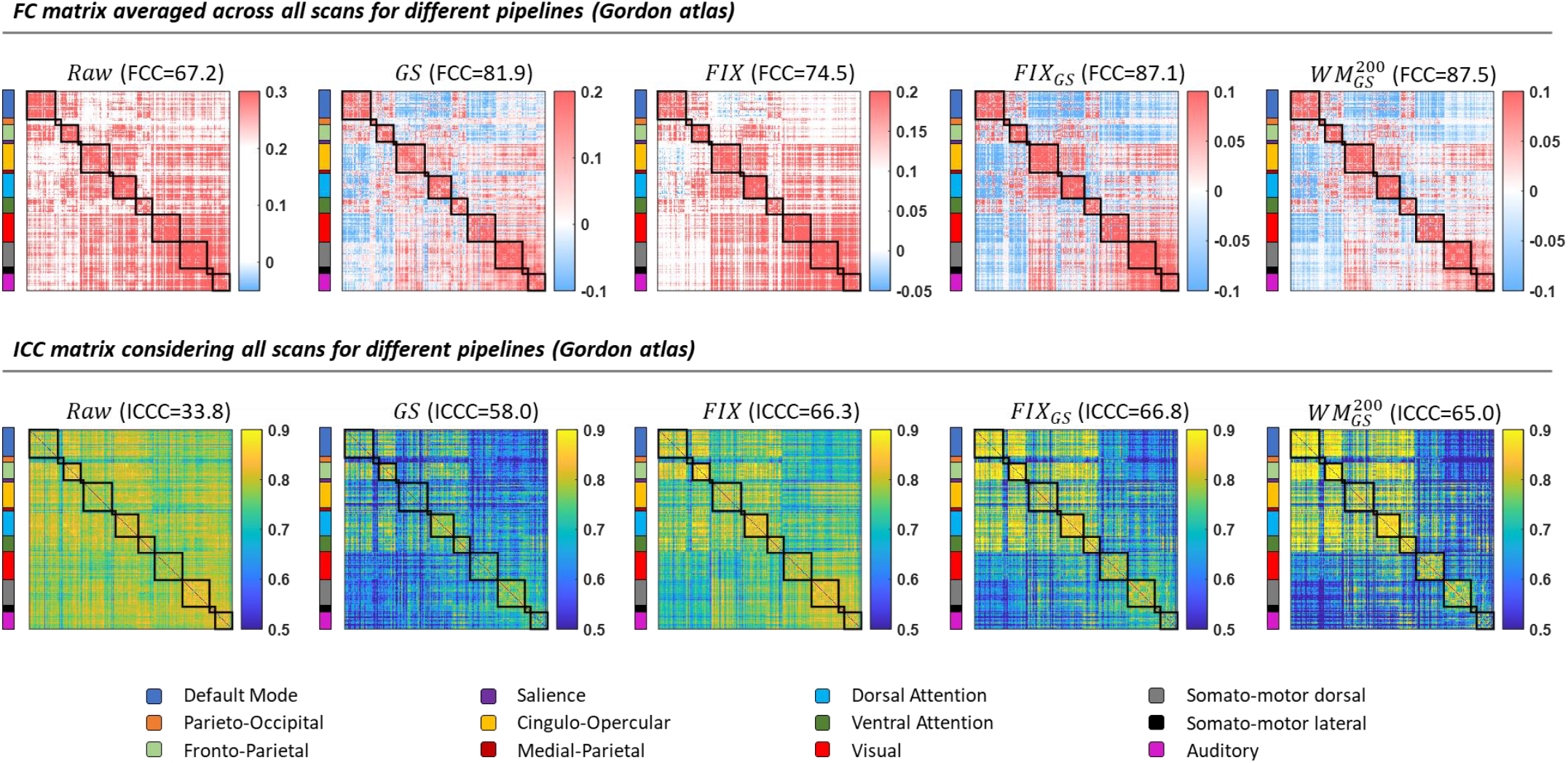
FC (top) and ICC (bottom) matrices considering all scans for different pipelines obtained from the data in the Gordon parcel space. Averaging the FC matrices across all 1560 scans improved the identifiability of the networks considerably for both the raw and preprocessed data. As a consequence, the associated *FCC* scores reported on the top of each matrix were higher than the scores presented in Fig. 2a, which were obtained on a scan-specific basis and subsequently averaged within groups of 39 subjects. Similarly, the contrast estimated from the *ICC* matrices (i.e., *ICCC*) when considering all 1560 scans was higher compared to the *ICCC* estimated from the smaller groups of 39 subjects each (Fig. 2d). Interestingly, we observe that a large number of BNEs, and especially edges between the default mode and fronto-pariental networks, exhibited low FC values but high *ICC* values.

##### Intraclass correlation contrast (ICCC)

The metric *ICCC* proposed in this work to assess subject specificity in the fMRI data, showed an increasingly monotonic behavior in the range of mild pipelines, for both WM and CSF (with and without GSR), reaching a plateau at about 30 components, followed by a small decline for the most aggressive pipelines. However, as shown in Suppl. Fig. 2 (a & d), *FCC* exhibited higher *Z*-scores compared to *ICCC* and, therefore, contributed the most to the scores in the signal-related summarized metric *QC*_signal_ (Fig. 3a), which was defined as the average *Z*-score between *FCC* and *ICCC*.

Fig. 5 shows the *ICC* matrices for the raw data and four preprocessed fMRI datasets estimated using all 1560 scans. It can be seen that in the case of the raw data the *ICC* values were high for all edges, which resulted in a low *ICCC* score. On the other hand, when an aggressive pipeline was used (e.g. 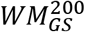), the *ICC* values for most of the BNEs dropped to significantly lower values compared to the rest of the edges leading to an increase in *ICCC* score. Nevertheless, even with aggressive pipelines, many BNEs, and particularly edges corresponding to interactions between the default mode and fronto-parietal networks, demonstrated high *ICC* scores even though the corresponding edges in the group-level FC matrix exhibited low correlation values (Fig. 5). Similar results were observed for the Seitzman and MIST atlases (Suppl. Fig. 8-Suppl. Fig. 9). In addition, note that the *ICCC* scores reported in Fig. 5 were higher compared to the *ICCC* scores extracted from the smaller groups of subjects shown in Fig. 2d (groups of 39 subjects each). Also, differences in *ICCC* between pipelines found when scores were obtained for each group of subjects separately were decreased when *ICCC* was obtained from all subjects in one step (e.g. differences in *ICCC* scores between *GS* and *FIX*_*GS*_). The aforementioned property of the metric *ICCC* suggests that its sensitivity in comparing the performance between preprocessing strategies decreases when larger number of subjects is considered.

#### Motion-related QC metrics

##### FD-FCC

The raw data yielded a mean *FD*-*FCC* score of −0.42 implying that lower levels of motion were associated with higher *FCC* scores (Fig. 2b). Importantly, when performing WM denoising with more than 30 components, the strength of *FD*-*FCC* dropped to about −0.15 (*Z*-score 2.2). Fig. 6 shows scatterplots of mean *FD* vs *FCC* for the first scan of 370 subjects (20 subjects that demonstrated a mean *FD* three median absolute deviations (MADs) above the median were excluded). We observe that even though GSR applied alone on the raw data improved the *FCC* score, it also increased the negative correlation between mean *FD* and *FCC* or, in other words, it enhanced the dependence of *FCC* score on the levels of motion (*r* = −0.46; *p* = 10^−19^). However, when GSR was performed along with WM denoising of 200 regressors 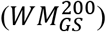 the negative correlation between *FD* and *FCC* almost vanished (*r* = −0.11; *p* < 0.04).

**Fig. 6.**
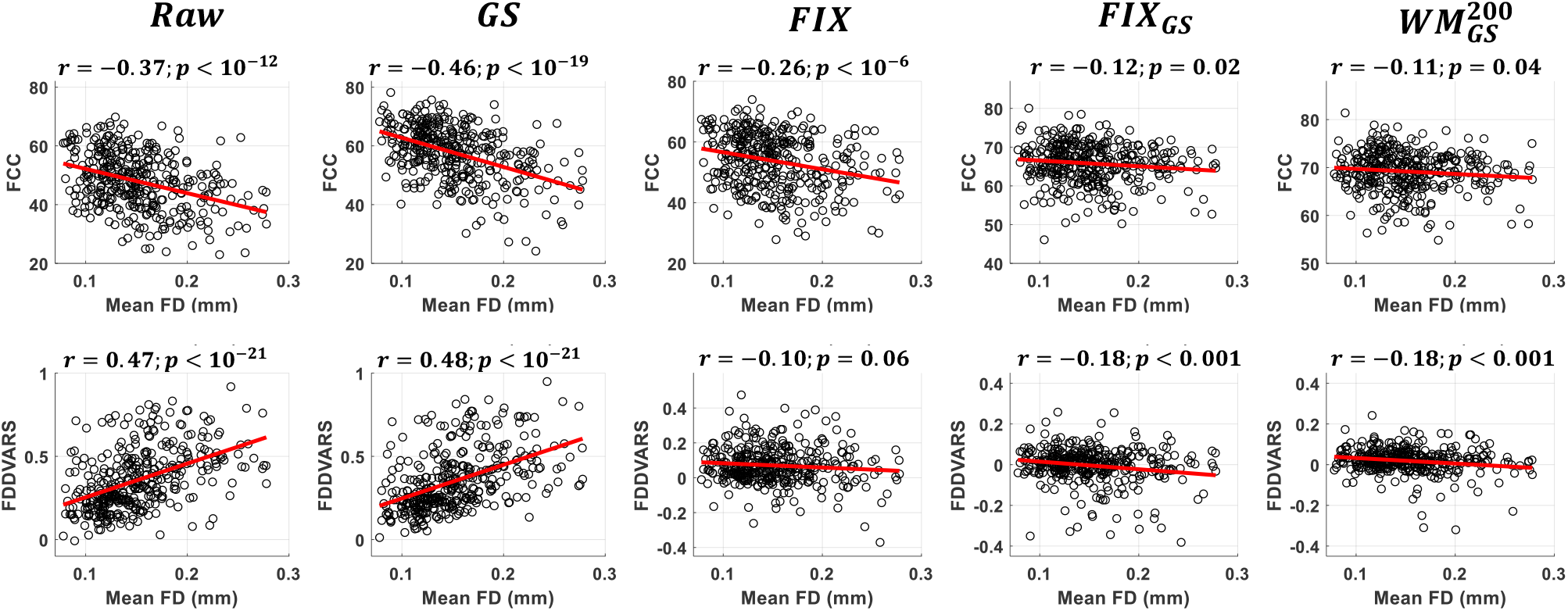
Scatterplots of mean FD vs FCC (top) and mean FD vs FDDVARS (bottom) considering the first scan from all subjects*. In the case of raw data, higher levels of motion in a scan made network identification more difficult (lower FCC scores) and resulted in significantly larger motion artifacts in the fMRI data (higher FDDVARS values). Using the pipelines 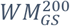 and *FIX*, resulted in a significant reduction of the dependence of FCC and FDDVARS on the levels of motions. *Scans with mean FD three median absolute deviations (MADs) above the median were excluded (based on this criterion, 20 out of 390 subjects were excluded).

##### FDDVARS

The raw data demonstrated an *FDDVARS* score of 0.37, suggesting that the parcel time series were strongly contaminated by motion artifacts (Fig. 2e). WM denoising with 200 components 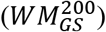 was able to drop *FDDVARS* to 0.02, corresponding to a *Z*-score of 11.3. Suppl. Fig. 10-Suppl. Fig. 11 present traces of *FDDVARS* from individual scans, where it can be appreciated that WM denoising led to a strong reduction of motion artifacts during time windows with high levels of motion. Note that *FDDVARS* exhibited significantly higher *Z*-scores than the rest of the motion-related QC metrics (Suppl. Fig. 2e) and, therefore, contributed the most to the scores of the summarized metric *QC*_motion_ (Fig. 3).

##### FD-FDDVARS

The raw data exhibited a mean correlation *FD*-*FDDVARS* of 0.48 (Fig. 2f), implying that higher levels of motion in a scan resulted in significantly larger motion artifacts in the fMRI data. The set of regressors 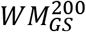 achieved the smallest absolute score of *FD*-*FDDVARS* (score: −0.07), corresponding to a *Z*-score of 2.7. Fig. 6 shows scatterplots of the mean *FD* vs the *FDDVARS* score (i.e., *FD*-*FDDVARS*) for the raw data and four different preprocessed datasets considering the first scan from 370 subjects (20 subjects were excluded due to extreme values in mean *FD*). As we can see from the raw data, the levels of motion during a scan (as evaluated with mean *FD*) had a strong effect on *FDDVARS*, which reflects the degree of motion artifacts in the fMRI data (*r* = 0.47; *p* < 10^−21^). The pipelines *FIX*_*GS*_ and 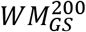 were able to reduce the correlation of *FD*-*FDVARS* to −0.18 (*p* < 0.001). We also observed that *FIX* achieved a lower correlation *FD*-*FDDVARS* of −0.10 compared to *FIX*_*GS*_ and 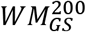. However, *FIX* exhibited also larger absolute scores of *FDDVARS* than *FIX*_*GS*_ and 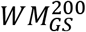.

##### *FDFC*_*median*_

When GS was included in the sets of regressors, the scores for *FDFC*_median_ exhibited a monotonically decreasing trend for an increasing number of components, beginning at 0.15 for raw data (*Z*-score: 0.1) and reaching 0.13 (*Z*-score: 0.7) for both *FIX*_*GS*_ and 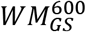 (Fig. 2g). However, when GS was not included in the preprocessing, increasing the number of WM components from 1 to 7 PCA regressors resulted in an increase for *FDFC*_median_ from 0.15 to 0.22; for an even higher number of components, FDFC_median_ decreased, reaching 0.13 in the case of 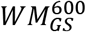.

##### *FDFC*_*dist*_

In the case of raw data, *FDFC*_dist_ was equal to −0.16, which reflects that short-distance pairwise correlations were more severely confounded by motion than long-distance correlations (Fig. 2i). Increasing the number of components in WM denoising resulted in a decrease in the correlation values, with the more aggressive sets *WM*^600^ and 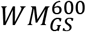 achieving the minimum *FDFC*_dist_ scores of −0.01 and −0.04. However, the associated *Z*-scores for the latter two sets were relatively low (1.1 and 1.0; Suppl. Fig. 2i) and, as a consequence, *FDFC*_dist_ did not have a significant weighting on the CQC metric.

##### FD-MFC

*FD*-*MFC* was proposed in this work and is based on the assumption that the more a subject moves during a scan, the higher the mean value of correlations in the FC matrix is. As we can see in Fig. 2h, the score for *FD*-*MFC* in the case of raw data was 0.22, confirming that motion can inflate the estimated correlations in the FC matrix. Importantly, when GSR was not performed, increasing the number of WM components from 1 to 30 led to an increase of *FD*-*MFC*, with *WM*^30^ exhibiting an *FD*-*MFC* score of 0.45. For a higher number of WM components, *FD*-*MFC* decreased monotonically reaching 0.19 for *WM*^600^. Overall, when WM denoising was combined with GSR, a lower *FD*-*MFC* was achieved, with 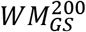 yielding a score of −0.06 (*Z*-score: 0.7). Similar results were found when *FD*-*MFC* was estimated using the first scan from all subjects, even though there was a mild decrease in the scores for all pipelines (Fig. 7).

**Fig. 7.**
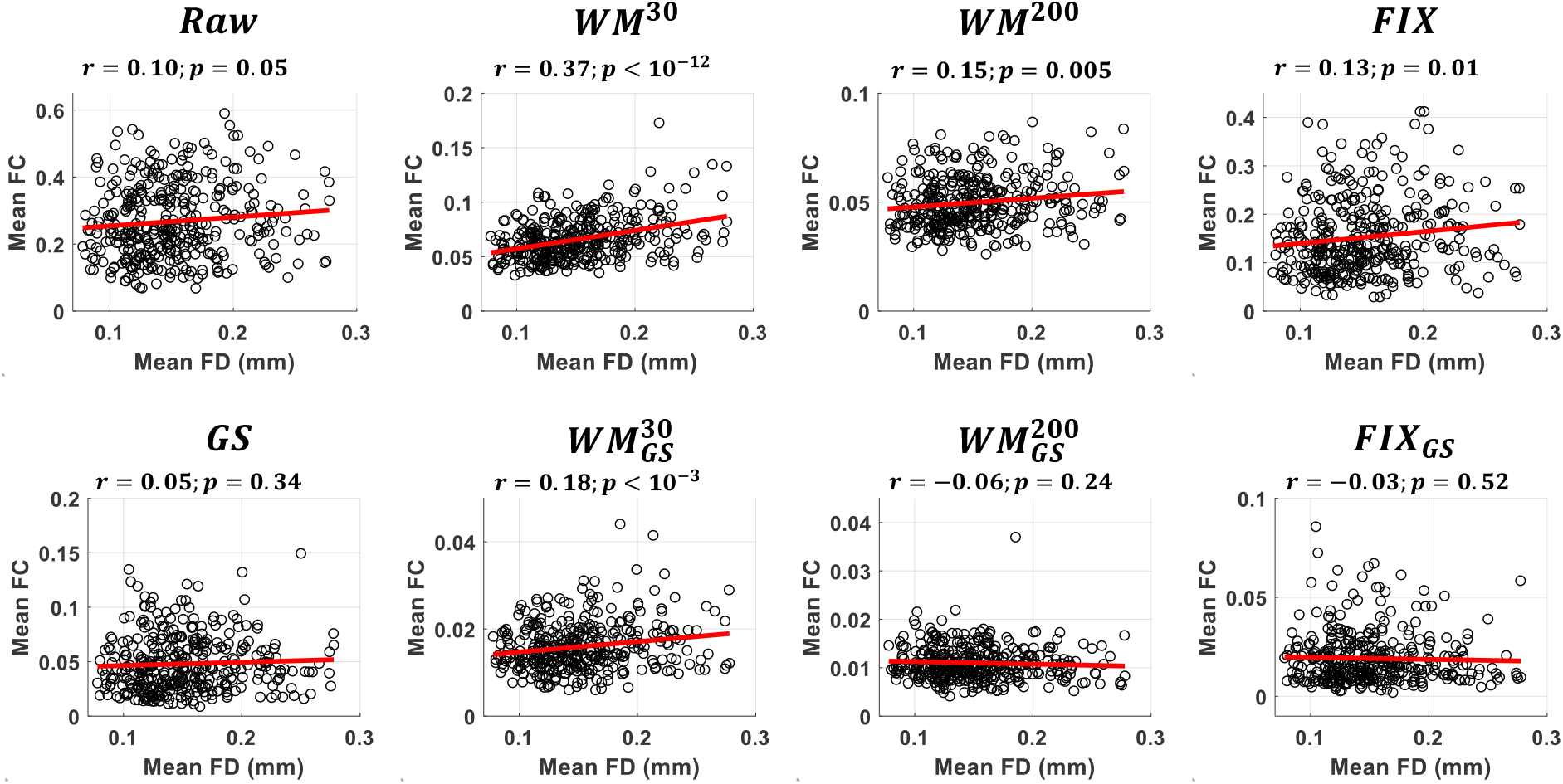
Scatterplots of mean *FD* vs mean FC for different pipelines with (bottom) or without (top) GSR considering the first scan from all subjects*. In the case of raw data, scans with high levels of motion were associated to high correlation values in FC. This dependence on the levels of motion vanished when the data were preprocessed with 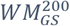 or *FIX*_*GS*_. Importantly, when a relatively low number of components were removed (e.g., *WM*^30^), the effect of motion was enhanced compared to the raw data. *Scans with mean *FD* three median absolute deviations (MADs) above the median were excluded (based on this criterion, 20 out of 390 subjects were excluded).

### 3.2 Evaluation of data-driven NCTs

In this analysis, we used the QC metrics to compare twenty different pipelines involving the removal of data-driven nuisance regressors from the fMRI data (Table 1). Fig. 8 shows the scores for the summarized metrics *QC*_signal_ and *QC*_motion_, as well as the combined metric *CQC* for the Gordon atlas (the results for the Seitzman and MIST atlas are shown in Suppl. Fig. 12-Suppl. Fig. 13). Looking at the first three pipelines that correspond to the 6, 12 and 24 MPs, we observe that motion regressors reduced the effect of motion and (to a less extent) improved the SNR in the data, with the more aggressive pipeline (24 MPs) exhibiting the strongest impact for all three atlases. GSR alone (pipeline 4) significantly improved the SNR even though, for the Seitzman and MIST atlas, it also led to a small decrease in the *QC*_motion_ score (Suppl. Fig. 12-Suppl. Fig. 13). As can be seen from Suppl. Fig. 3-Suppl. Fig. 4, *FD*-*FCC* and *FD*-*MFC* increased after GSR, while *FDDVARS* was at similar or lower levels compared to the raw data, suggesting that even though there was not any enhancement of motion artifacts, the systematic differences across scans due to motion increased.

**Fig. 8.**
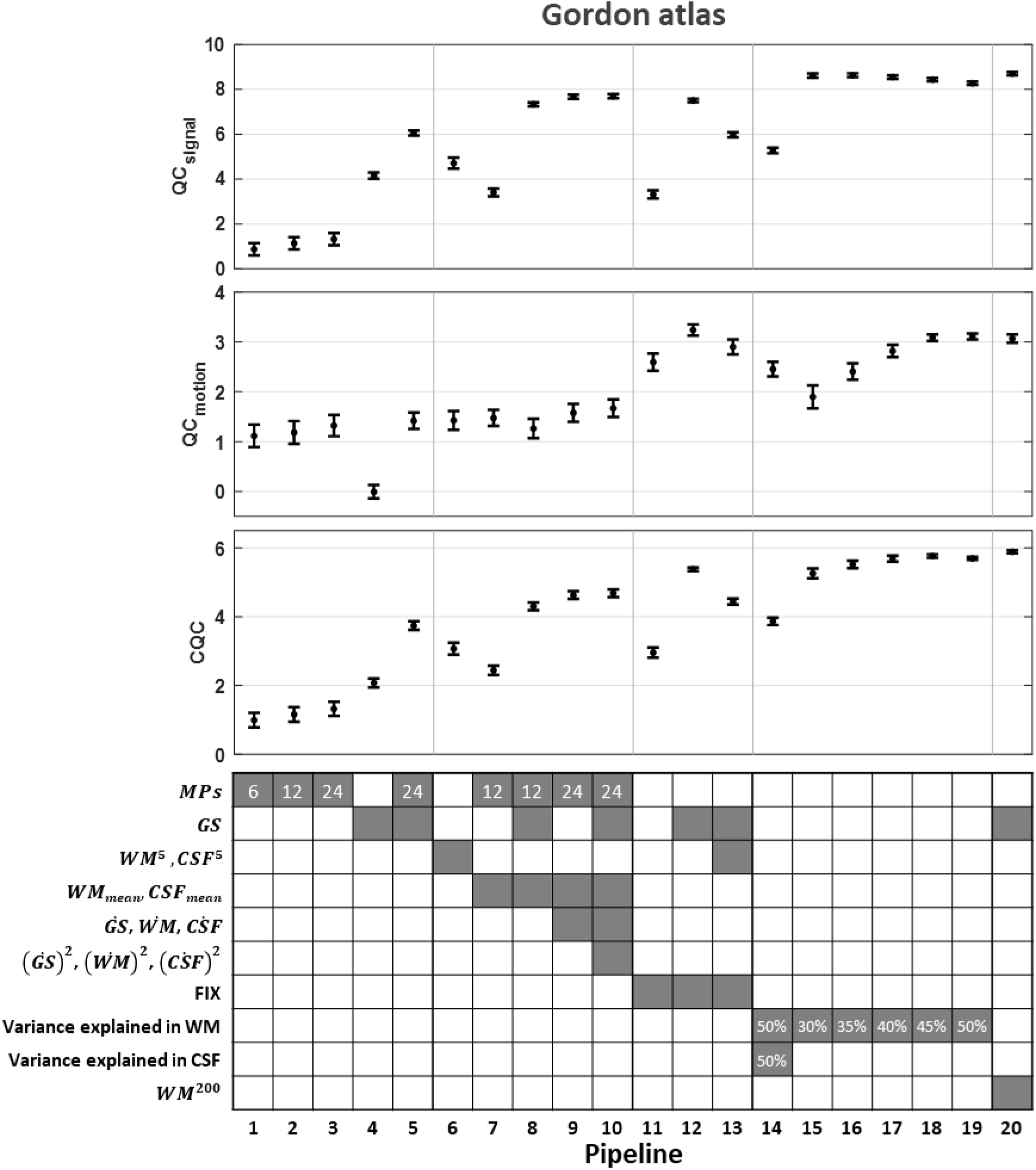
Evaluation of data-driven NCTs (Gordon atlas). Twenty different data-driven pipelines were examined, as listed in Table 1. Among all pipelines, pipelines that included GSR and WM or FIX denoising yielded the highest scores in *QC*_signal_, *QC*_motion_ and *CQC* (i.e., pipelines 12 and 17-20).

Several studies employ aCompCor as NCT, removing five WM and five CSF regressors (Wang et al., 2017; Xiao et al., 2016). Our results derived from the HCP data suggest that this set of regressors demonstrates a moderate improvement with respect to both *QC*_signal_ and *QC*_motion_ (pipeline 6). Similar improvement in the quality of data was achieved when the mean time series from WM and CSF, and the 12 MPs were regressed out (pipeline 7; Urchs et al., 2017) whereas when including also the GS to the set of regressors, the *QC*_signal_ score reached a higher value (pipeline 8; Finn et al., 2015). Pipelines 9 and 10 were more aggressive variants of pipeline 8, including 24 instead of 12 MPs, as well as the derivatives and squared terms of the tissue mean time series from GM, WM and CSF (Ciric et al., 2017; Laumann et al., 2017; Xia et al., 2018). Considering more nuisance regressors in the preprocessing (36 rather than 15 regressors), pipeline 10 exhibited a small but significant improvement compared to pipeline 8, with respect to *QC*_signal_ and *QC*_motion_ for the Gordon and Seitzman atlases, whereas for the MIST atlas *QC*_motion_ increased and *QC*_signal_ slightly decreased.

Pipelines 11 to 13 evaluated the data quality for the FIX-denoised data provided in HCP with and without further denoising (Fig. 8). We observe that, as proposed in Burgess et al. (2016), regressing out the GS from the FIX-denoised data improved both *QC*_signal_ and *QC*_motion_ scores (pipelines 11 vs 12). However, when five WM and five CSF regressors were removed in addition to the GS (pipeline 13; Siegel et al., 2017) both summarized metrics were lower compared to performing only GSR (pipeline 12).

Pipeline 14 was based on the NCT recommended by Muschelli et al. (2014), which considers as set of regressors the necessary number of WM and CSF regressors needed to explain 50% of variance in their associated compartments. As we see, in all three atlases pipeline 14 achieved a satisfactory reduction in motion artifacts, even though the SNR was much lower compared to other pipelines. Earlier results presented here showed that, the scores of the *QC*_signal_ metric were relatively low when CSF denoising was performed and high for WM denoising, particularly when GSR was also performed (Fig. 3). Based on these results, we also considered pipelines 15 to 19 that consider the GS as well as the WM regressors needed to explain a predefined fraction of variance in WM ranging from 30 to 50%. Our results suggest that pipelines 15 to 19 achieved high scores for both *QC*_signal_ and *QC*_motion_, with the highest scores achieved when 45-50% of the variance was used as a threshold to select the WM regressors.

Finally, the set of regressors 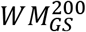 that was found in the previous section to perform the best was considered as pipeline 20. For all pipelines, we observed that the highest QC scores were obtained when GSR was performed in combination with FIX or WM denoising (i.e., pipelines 12 and 17-20).

### 3.3 Evaluation of model-based (motion and physiological) NCTs

Four sets of model-based regressors were examined with respect to improvement in SNR and reduction of motion artifacts and biases. The four sets were related to head motion (24 MPs), cardiac pulsatility, breathing motion and SLFOs To assess their contribution when tissue-based regressors were also included in the set of nuisance regressors, we examined 64 pipelines presented in Fig. 9 in the form of a design matrix that refers to combinations of the four sets of model-based regressors, the GS and a set of 200 PCA regressors from WM. When only model-based regressors were considered, accounting for SLFOs improved the *QC*_signal_ score, whereas correcting for either head or breathing motion improved both *QC*_signal_ and *QC*_motion_ scores. Accounting for cardiac pulsatility led to an increase in *QC*_signal_ and decrease in *QC*_motion_, even though the effect of cardiac regressors was lower compared to the rest of the model-based regressors. Finally, when GS and 200 WM regressors were considered 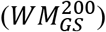, accounting also for breathing motion, cardiac pulsatility or SLFOs, using model-based regressors, did not have any impact on the data quality, whereas correcting for motion with the 24 MPs led to a small decrease in the score for *QC*_motion_.

**Fig. 9.**
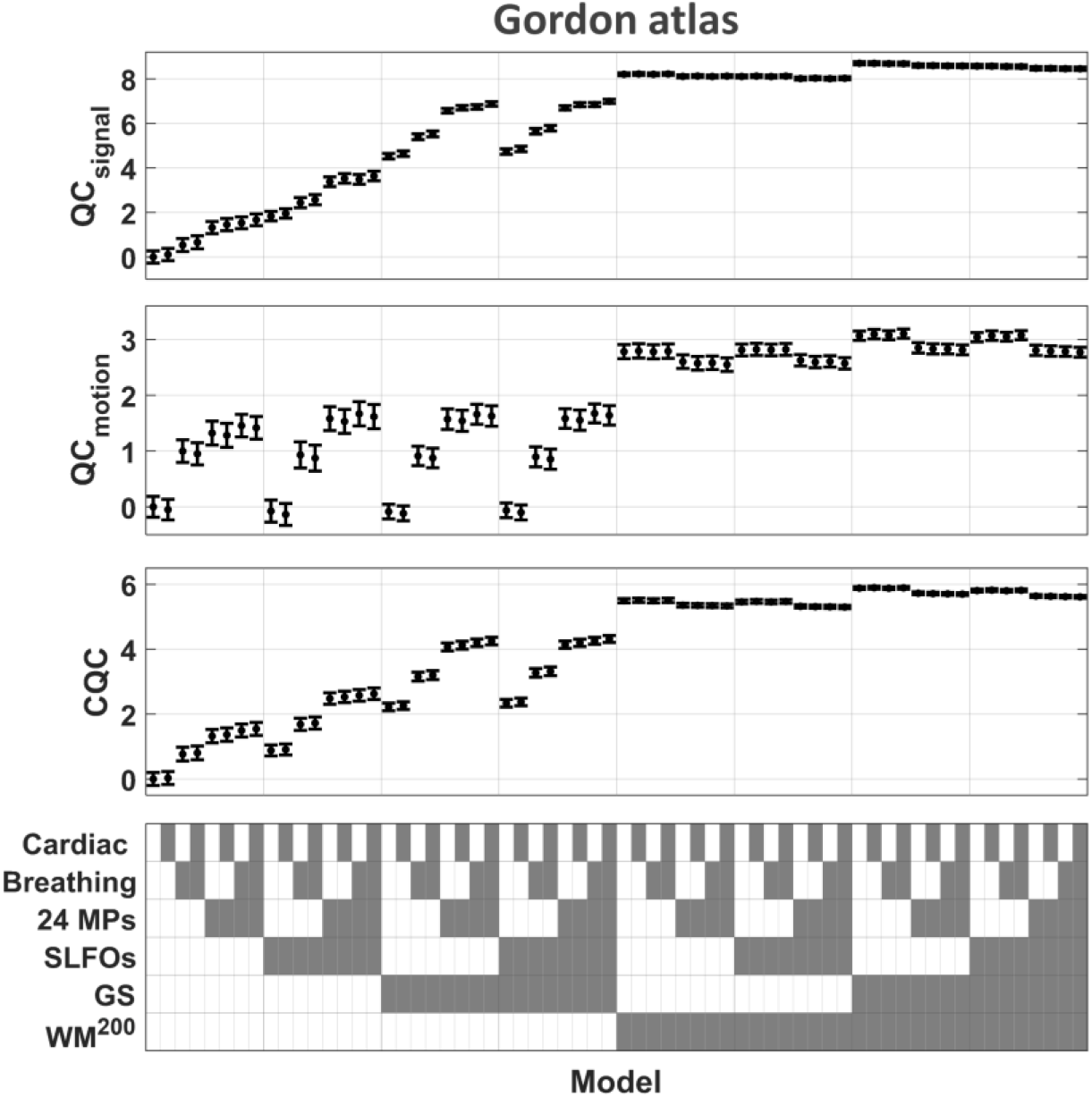
Evaluation of model-based NCTs (Gordon parcel space). Model-based regressors were obtained from the motion (realignment) parameters and physiological recordings to correct for artifacts due to head motion (24 MPs), cardiac pulsatility (Cardiac), breathing motion (Breathing) and SLFOs (i.e., BOLD fluctuations due to changes in heart rate and respiratory flow; Kassinopoulos and Mitsis, 2019). Overall, none of the examined model-based NCTs contributed further to data quality improvement beyond that achieved with the set of tissue-based regressors 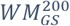. Similar results were found using the data in the Seitzman and MIST parcel space (Suppl. Fig. 14).

### 3.4 Evaluation of scrubbing

Discarding volumes contaminated with motion artifacts before regressing out the set of nuisance regressors 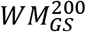 did not provide any gain with respect to the fMRI data quality (Fig. 10). Instead, stricter threshold values *FD*_thr_ resulted in lower scores for *QC*_signal_ and *QC*_motion_. Also, discarding volumes with *DVARS* values beyond the threshold did not have any impact on the *QC*_motion_ score, while it significantly decreased *QC*_signal_.

**Fig. 10.**
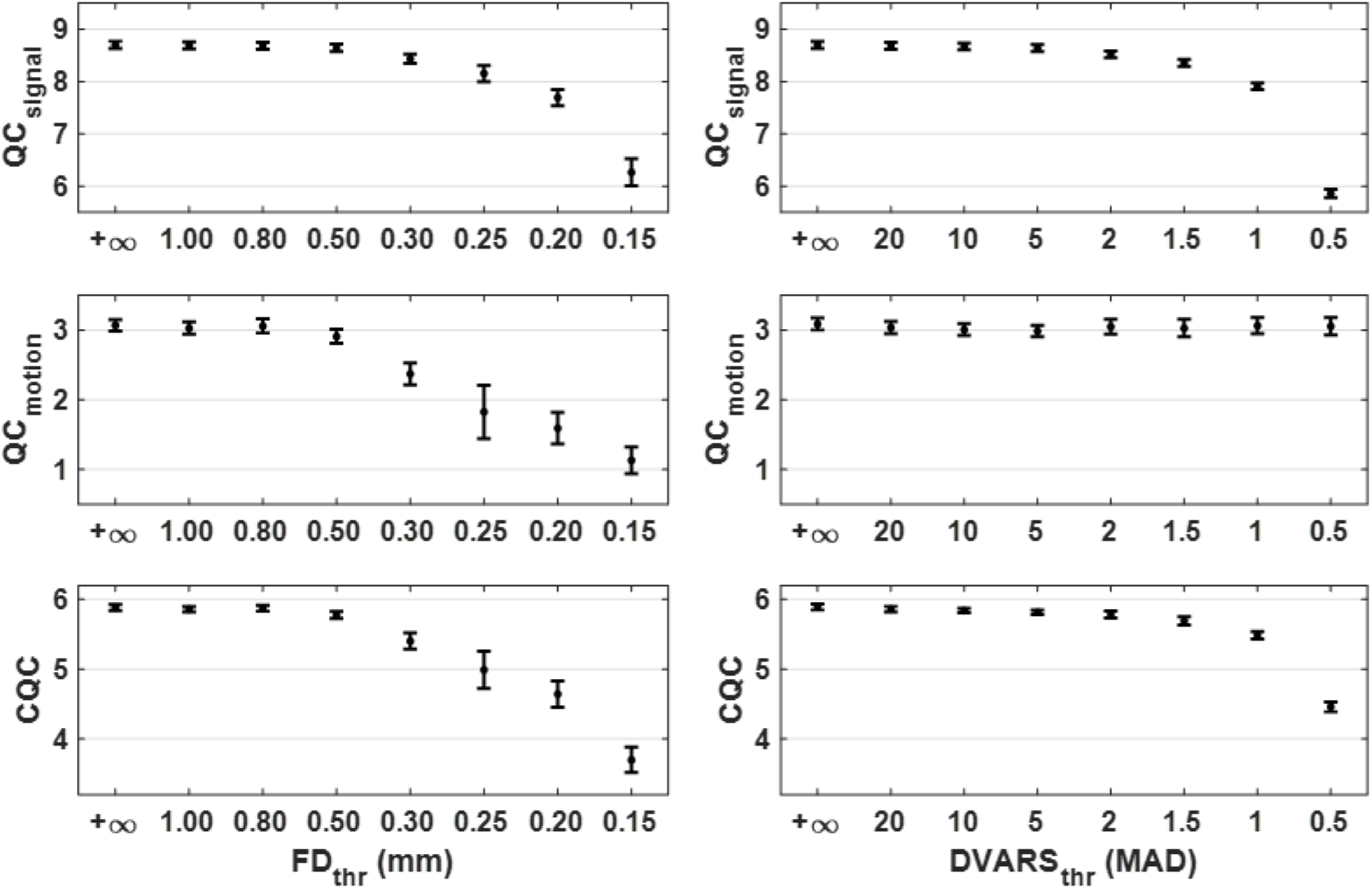
Effect of scrubbing in data quality for different threshold values. The framewise data quality indices *FD* and *DVARS* were used to flag volumes contaminated with motion artifacts. Subsequently, the motion-contaminated volumes were discarded before preprocessing the data with the set 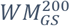 and estimating the *QC*_signal_, *QC*_motion_ and *CQC* scores. The obtained scores for varying values of thresholds *FD*_thr_ and *DVARS*_thr_ are shown on the left and right columns, respectively. For both *FD* and *DVARS* scrubbing, the lower (stricter) were the threshold values, the worse was the resulting data quality. Similar results were found using the data in the Seitzman and MIST parcel space (Suppl. Fig. 15). For the fraction of volumes retained after scrubbing at a given threshold please see Suppl. Fig. 18.

### 3.5 Evaluation of low-pass filtering

To examine the effect of low-pass filtering on data quality as well as its dependence on the cut-off frequency, we repeated the denoising of the data with variants of WM denoising (i.e. 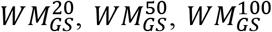 and 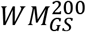) after low-pass filtering the data and the regressors at different cut-off frequencies. For all variants of WM denoising, the highest *CQC* scores were achieved when low-pass filtering was performed at 0.20 Hz (Fig. 11). At this cut-off frequency, the sets of regressors 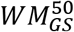 and 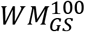 outperformed the larger set 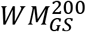. In addition, we observed that the lower cut-off frequency 0.08 Hz that is commonly used in the literature, as well as the cut-off frequency 0.05 Hz, led to a decrease in the *CQC* score which was attributed to a reduction in *QC*_signal_. The pipelines that consisted of a low-pass filtering at 0.20 and removal of 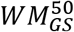 or 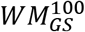 were among the pipelines that achieved the highest *CQC* scores for the Seitzman and MIST atlas as well (Suppl. Fig. 19). Overall, we observed that mild variants of WM benefitted the most from low-pass filtering, particularly in terms of reducing motion artifacts and biases (Fig. 11, Suppl. Fig. 19).

**Fig. 11.**
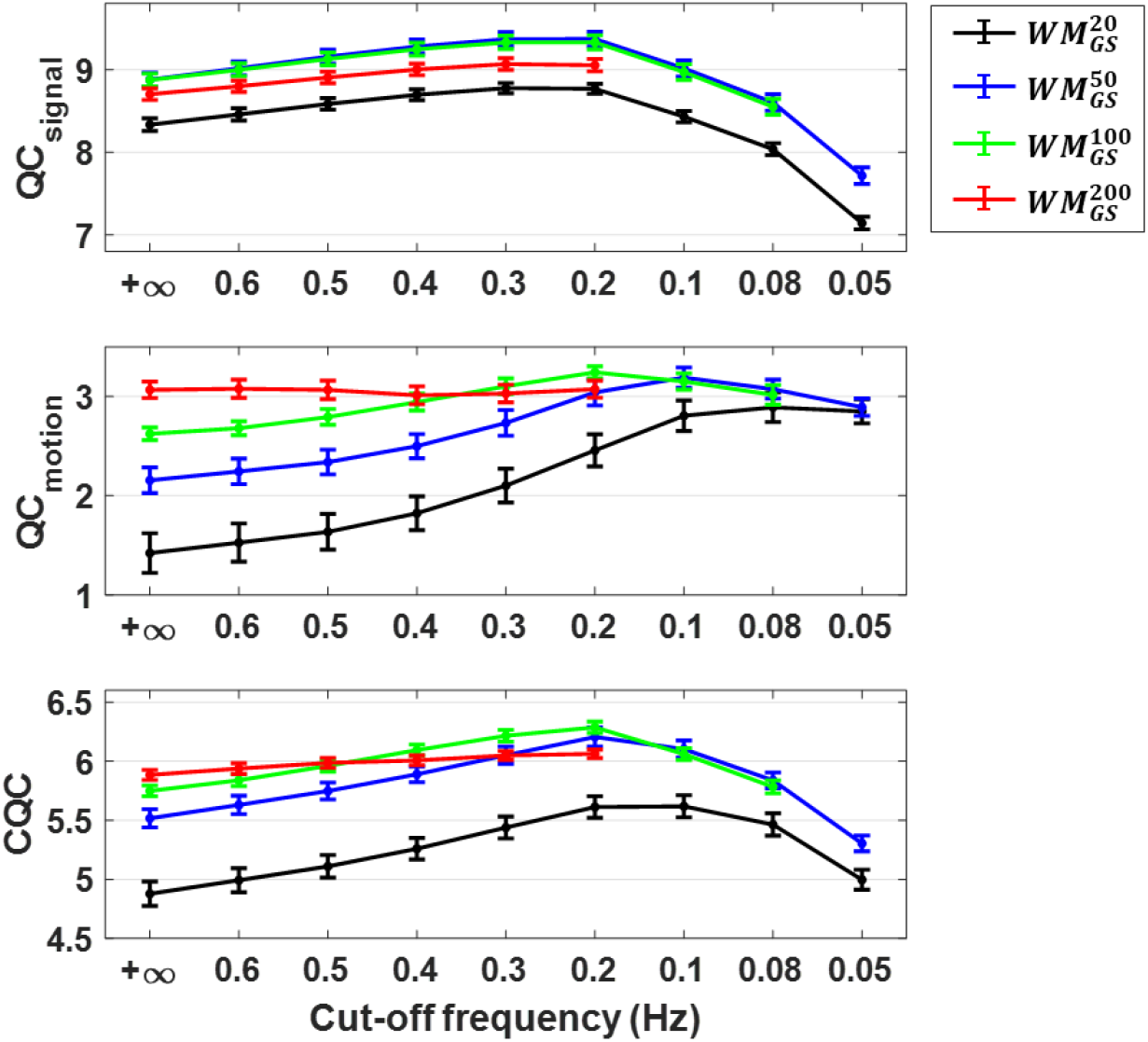
Effect of low-pass filtering in data quality for different cut-off frequencies. For all variants of WM denoising examined, low-pass filtering with a cut-off frequency of 0.2 Hz yielded the highest *CQC* score. At this cut-off frequency, 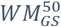 and 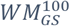 yielded the highest CQC scores with mean scores 6.2 and 6.3, respectively. No significant difference was observed between 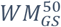 and 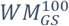 (*p* > 0.05, two-sample *t*-test). The cut-off frequency of 0.2 Hz, and particularly when combined with 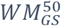 and 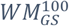, yielded the highest *CQC* score also for the data registered to the Seitzman and MIST atlases (Suppl. Fig. 19). Note that the lowest cut-off frequencies examined for 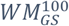 and 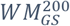 were 0.08 Hz and 0.2 Hz, as for these sets of regressors lower cut-off frequencies lead to fMRI data with a substantially low number of degrees of freedom (Bright et al. 2017). Data that had not been filtered are indicated with a ∞ cut-off frequency.

### 3.6 Identifiability of large-scale networks

Finally, we sought to quantify the identifiability of each of the large-scale networks defined in the three functional atlases employed here and its dependence on the preprocessing pipeline. To this end, we calculated the *FCC* score of each network for the raw dataset as well as four preprocessed datasets. To obtain the *FCC* score per network, for a given network, when estimating the *FCC* score, we compared WNEs with BNEs considering only WNEs that belonged to the examined network (for more information see Section 2.6).

In Suppl. Fig. 20, we see that for the Gordon and Seitzman atlases, there was larger variability in *FCC* score across networks rather than across pipelines. Networks consisting of a small number of parcels, such as the salience network in the Gordon atlas and the medial temporal lobe network in Seitzman atlas, exhibited small negative *FCC* scores for the raw data, whereas when the data were preprocessed with a pipeline that included GSR the *FCC* scores were increased to small positive values. On the other hand, large networks such as the default mode network exhibited significantly higher *FCC* scores.

In the case of the MIST atlas there was less variability in *FCC* score across networks compared to the Gordon and Seitzman atlas, which may be due to the fact that these networks consisted of a similar number of parcels. However, two out of the seven networks demonstrated a somewhat unexpected behavior. Specifically, the mesolimbic network demonstrated a large negative *FCC* score for the raw and FIX-denoised data, despite the fact that it consists of a similar number of parcels to other networks in the atlas. Furthermore, regarding the cerebellum network, even though the *FCC* score in the raw data was relatively high, when FIX denoising was applied the *FCC* score dropped to zero.

Finally, while some networks in the three atlases were assigned the same name, they did not necessarily demonstrate the same behavior in terms of differences in *FCC* across the five fMRI datasets. For example, in the Gordon atlas we observe that the fronto-pariental network yielded the highest *FCC* score when the data were preprocessed using 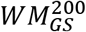, whereas in the MIST atlas, for the same network, the highest *FCC* score was observed with the raw data. Nevertheless, for the majority of networks, *FCC* scores were maximized when preprocessing was done with *FIX*_*GS*_ or 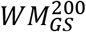.

## 4. Discussion

In this study, we have rigorously examined the effects of different preprocessing steps on SNR and degree of motion artifacts and biases in resting-state fMRI data, focusing on functional networks. As in previous studies, the QC metrics used to compare preprocessing pipelines illustrated different trends between them (Fig. 2). Therefore, to facilitate the comparison across pipelines, we introduced a novel framework that initially normalizes each of the 8 QC metrics to *Z*-scores so that they reflect relative improvement in standard deviations with respect to the raw data. Subsequently, the two normalized signal-related metrics *FCC* and *ICCC*, and the six normalized motion-related metrics FD-FCC, *FDDVARS*, *FD*-*FDDVARS*, *FDFC*_median_, *FDFC*_dist_ and *FD*-*MFC* are averaged to obtain the metrics *QC*_signal_ and *QC*_motion_, respectively. Finally, the combined QC metric *CQC* defined as the mean of the *QC*_signal_ and *QC*_motion_ scores is calculated. Using this framework and resting-state fMRI data from the HCP registered to the Gordon atlas, we found that the largest improvement in the score of *CQC* was obtained when the GS and 200 PCA regressors from WM were regressed out (Fig. 3). Similar results were found when the fMRI data were registered to the Seitzman and MIST atlases (Suppl. Fig. 3-Suppl. Fig. 6). Note that 200 WM regressors correspond to about 17% of the regressors derived with PCA from WM as the fMRI scans consisted of 1160 volumes each, and explain on average 36±6% of the variance in the WM voxel time series (Suppl. Fig. 1).

Although we considered only subjects with good quality physiological data in all four scans, none of the model-based techniques examined here yielded further improvement in terms of data quality when compared to WM denoising (Fig. 9). Note that similar conclusions were derived from a subsequent study from our lab in which the effects of physiological processes on FC were more systematically examined (Xifra-Porxas et al., 2021). Specifically, in Xifra-Porxas et al. (2021), we provided evidence that data decomposition techniques (FIX and WM denoising) combined with GSR lead to a substantial mitigation of the effects of head motion and physiological fluctuations on FC but also improve connectome-based subject discriminability. In addition, when data decomposition techniques were considered, model-based preprocessing approaches did not provide any additional benefit. These observations may not be surprising as it has been previously shown that artifacts due to head motion and physiological fluctuations can be corrected by removing WM and CSF regressors (aCompCor) as well (Behzadi et al., 2007; Muschelli et al., 2014). Also, WM denoising, and in general model-free approaches such as FIX (Salimi-Khorshidi et al., 2014) and AROMA (Pruim et al., 2015b), have the benefit that they do not require physiological data and are not based on any assumptions related to the mechanisms by which physiological processes affect the BOLD signal. For example, the convolution models used here to account for the effect of heart rate and breathing pattern assume that a linear stationary system can describe these effects, which may not be entirely true (Kassinopoulos and Mitsis, 2019).

Even though the model-based techniques were not found to yield any additional improvement as compared to data decomposition techniques, it is important to bear in mind that the QC metrics considered in the present study are mainly intended for whole-brain FC studies and, thus, are not necessarily informative or even applicable in different contexts. Model-based techniques are of great importance in several cases, such as studies with limited filed-of-view (e.g. laminar fMRI, brainstem imaging) where data-driven techniques (e.g. FIX denoising) cannot be directly used. Furthermore, model-based techniques are useful in studies of the autonomic nervous system (Kassinopoulos et al., 2021; Mulcahy et al., 2019) and task-based studies whereby physiological noise is often correlated to the signal of interest (Glasser et al., 2018) and, therefore, conservative approaches based on concurrent physiological recordings and physiologically-inspired models are needed to account for the associated noise, while also preserving the signal of interest.

Performing scrubbing before WM denoising 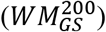 was found to deteriorate the quality of the data rather than improving it (Fig. 10). As this result contradicts with the benefits of scrubbing reported in the literature (Ciric et al., 2017; Gratton et al., 2020; Parkes et al., 2018; Power et al., 2015), we repeated the analysis using standard aCompCor, which removes a significantly smaller number of regressors (i.e. five WM and five CSF regressors), and found that scrubbing at specific threshold values (*FD* > 0.20 mm and *DVARS* > 1.5 MAD) improved the data quality with respect to motion-related QC metrics and, to a smaller extent, signal-related metrics (Suppl. Fig. 16). However, our findings suggest that the improvement in data quality observed with mild preprocessing and scrubbing was still inferior to that obtained with more aggressive pipelines (e.g. WM denoising and FIX) that are not preceded by scrubbing.

Recent studies have suggested that in multi-band fMRI datasets with short TR, such as the HCP dataset, the effect of breathing activity on *FD* traces should be suppressed to ease the identification of motion-contaminated volumes and avoid removing presumably noise-free volumes (Gratton et al., 2020; Power et al., 2019). With this in mind, we removed the effects of breathing from the six motion realignment parameters using concurrent breathing recordings and 3^rd^ order RETROICOR, and estimated *FD* traces free of breathing effects. Consistent with previous studies (Gratton et al., 2020; Power et al., 2019), these *FD* traces were characterized by lower amplitude and smaller oscillatory fluctuations and yielded in turn lower numbers of motion-contaminated volumes for a given threshold value (Suppl. Fig. 18). However, when the data were preprocessed using 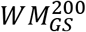, even with the proposed *FD* traces, scrubbing was found to deteriorate the quality of the data (Suppl. Fig. 17). Note that, for the motion-related metrics examined in this study that are based on *FD*, we considered the standard *FD* traces, which include breathing-related fluctuations. Breathing, apart from shifting the acquired images due to disturbances in the magnetic field, can also alter the intensity in fMRI voxel time series (Raj et al., 2001, 2000), and these alterations are not corrected via volume realignment. Therefore, the standard *FD* traces were used in the QC metrics to render them sensitive to the presence of breathing-induced artifacts.

Finally, we found that low-pass filtering at 0.2 Hz led to further improvement in data quality beyond the improvement achieved with WM denoising and, importantly, this improvement was more prominent for mild variants of WM denoising (i.e. 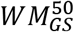 and 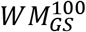; Fig. 11). However, a substantial decrease in SNR was observed when the 0.08 Hz cut-off frequency commonly used in fMRI studies was considered. The rationale behind choosing the 0.08 Hz cut-off frequency for low-pass filtering in resting-state FC is that well-established large-scale networks have been found to oscillate at frequencies below 0.10 Hz (Damoiseaux et al., 2006), while breathing motion and other sources of noise appear at frequencies above this frequency (Caballero-Gaudes and Reynolds, 2017; Liu, 2016). Nevertheless, several studies have found activity in RSNs in the range from 0.1 to 0.5 Hz (Chen and Glover, 2015; Niazy et al., 2011), suggesting that low-pass filtering at 0.08 Hz may potentially remove signal of interest. Based on our results, when considering whole-brain FC, low-pass filtering at 0.2 Hz yields the highest SNR, which may be related to reduction in breathing motion artifacts that appear at around 0.3 Hz and may not be fully corrected with WM denoising.

### 4.1 QC metrics

Nine QC metrics were initially considered with three metrics related to the SNR in the fMRI data and six metrics related to motion artifacts and biases. To assess the sensitivity of each metric, the subjects were split into 10 groups of 39 subjects each with similar levels of motion across groups, as assessed with within-scan mean FD. Subsequently, the QC scores were estimated for each group separately. Based on the fact that the 10 groups of subjects were characterized by similar distributions of mean *FD* values, we hypothesized that more sensitive QC metrics would be associated with a lower variability (or standard deviation) of scores across groups. Furthermore, to obtain QC metric values that are easier to interpret and compare, the score for a given metric and group of subjects was expressed as a *Z*-score, which reflects the improvement in standard deviations compared to the distribution of values found in the raw data across the ten groups of subjects (for more information see Section 2.7).

To keep the computational load of the present study feasible, we considered three different functional atlases based upon hard non-overlapping parcellations, which do not account for cross-subject variations in the parcel locations (Seitzman atlas) and boundaries (Gordon and MIST atlas), and may therefore provide incomplete descriptions of the brain functional organization (Pervaiz et al., 2020; Van Essen and Glasser, 2018). However, the QC metrics can be easily extended to data-driven soft parcellations as well (e.g. spatial ICA), provided that the main functional connections expected to be observed at the group level, as well as their strength, are known.

#### Signal-related metrics

Among the three signal-related metrics (i.e., metrics related to the SNR), *FCC* demonstrated a substantially higher improvement in *Z*-score compared to the other two metrics (Suppl. Fig. 2), and as such was the main determinant of the summarized metric *QC*_signal_. The *FCC* is based on the assumption that the strength of correlation for WNEs in FC is on average larger than BNEs. Previous studies have used similar metrics to assess spatial specificity in FC considering though only interactions between specific regions in the brain rather than whole-brain interactions (Birn et al., 2014; Chai et al., 2012; Muschelli et al., 2014), whereas Shirer et al. (2015) used a metric that compares the correlations of WNEs with correlations between brain regions and regions outside the brain. While we acknowledge that some of the BNEs may correspond to neuronal-related connections, these edges would likely be the minority. In addition, the relative magnitude of within- vs −between- network edges allows the identification of clusters or networks. Therefore, we believe that considering all BNEs to form the null distribution, rather than connections with voxels outside the brain, is more appropriate.

*FCC* can be considered as a measure of segregation, similar to the modularity index adopted in graph theory approaches for assessing the degree to which a network topology can be subdivided into distinct nonoverlapping communities (Rubinov and Sporns, 2010). Note that our results were consistent with an earlier study that used the modularity index as a signal-related QC metric for the evaluation of preprocessing strategies (Ciric et al., 2017). For instance, in the present study, preprocessing approaches based on decomposition techniques (WM and FIX denoising) were characterized by high *FCC* scores, which is in agreement with the high modularity index reported for the ICA-based technique AROMA (Pruim et al., 2015b) and aCompCor in Ciric et al. (2017). However, a main difference between FCC and the modularity index (Ciric et al., 2017; Cisler, 2017) is that FCC is based on *a-priori* information about the large-scale network (community) to which each parcel belongs, whereas the modularity index assigns the parcels to communities for each subject separately, a step that, depending on the algorithm employed, can be computationally demanding (Rubinov and Sporns, 2010).

The signal-related metric *MICC* was used to assess test-retest reliability across the four sessions of each subject in whole-brain FC estimates. However, as in previous studies, more aggressive pipelines were associated with lower *MICC* scores, which has been interpreted as this metric reflecting subject-specificity due to presence of noise rather than signal of interest (Fig. 2, Birn et al., 2014; Parkes et al., 2018). As *MICC* scores did not seem to correspond to SNR, it was excluded from the rest of the analysis. Interestingly, Birn et al. (2014) reported smaller decreases in ICC for significant connections compared to the remaining connections, which was also confirmed in our data (see for example Fig. 5). Therefore, in the present work, based on these findings, we proposed a novel metric termed *ICCC*, which reflects the extent to which *ICC* values in WNEs are higher as compared to BNEs. *ICCC* was found to behave in a similar manner with *FCC* and, thus, later in the analysis it was combined with the *FCC* score to obtain the summarized metric *QC*_signal_.

Note that the metrics *FCC* and *ICCC* assume that each parcel belongs to only one large-scale network and that only parcels from the same network interact with each other, which is an oversimplified description of the brain functional organization and can be potentially misleading, as numerous resting-state fMRI studies have documented interactions between networks. For instance, in healthy controls, a positive intrinsic functional connection has been reported between the amygdala and medial prefrontal cortex, which consist of key nodes of the limbic and default mode network, respectively (Roy et al., 2009). In our work, pipelines that yielded some of the highest *FCC* and *ICCC* scores, such as WM and FIX denoising along with GSR, were also characterised by strong functional connections and connections with high subject-specificity for a large fraction of BNEs (Fig. 5 & Suppl. Fig. 13). Therefore, the use of *FCC* and *ICCC* did not seem to suppress connectivity between brain regions of different networks. Nevertheless, to address the potential limitations of *FCC* and *ICCC*, we would need to incorporate information in QC metrics about both within- and between-network interactions associated with the population examined. However, this is challenging as the literature currently lacks a detailed characterization of the brain functional organization at this level of detail.

Intriguingly, when the data were preprocessed with *FIX*_*GS*_ or 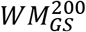, edges corresponding to interactions between the default mode and fronto-parietal networks, despite the low correlation values in group-level FC, demonstrated significantly higher *ICC* values compared to other BNEs (Fig. 5). This finding suggests that regions in the default mode and fronto-parietal networks may be functionally connected but in a subject-specific manner. On a side note, the values of connectivity strength between regions in the aforementioned two networks were found in recent studies to contribute to the identification of individuals using fMRI FC (Finn et al., 2015) as well as to the prediction of behavioral measures (Smith et al., 2015).

A caveat of using *ICCC* as a metric to compare pipelines is that it requires a dataset with several subjects and more than one scan per subject. As a result, in contrast to *FCC*, it cannot be used to assess the data quality for a specific scan. In addition, looking at Fig. 2 & Fig. 5, we see that *ICCC* was increased both with a better preprocessing strategy or with a larger sample size. In addition, when *ICCC* was estimated from all 390 subjects in one step rather than in groups of 39 subjects, apart from the increase in *ICCC* scores for all pipelines, we also observed smaller differences between pipelines which can be translated as a lower sensitivity of *ICCC* when comparing pipelines. We found the dependence of the metric *ICCC* on sample size somewhat puzzling. Due to the aforementioned dependence of *ICCC* sensitivity on the sample size, for future studies with large sample sizes interested in assessing the performance of pipelines, we would recommend estimating *ICCC* in small groups of subjects as done here.

#### Motion-related metrics

Head motion during the scan is a major confound in fMRI FC studies as it diminishes the signal of interest in the data but also affects the strength of connectivity across regions and across populations in a systematic manner. While the majority of edges in FC are typically inflated by motion, short-distance edges tend to be inflated even more than long-distance edges (Satterthwaite et al., 2013). In addition, different populations often exhibit different tendency for motion (e.g., young vs older participants), which has been shown to lead to artificial differences in *FC* (Power et al., 2015). To assess the performance of each preprocessing strategy examined here on the aforementioned aspects of motion effects, three previously proposed metrics (i.e., *FDDVARS*, *FDFC*_median_ and *FDFC*_dist_) as well as three novel metrics (i.e., *FD*-*FCC*, *FD*-*FDDVARS* and *FD*-*MFC*) were considered in the present study. While the main trend in all motion-related metrics was that removing more WM regressors resulted in bringing the scores closer to zero, a different pipeline was favored by each metric (Fig. 2). For example, in the case of WM denoising with GSR, the metric *FD*-*FCC* demonstrated the smallest absolute score when 70 WM regressors were removed, whereas the metric *FDFC*_median_ yielded the smallest absolute score for the most aggressive pipeline examined here (600 WM regressors). However, after normalizing the metrics to *Z*-scores, *FDDVARS* was found to be considerably more sensitive than the remaining metrics (Suppl. Fig. 2) and, thus, was the main determinant of *QC*_motion_. As a result, the summarized metric *QC*_motion_, despite being defined as the average of all six motion-related metrics, favored the set 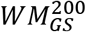 that was characterized by a high *Z*-score of *FDDVARS*.

#### Combined QC metric

While the summarized metric associated to SNR *QC*_signal_ reached a maximum score for a broad range of pipelines (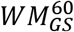 to 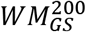), to ensure an efficient mitigation of motion artifacts and biases, the combined QC metric favored the more aggressive option of WM denoising (i.e., 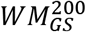; Fig. 3). Due to the lack of ground truth in resting-state fMRI, we cannot be certain that the combined QC metric is successful in identifying the best preprocessing pipeline. However, it is reassuring that the combined QC metric favors data decomposition techniques combined with GSR as these preprocessing techniques have been shown to improve subject discriminability of functional connectomes the most (Xifra-Porxas et al., 2021). In addition, we find reassuring that the combined QC metric favors the inclusion of GSR in the preprocessing, as GSR has been shown to strengthen the association of functional connectomes with behavioral measurements (Li et al., 2019).

We acknowledge that depending on the fMRI study, the researchers may prefer, rather than using the combined QC metric as it is, to give more weighting to the metric *QC*_signal_ and, thus, apply a milder WM denoising, particularly when two populations with similar levels of motion are compared. In addition, we observe that for the majority of analyses, *FCC* and *FDDVARS* favor similar pipelines with the summarized metrics *QC*_signal_ and *QC*_motion_ (e.g. Suppl. Fig. 3 & Suppl. Fig. 5). For example, in the search of the variant of WM denoising and low-pass filter that maximize the SNR, we observe that *FCC* favors the cut-off frequency 0.2 Hz with the set of regressors 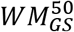 (Suppl. Fig. 22; Gordon and Seitzman atlas), and this finding is consistent with the indications based on the summarized metric *QC*_signal_ (Fig. 9; Suppl. Fig. 19). Similarly, in the search of the pipeline that mitigates the effects of motion the most, FDDVARS is in favor of the cut-off frequency 0.2 Hz with the set of regressors 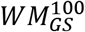 (Suppl. Fig. 22; Gordon and Seitzman atlas), which is also consistent with the indications based on the summarized metric *QC*_motion_ (Fig. 9; Suppl. Fig. 19). Therefore, considering that *FCC* and *FDDVARS* demonstrate significantly higher sensitivity than the rest of the metrics (Suppl. Fig. 2), it would be sensible for a study to determine the preprocessing strategy based solely on these two metrics. If the levels of motion in a cohort are relatively low, the mild pipeline proposed by *FCC* could be employed, whereas in the case that the levels of motion are high, a more aggressive pipeline towards the indications of *FDDVARS* would be more appropriate.

### 4.2 PCA-based WM denoising improves SNR and mitigates motion effects

In the original study introducing the aCompCor technique (Behzadi et al. 2007), the authors proposed the removal of 6 PCA regressors from WM and CSF to account for cardiac and breathing artifacts. However, this proposal was based on Monte Carlo simulations using a modified version of the “broken stick” method described in Jackson (2016), which does not take into account QC metrics that reflect in some way improvement in the quality of the fMRI data. A few years later, Chai et al., (2012) also proposed the removal of five PCA regressors from each noise ROI based on observations related to the connectivity between a region in the medial prefrontal cortex with other brain regions. They also showed that regressing out higher number of PCA regressors led to reduced correlation strengths, which may be associated to reduction of degrees of freedom in the data. Very likely based on these findings, many subsequent fMRI studies considered only 5 PCA regressors from each noise ROI (Ciric et al., 2017; Wang et al., 2017; Xiao et al., 2016).

In this study, we sought to examine the effect of varying the number of PCA regressors on data quality based on QC metrics that account for the effect of motion, as well as the SNR in whole-brain FC rather than interactions between specific regions. Moreover, as there is evidence that neuronal-related activation can be detected in WM (Grajauskas et al., 2019), we examined separately the effects of WM and CSF denoising to determine whether CSF denoising could be sufficient for preprocessing. Interestingly, our results showed that even though WM and CSF denoising achieved similar reduction with respect motion artifacts and biases, the former exhibited substantially better performance in terms of SNR improvement compared to the latter (Fig. 2-Fig. 3). Particularly, the set of regressors 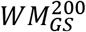, which consists of 200 PCA regressors from WM and the GS illustrated one of the best overall performance among all sets of nuisance regressors examined here (Fig. 8). In addition, our results suggested that a small, albeit significant, further improvement can be achieved when combining mild variants of WM denoising (e.g. 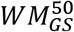) with low-pass filtering at 0.2 Hz.

The standard aCompCor technique that employs 5 PCA regressors from each noise ROI was found to increase the summarized metrics *QC*_signal_ and *QC*_motion_ compared to the raw data but not as much as the set 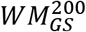 (pipeline 6 vs 20 in Fig. 8). However, we observed that, when GSR was not considered, removing a low number of WM or CSF regressors yielded more negative scores in *FDFC*_median_ and *FD*-*MFC* compared to the raw data, suggesting that the biases in FC due to differences in motion across scans were enhanced (Fig. 2). While this may seem counterintuitive, a possible explanation, based on Fig. 7, is that in raw data high-motion scans are associated with a larger inflation in connectivity due to motion artifacts as compared to low-motion scans. Even though the first few PCA regressors may correct for this inflation, this correction is more effective for low-motion scans which in turn increases the differences in inflation between low- and high-motion scans even more. This phenomenon was not observed for *FDFC*_median_ when GSR was considered and it was diminished for *FD*-*MFC*, suggesting that the inflation in connectivity may be associated to motion-related fluctuations that are also reflected on the GS.

While the practice of regressing out from the data 200 WM regressors may raise concerns with regards to loss of signal of interest, it is important to bear in mind that the examined fMRI data lasted about 15 minutes and had a repetition time TR of 0.72. Therefore, each of the scans examined here corresponded to the relatively large number of 1200 volumes and the voxel time series in WM and CSF were decomposed into 1200 PCA components (note that the first 40 volumes were subsequently discarded to allow modelling of the SLFOs; for more information see Section 2.4). Note also that during the training phase of FIX conducted by the HCP group, the average number of components estimated by ICA was 229 and from these components, on average 205 components were labelled as noisy (Smith et al., 2013a) which is overall in agreement with the optimal number of WM components reported in the present study (200). Nevertheless, to examine the dependence of the optimal number of regressors on the TR, we repeated the aCompCor analysis for a downsampled version of the dataset that was generated by retaining every 4^th^ functional volume of each scan, resulting this way to an effective TR of 2.88 s. Doing so, we found that for this subset of data that is more similar to a conventional fMRI dataset than the original dataset, the best improvement in data quality was achieved with 30 WM regressors combined with GSR (i.e. 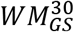; Suppl. Fig. 7). This result suggests that for data consisting of fewer timepoints a lower number of PCA regressors would likely yield the best performance and vice versa.

An alternative preprocessing strategy proposed by Muschelli et al. (2014) is to use the number of PCA regressors needed to explain 50% variance in the two noise ROIs. To compare the performance of this strategy, referred to as aCompCor50, with the original aCompCor they used the QC metric FDDVARS as well as two metrics similar to the *FD*-*FDDVARS* and *FCC* used here. Based on their results, aCompCor50, as compared to aCompCor, exhibited a larger reduction in motion artifacts and improvement in FC specificity, even though the difference for the latter was only marginal when corrected for multiple comparisons. In our dataset, aCompCor50, which corresponded to about 360 (±60) WM and 90 (±40) CSF regressors (Suppl. Fig. 1), also performed better compared to aCompCor (pipelines 14 vs 6 in Fig. 8). In the present study, however, we also examined variants of aCompCor50 that consisted of GSR and WM denoising with different thresholds of variance for choosing the optimal number of regressors (pipelines 15-19). In the case of the Gordon and Seitzman atlas, GSR combined with WM regressors needed to explain about 45% variance performed almost as well as 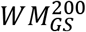, whereas for the MIST atlas, GSR with WM regressors needed to explain 50% variance performed slightly better than 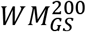 (Fig. 8). It should be noted that, in the case that the number of WM regressors was determined based on the variance accounted for, the exact number of regressors, and equivalently the loss of degrees of freedom from the data, varied across participants. However, despite the varying number of degrees of freedom in the data, this approach performed, overall, as well as 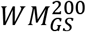, both in terms of enhancement of SNR and mitigation of motion artifacts and biases. Based on these observations, we believe that it will be important for future studies to examine more systematically whether choosing the number of regressors based on a specific percentage of variance accounted for can lead to improved data quality for a variety of fMRI datasets that may differ in terms of pulse sequence (single-band vs multi-band fMRI, scan duration, main magnetic field, etc.) or population examined.

Another potential means of improving the performance of aCompCor is by defining the WM and CSF masks in a way that partial volume effects are minimized, such as by restricting the CSF mask to the lateral ventricles or by applying erosion along the boundaries, as was done in previous studies (Behzadi et al., 2007; Muschelli et al., 2014). It should be noted that in the present study, the probability thresholds for defining the WM and CSF compartments (0.80 and 0.90, respectively) were determined based on visual inspection. However, based on a secondary analysis, the performance of CSF denoising can be improved if a stricter probability threshold is considered, albeit it still cannot outperform WM denoising (Suppl. Fig. 21). Given this finding, we believe that future work should also examine the role of the probability threshold for defining the WM and CSF masks in the context of denoising, as well as whether the optimal threshold depends on factors such as the spatial resolution of fMRI data and the software used for tissue segmentation.

### 4.3 GSR combined with WM or FIX denoising further improves SNR and mitigates motion effects

The GS, which is defined as the average fMRI time series across all voxels in the brain or GM, is often regressed out from the data. In our study, GSR improved the scores for the signal-related QC metrics and, to a less extent, the scores for the motion-related QC metrics for both WM and FIX denoising (Fig. 2-Fig. 3). For a low number of PCA regressors in WM denoising, we observed that the effect of GSR was stronger compared to higher regressor numbers, which implies that WM regressors share common variance with the GS. Previous studies have shown that the GS derived either by the whole brain or GM are very similar to each other and also that the GS is highly correlated with the mean time series across voxels in WM and CSF (Kassinopoulos and Mitsis, 2019; Power et al., 2017), which further lends support to the idea that WM regressors share common variance with the GS. Furthermore, we observed that the SLFOs that reflect BOLD fluctuations due to changes in heart rate and breathing patterns, and account for a significant fraction of GS fluctuations (Falahpour et al., 2013; Kassinopoulos and Mitsis, 2021, 2019), were well explained using the first 20-30 WM and CSF regressors (Fig. 1). This result suggests that the practice of considering PCA regressors from WM or CSF exhibits to some extent similar effects to GSR. As a result, the effect of GSR when considering 200 WM regressors (i.e., 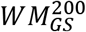 vs *WM*^200^) is relatively small (Fig. 2-Fig. 3). In contrast, GSR has a strong effect on FIX denoising, which suggests that the ICA regressors that are removed by FIX denoising do not share a large fraction of common variance with the GS. This is not surprising, as it has been suggested that spatial ICA used in FIX is, by design, unable to separate global temporal artifacts from fMRI data (Glasser et al., 2018). For instance, it has been shown that the default mode network identified with ICA is often confounded by variations in breathing patterns (Birn et al., 2008a). And since the BOLD fluctuations due to variations in breathing patterns or heart rate are in low frequencies (~0.1 Hz), FIX is unlikely to classify components that are confounded by these sources as noisy with the result of preserving global artifacts in the fMRI data.

Despite the simplicity of GSR, there has been much debate about its use (Liu et al., 2017; Murphy and Fox, 2017). Even though several studies have shown that a large fraction of the GS is associated to physiological processes such as heart rate and breathing activity (Birn et al., 2006; Chang et al., 2009; Falahpour et al., 2013; Kassinopoulos and Mitsis, 2021, 2019; Shmueli et al., 2007; Wise et al., 2004) as well as head motion (Power et al., 2014; Satterthwaite et al., 2013), there is accumulating evidence that GS is also driven by neuronal activity as assessed by intracranial recordings (Schölvinck et al., 2010) and vigilance-related measures (Chang et al., 2016; Falahpour et al., 2018; Liu and Falahpour, 2020; Wong et al., 2016, 2013). Therefore, while our results are in support of GSR for both WM and FIX denoising, we cannot exclude the possibility of removing some neuronal-related fluctuations from the data when the GS is removed. Finally, we should acknowledge that several attempts have been made to address some of the limitations of GSR (Aquino et al., 2020; Carbonell et al., 2014, 2011; Erdoğan et al., 2016); however, whether the proposed techniques suppress physiological noise while also preserving neuronal activity still remains an open question.

## 5. Conclusion

In summary, the current study evaluated the performance of a large range of model-based and model-free techniques using previously proposed as well as novel QC metrics. As the QC metrics did not uniformly favor a specific preprocessing strategy, we proposed a framework that evaluates the sensitivity of each metric. Among eight QC metrics, *FCC* proposed here as well as *FDDVARS* employed in Muschelli et al. (2014) exhibited the highest sensitivity. *FCC* reflects the difference between WNE and BNE correlation values, whereas *FDDVARS* reflects the levels of motion artifacts in the parcel time series. Our results suggest that the choice of the preprocessing strategy to be employed in a study could be based solely on the metrics *FCC* and *FDDVARS*, where the former tends to favor relatively mild strategies for improving the identification of large-scale networks, while the latter is in favor of more aggressive strategies in an attempt to minimize the presence of motion artifacts.

The data-driven approaches (WM denoising and FIX denoising) combined with GSR demonstrated the largest increase in SNR as well as reduction in motion artifacts and biases. In the case of WM denoising, using resting-state fMRI data from the HCP, we found that removing about 17% of the WM regressors yielded one of the largest improvements in QC scores. Scrubbing did not provide any gain to the data quality when it was followed by WM denoising, whereas low-pass filtering at 0.2 Hz led to an additional improvement, particularly when combined with mild variants of WM denoising.

Similar conclusions were derived using three different functional atlases. However, unless the framework followed here is repeated with different datasets that vary in terms of the population examined or acquisition parameters (e.g. repetition time TR and duration of scan) we cannot be certain whether our conclusions can be directly generalized to other datasets. Therefore, we recommend investigators to consult the QC metrics when deciding about the pipeline they want to employ in a study. Finally, as has been suggested in previous studies (Ciric et al., 2017; Parkes et al., 2018), we recommend investigators to report scores of QC metrics for the preprocessed data so that readers can independently interpret the findings with respect to possible biases that can arise due to motion. To assist with this, we provide the codes used in this study (https://github.com/mkassinopoulos/Estimation_of_QC_metrics), which can be used for preprocessing of the data and estimation of the QC scores.

## Acknowledgments

This work was supported by the Natural Sciences and Engineering Research Council of Canada (Discovery Grant 34362 awarded to GDM), the Fonds de la Recherche du Quebec - Nature et Technologies (FRQNT; Team Grant PR191780-2016 awarded to GDM) and the Canada First Research Excellence Fund (awarded to McGill University for the Healthy Brains for Healthy Lives initiative). MK acknowledges funding from Québec Bio-imaging Network (QBIN). Data were provided by the Human Connectome Project, WU-Minn Consortium (Principal Investigators: David Van Essen and Kamil Ugurbil; 1U54MH091657) funded by the 16 NIH Institutes and Centers that support the NIH Blueprint for Neuroscience Research; and by the McDonnell Center for Systems Neuroscience at Washington University. The authors would like to thank Alba Xifra-Porxas for her assistance in the selection of subjects from the HCP database with good quality physiological recordings.

## Author contributions

Michalis Kassinopoulos preprocessed and analyzed the data, designed the figures, interpreted the results, drafted the manuscript and arranged the final version. Georgios D. Mitsis contributed to the conceptualization of the research, supervised the findings, and edited and reviewed the manuscripts.

## Data and Code Availability Statement

For the purposes of this study we used resting-state scans from the Human Connectome Project (HCP) S1200 release (Matthew F Glasser et al., 2016; Van Essen et al., 2013) which are publicly available and can be found at https://www.humanconnectome.org/. Furthermore, the codes written for this work are publicly available and can be found at https://github.com/mkassinopoulos/Estimation_of_QC_metrics.

## Supplementary Material

**Suppl. Fig. 1.**
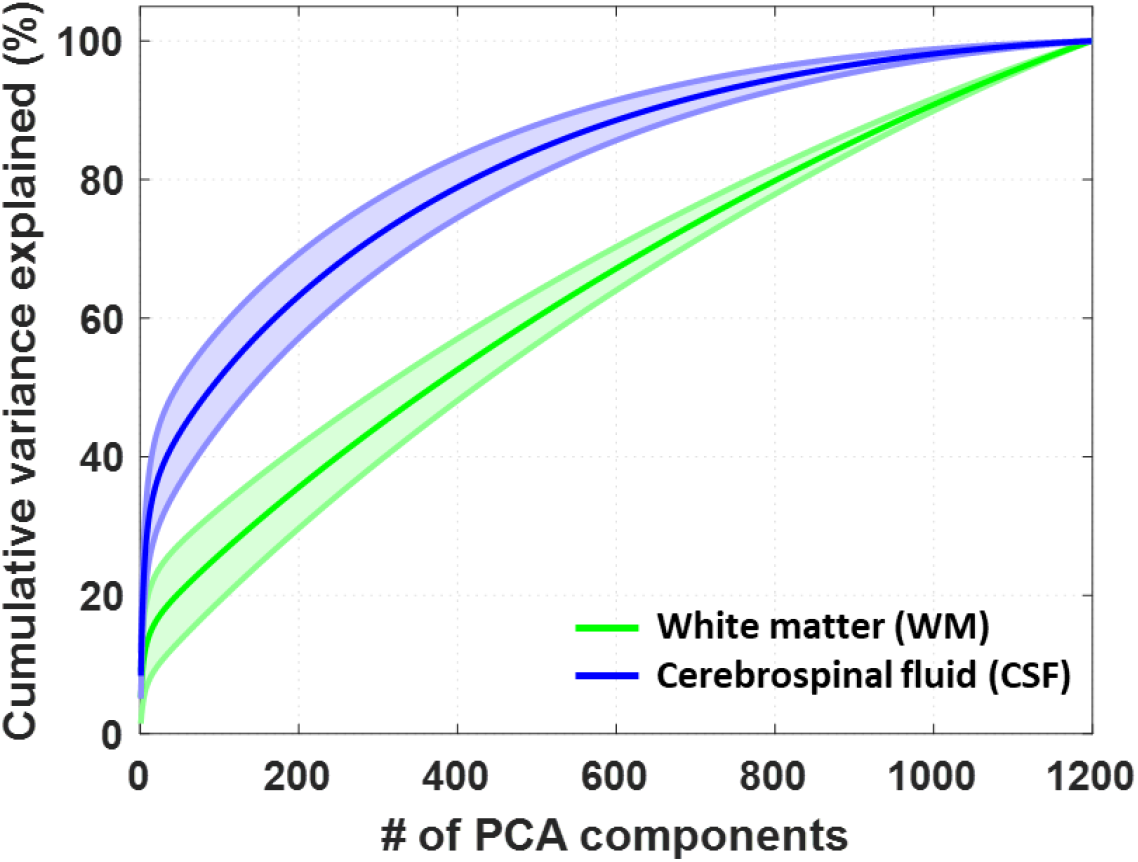
Cumulative variance explained (%) for PCA regressors obtained from WM and CSF. About 360 (±60) WM and 90 (±40) CSF regressors were required to explain 50% of the variance in the WM and CSF compartments, respectively. These numbers correspond to the number of PCA regressors to be removed from the fMRI data according to the variant of aCompCor proposed in Muschelli et al. (2014). The first 200 WM regressors used in *WM*^200^ correspond to an average explained variance of 36±6 %.

**Suppl. Fig. 2.**
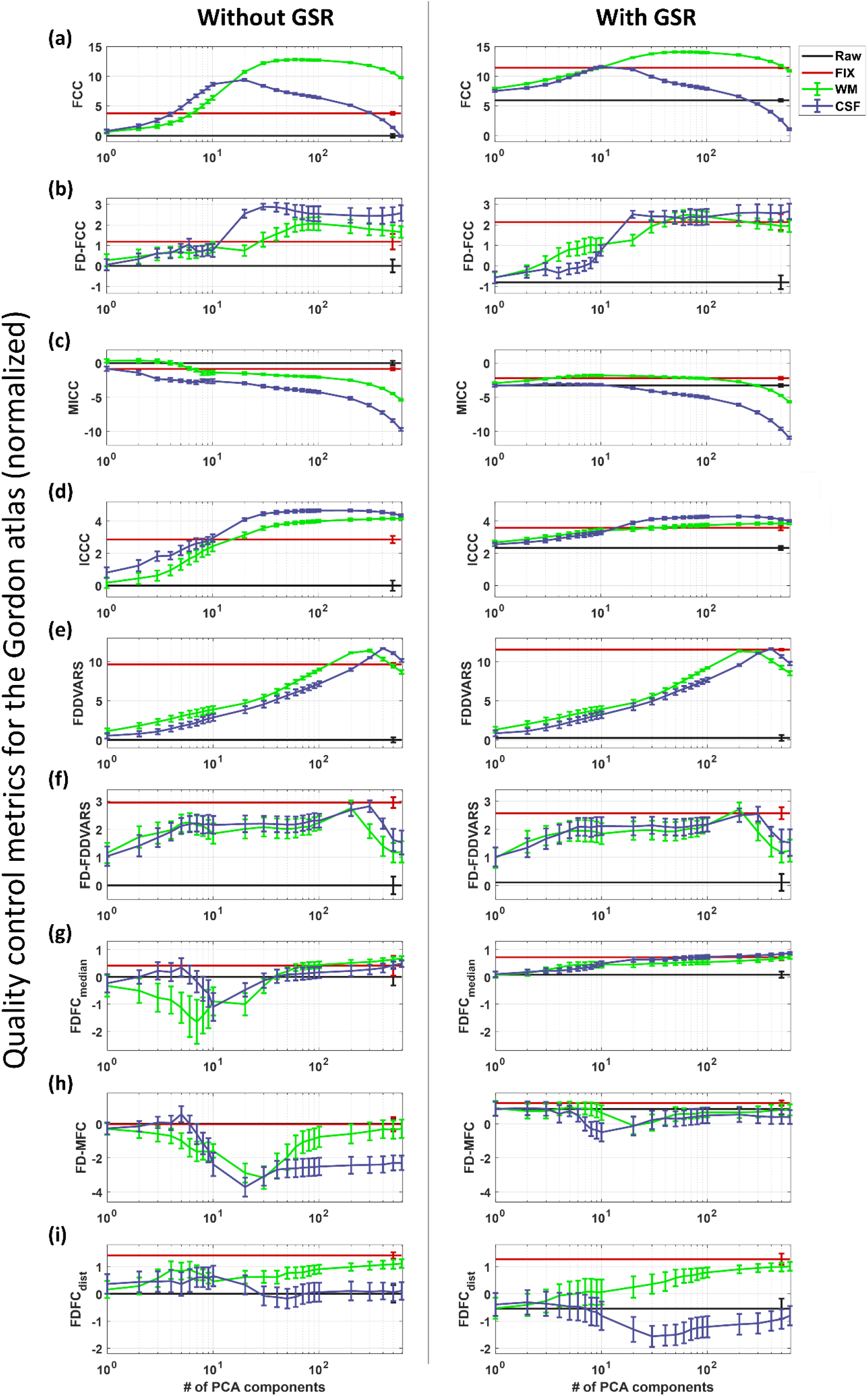
Quality control (QC) scores obtained by aCompCor, after normalization, using the data in the Gordon parcel space. To summarize all the QC metrics to signal-related and motion-related metrics it was important that the obtained scores from each group of subjects were transformed to *Z*-scores as described in Section 2.7. For both signal- and motion-related metrics, high normalized scores corresponded to better quality of data. Among all metrics, *FCC* and *FDDVARS* demonstrated the most significant improvement in quality with respect to the raw data. For the correspondence of the different curves and lines please see Fig. 2.

**Suppl. Fig. 3.**
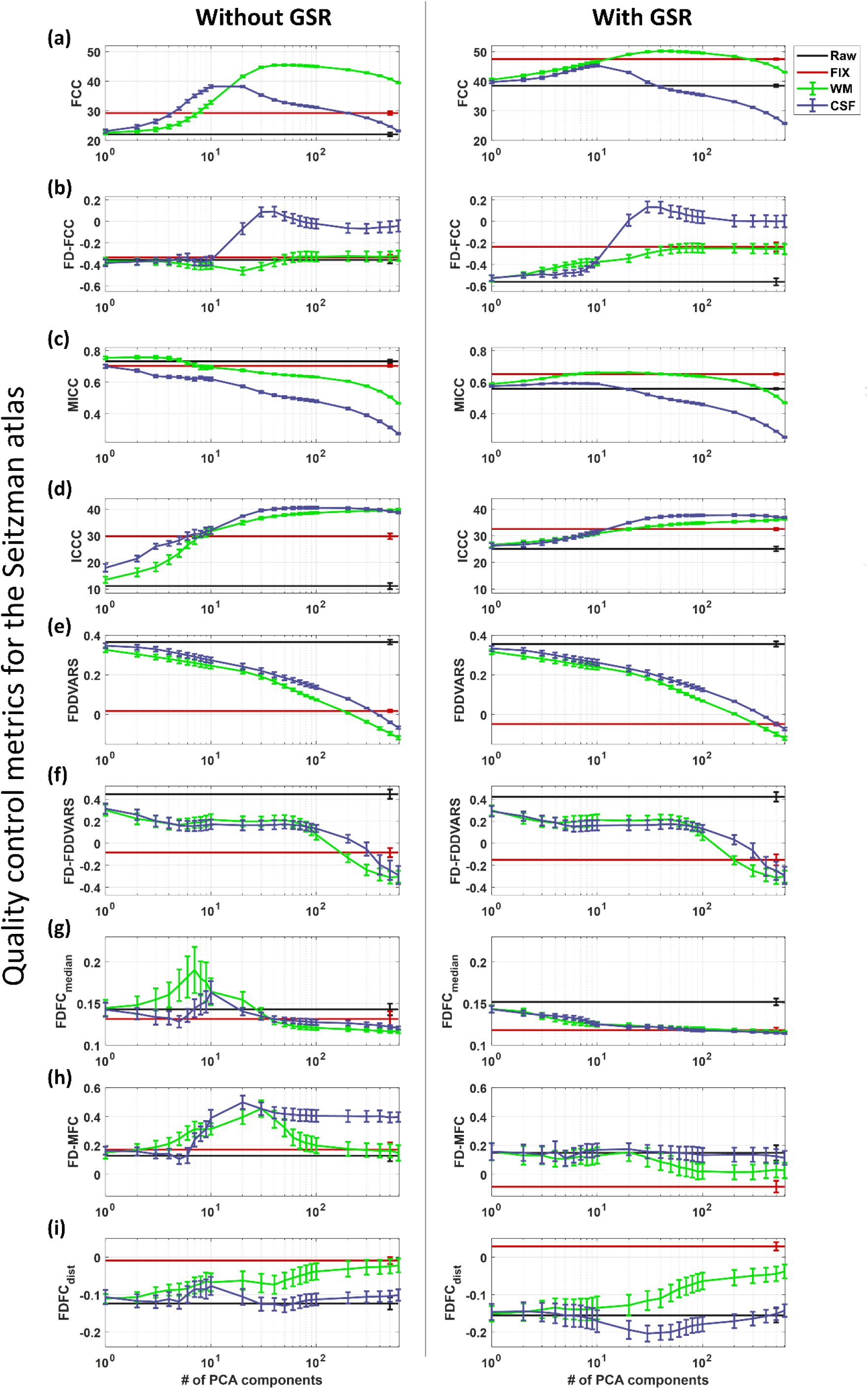
Quality control (QC) scores obtained by aCompCor using the data in the Seitzman parcel space. Similarly to the data at the Gordon parcel space, different trends were observed among the nine QC metrics. Furthermore, the two QC scores *FCC* and *ICCC* that are based on the contrast in the FC and *ICC* matrices exhibited different range of scores for the data in the Seitzman parcel space compared to the data in the Gordon parcel space which may be related to the different number of parcels and networks between the atlases. However, the trends of these two scores for varying number of components were similar. For the correspondence of the different curves and lines please see Fig. 2.

**Suppl. Fig. 4.**
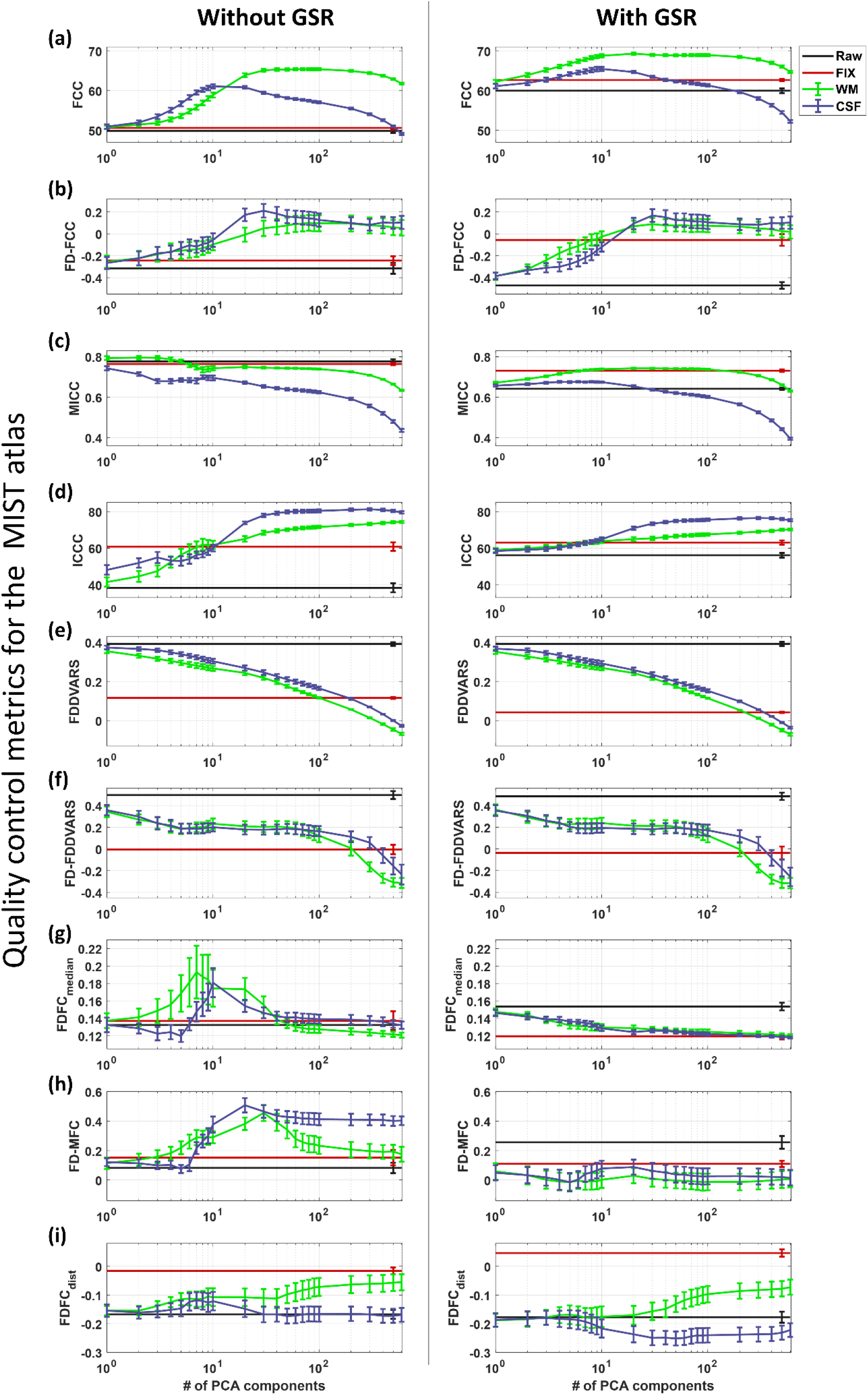
Quality control (QC) scores obtained by aCompCor using the data in the MIST parcel space. Similarly to the data at the Gordon parcel space, different trends were observed among the nine QC metrics. Furthermore, the two QC scores *FCC* and *ICCC* that are based on the contrast in the FC and *ICC* matrices exhibited different range of scores for the data in the MIST parcel space compared to the data in the Gordon parcel space which may be related to the different number of parcels and networks between the atlases. However, the trends of these two scores for varying number of components were similar. For the correspondence of the different curves and lines please see Fig. 2.

**Suppl. Fig. 5.**
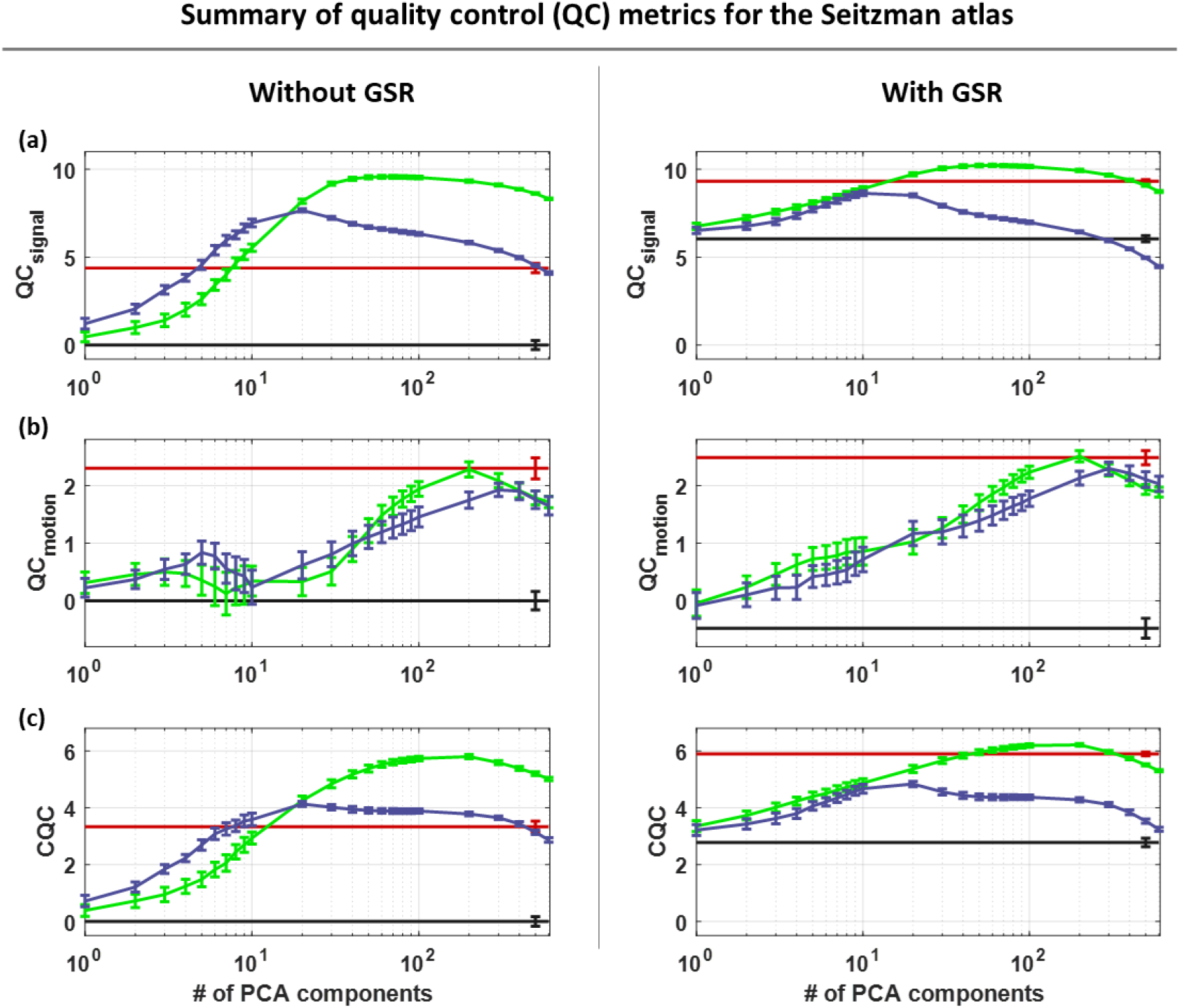
Summarized QC scores obtained by aCompCor using the data in the Seitzman parcel space. Similarly to the data in the Gordon parcel space (Fig. 3), GSR and white matter denoising with 50 to 100 PCA regressors yielded the highest scores for *QC*_signal_ whereas the more aggressive set of regressors 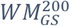 achieved the highest score in *QC*_motion_. Overall, the CQC score that accounts for both *C*_signal_ and *QC*_motion_ yielded its highest value when the set 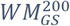 was used.

**Suppl. Fig. 6.**
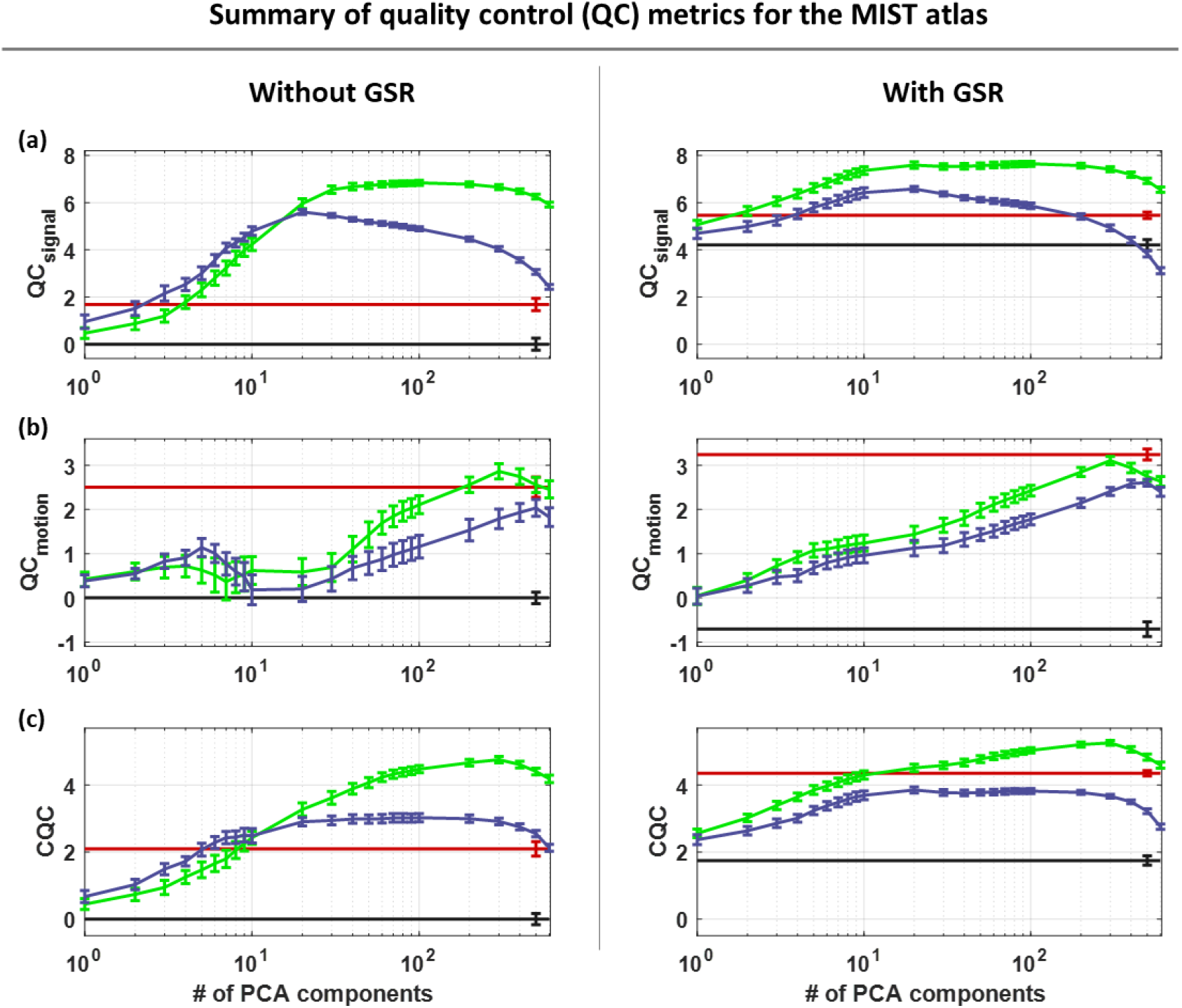
Summarized QC scores obtained by aCompCor using the data in the MIST parcel space. In the MIST parcel space, the *QC*_signal_ was kept relatively stable at its highest score for a larger range of sets 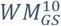 - 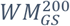 compared to the *QC*_signal_ in the Gordon and Seitzman parcel space. Moreover, the *QC*_motion_ yielded its highest score for the set 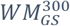. As a result, the set 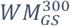 exhibited the highest score for the *CQC* as well.

**Suppl. Fig. 7.**
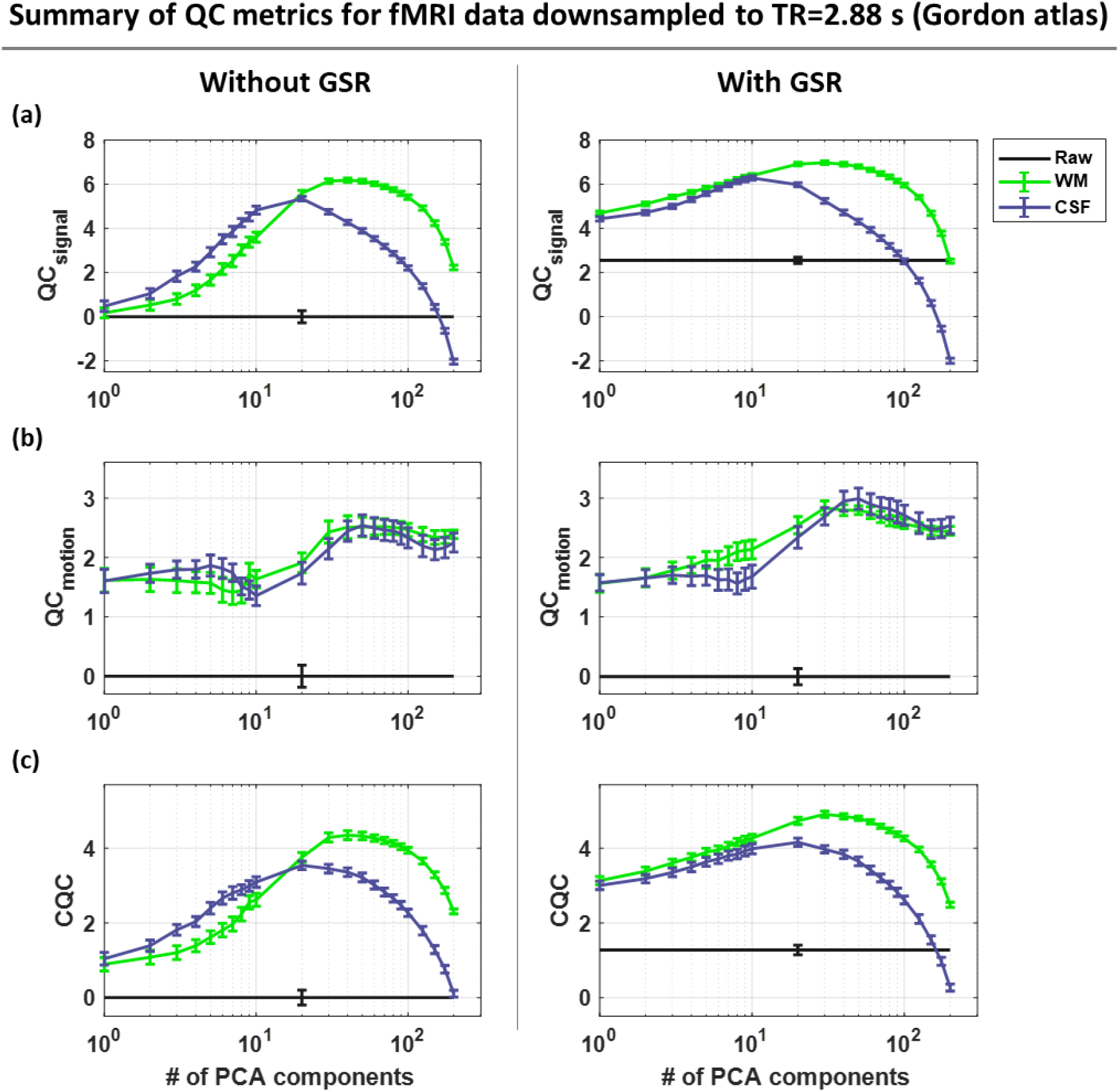
Summarized QC scores obtained from the fMRI data after being downsampled to TR=2.88 s (Gordon atlas). About 20 to 40 PCA regressors from WM were needed in order to achieve high *QC*_*signal*_ scores, while 50 regressors from either WM or CSF yielded the highest score in *QC*_*motion*_. Among all sets of regressors, 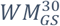 yielded the highest score (4.9) for the CQC metric.

**Suppl. Fig. 8.**
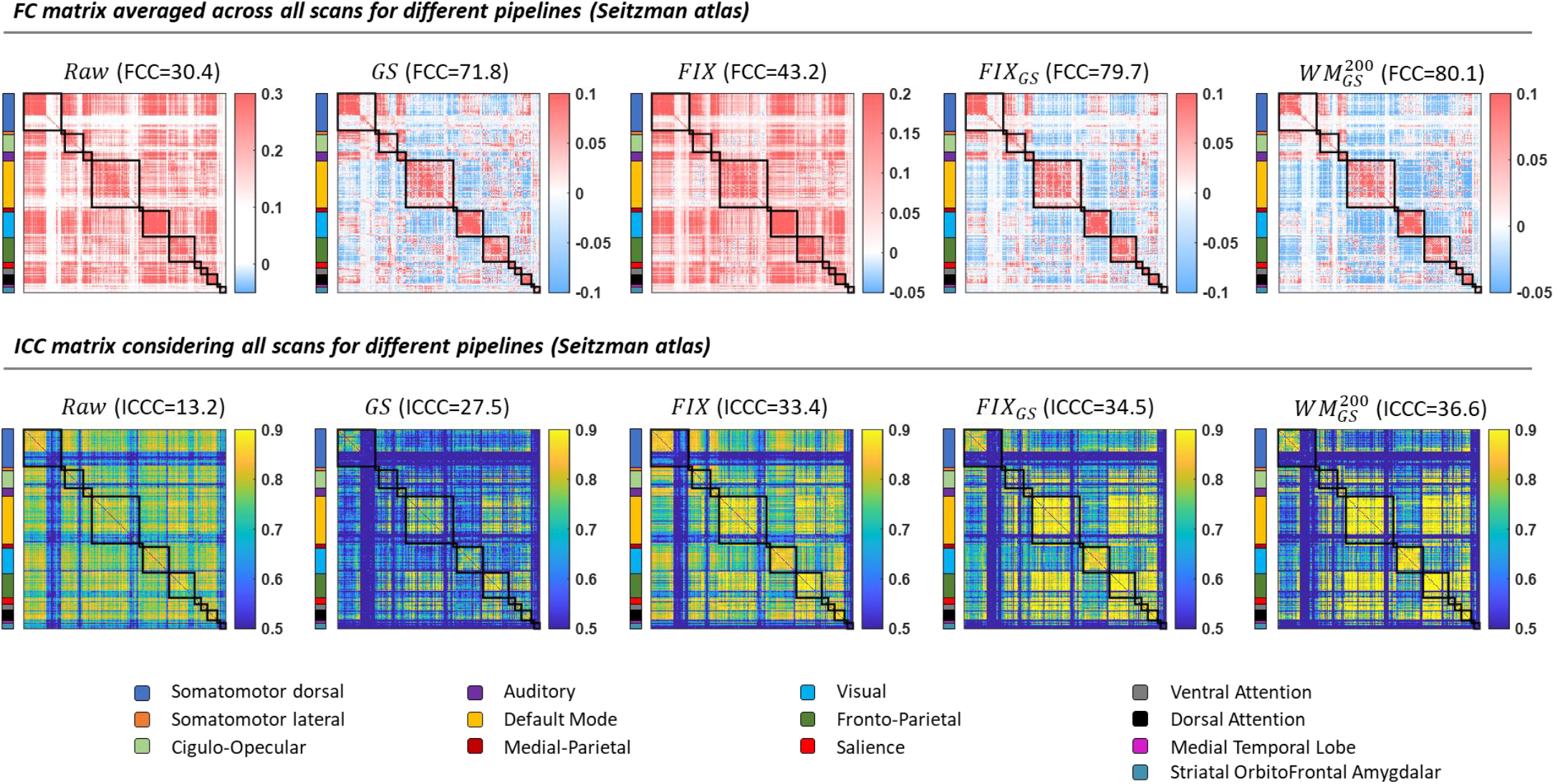
FC (top) and ICC (bottom) matrices considering all scans for different pipelines obtained from the data in the Seitzman parcel space. The pipelines 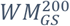 and *FIX*_*GS*_ significantly improved the identifiability of the networks. Note that many parcels appeared at the end of each network illustrated low correlation and *ICC* values. These parcels correspond to subcortical parcels and as reported by Seitzman et al. (2020), those parcels demonstrated low temporal signal-to-noise (SNR) in the HCP data which may explain the low correlation and *ICC* values observed here. Similar to the data in the Gordon parcel space, a large number of BNEs, and especially edges corresponding to interactions between the default mode and fronto-parietal networks, exhibited low FC values but high *ICC* values.

**Suppl. Fig. 9.**
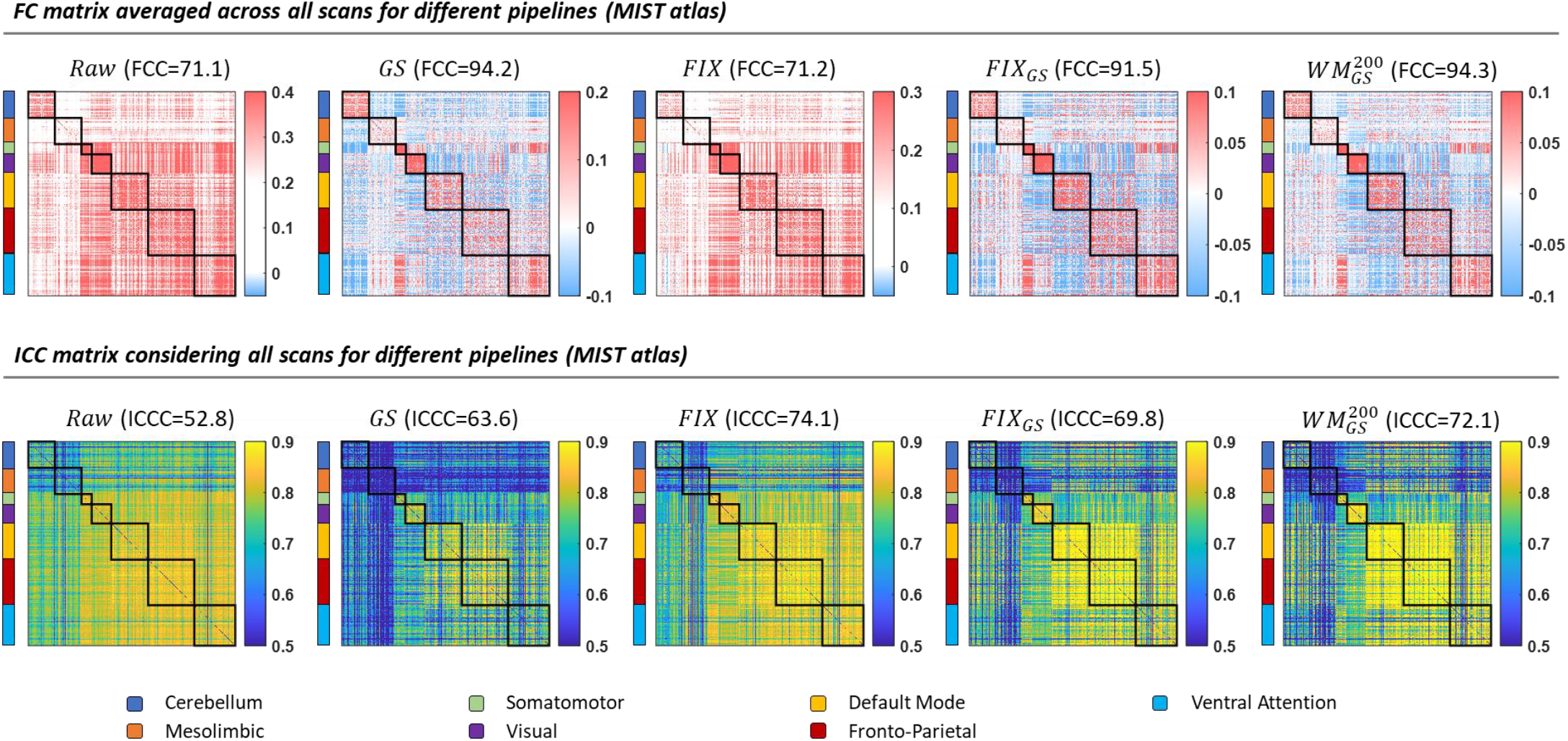
FC (top) and *ICC* (bottom) matrices considering all scans for different pipelines obtained from the data in the MIST parcel space. Based on the *FCC* scores obtained from the group-level FC matrices, the pipelines 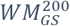 and *FIX*_*GS*_ significantly improved the identifiability of the networks. However, similar *FCC* score was observed when only the GS was regressed out. On the other hand, as seen in Suppl. Fig. 3a, on a scan-specific basis analysis, the *FCC* scores between the pipelines *GS*, *FIX*_*GS*_ and 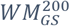 presented significant differences. Moreover, we observe that, similar to the data in the Gordon parcel space, a large number of BNEs, and especially edges corresponding to interactions between the default mode and fronto-parietal networks, exhibited low FC values but high *ICC* values.

**Suppl. Fig. 10.**
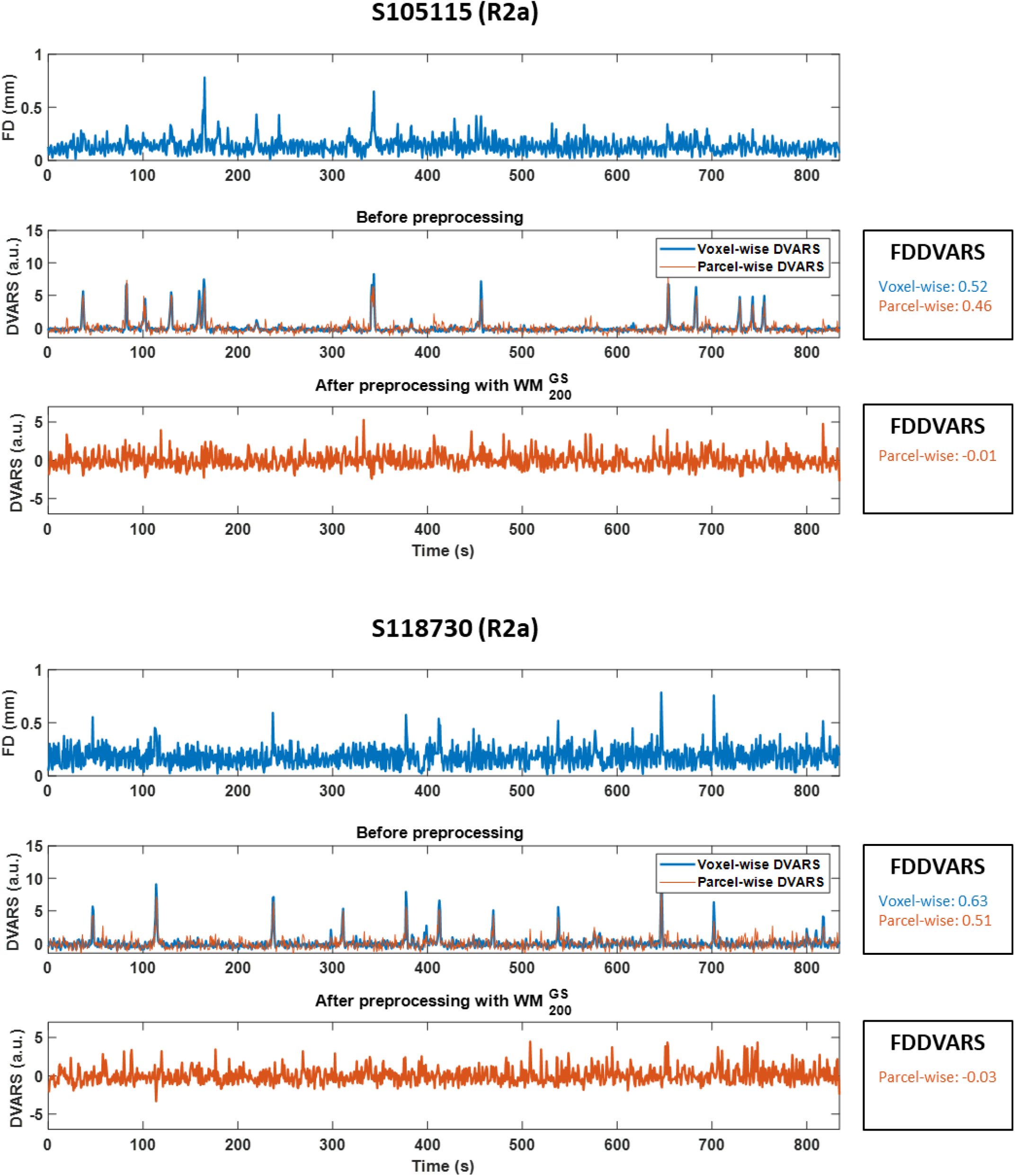
Traces of *DVARS* time-series for two scans for which WM denoising 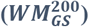 reduced the corresponding *FDDVARS* values (i.e. correlation of *FD* and *DVARS*) close to zero. For the raw data, *DVARS*, either estimated at the voxel- or parcel-level, exhibited frequent spikes at times of motion, as assessed with *FD*, which resulted in *FDDVARS* values above 0.45. Parcel-wise *FDDVARS* values were on average lower than voxel-wise *FDDVARS* (0.37±0.19 vs 0.53±0.20). After WM denoising, the spikes in *DVARS* were significantly attenuated, resulting in *FDDVARS* values close to zero (0.02±0.06).

**Suppl. Fig. 11.**
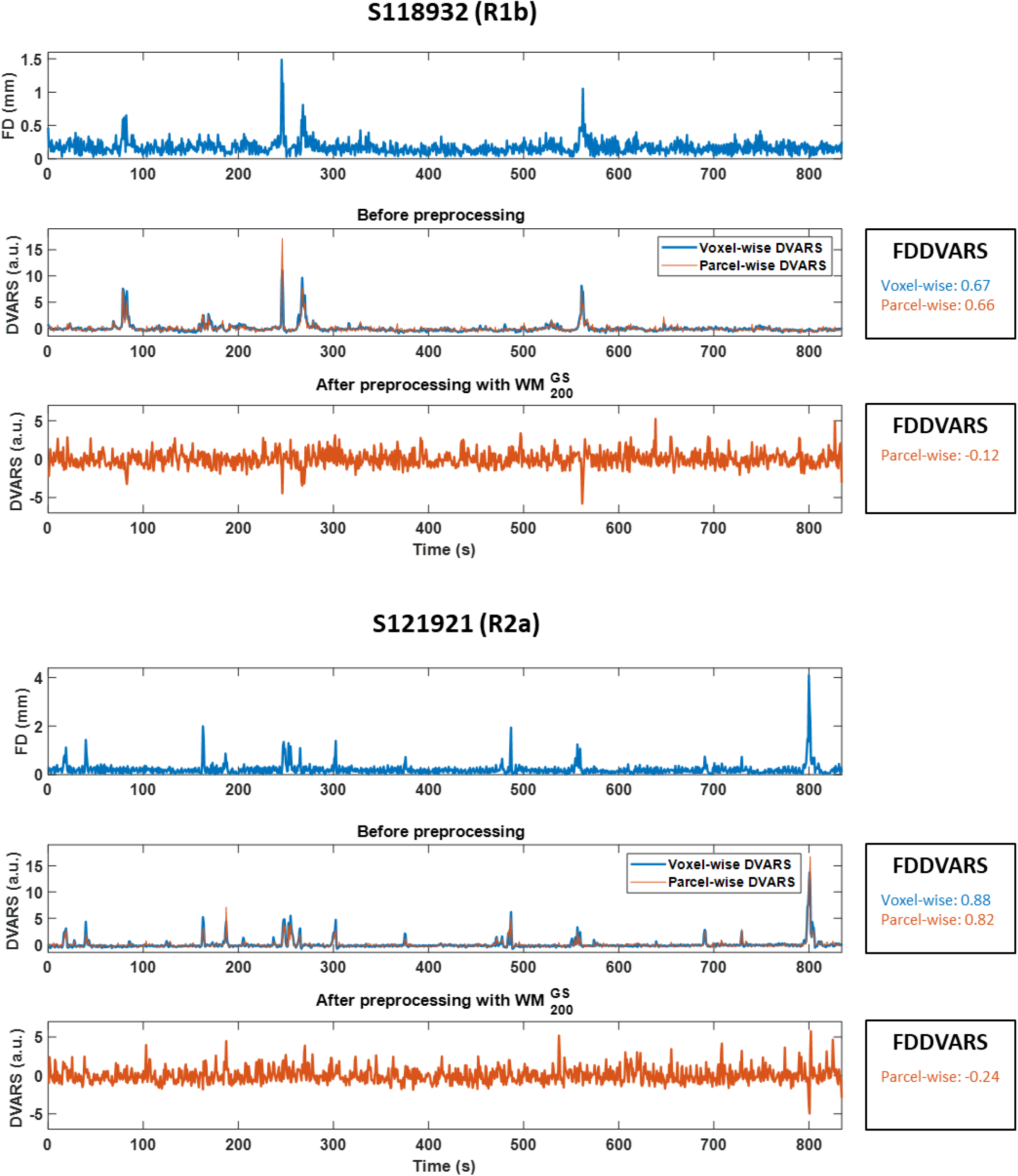
Traces of *DVARS* time-series for two scans for which WM denoising 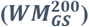 resulted in negative *FDDVARS* values. In some scans, while most of the spikes in *DVARS* were significantly attenuated after WM denoising, the sign of some spikes reversed from positive to negative which resulted in negative *FDDVARS* values (e.g. spikes at 670 s and 800 s for scans S118932 (R1b) and S121921 (R2a), respectively). As suggested by Power et al. (2020), these *DVARS* dips indicate that the framewise changes in fMRI time series are abnormally low at those times. As negative *FDDVARS* values may be associated with the presence of *DVARS* dips at times with motion, and these dips can presumably lead to systematic confounds, *FDDVARS* values, similar to the rest of the motion-related QC metrics, were normalized to 1 − abs(*FDDVARS*) so that a high positive value of normalized *FDDVARS* is assigned to good quality data.

**Suppl. Fig. 12.**
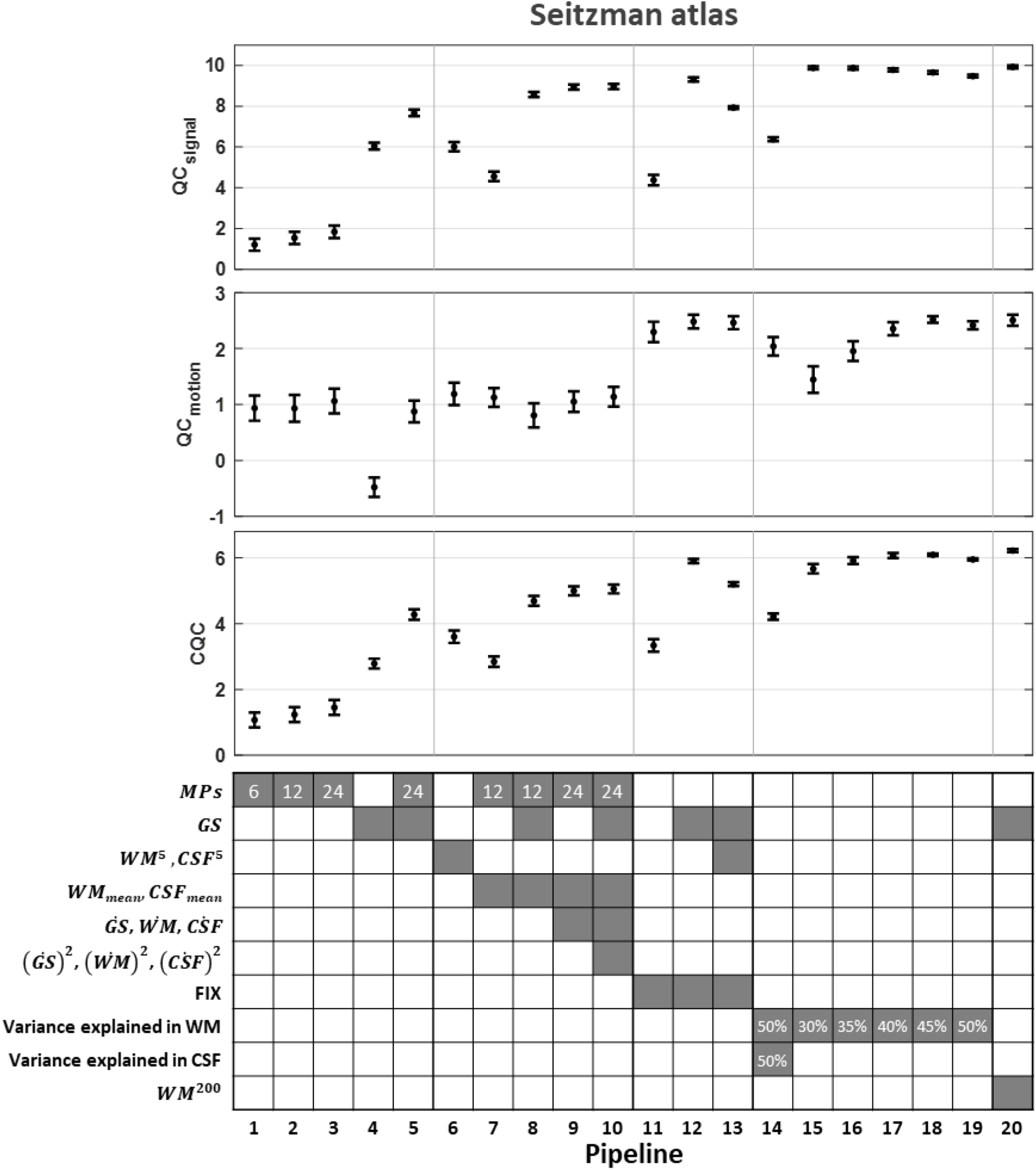
Evaluation of data-driven NCTs (Seitzman atlas). Twenty different data-driven pipelines were examined, as listed in Table 1. Similar to the Gordon atlas, among all pipelines, pipelines that included GSR and WM or FIX denoising yielded the highest scores in *QC*_signal_, *QC*_motion_ and *CQC* (i.e., pipelines 12 and 17-20).

**Suppl. Fig. 13.**
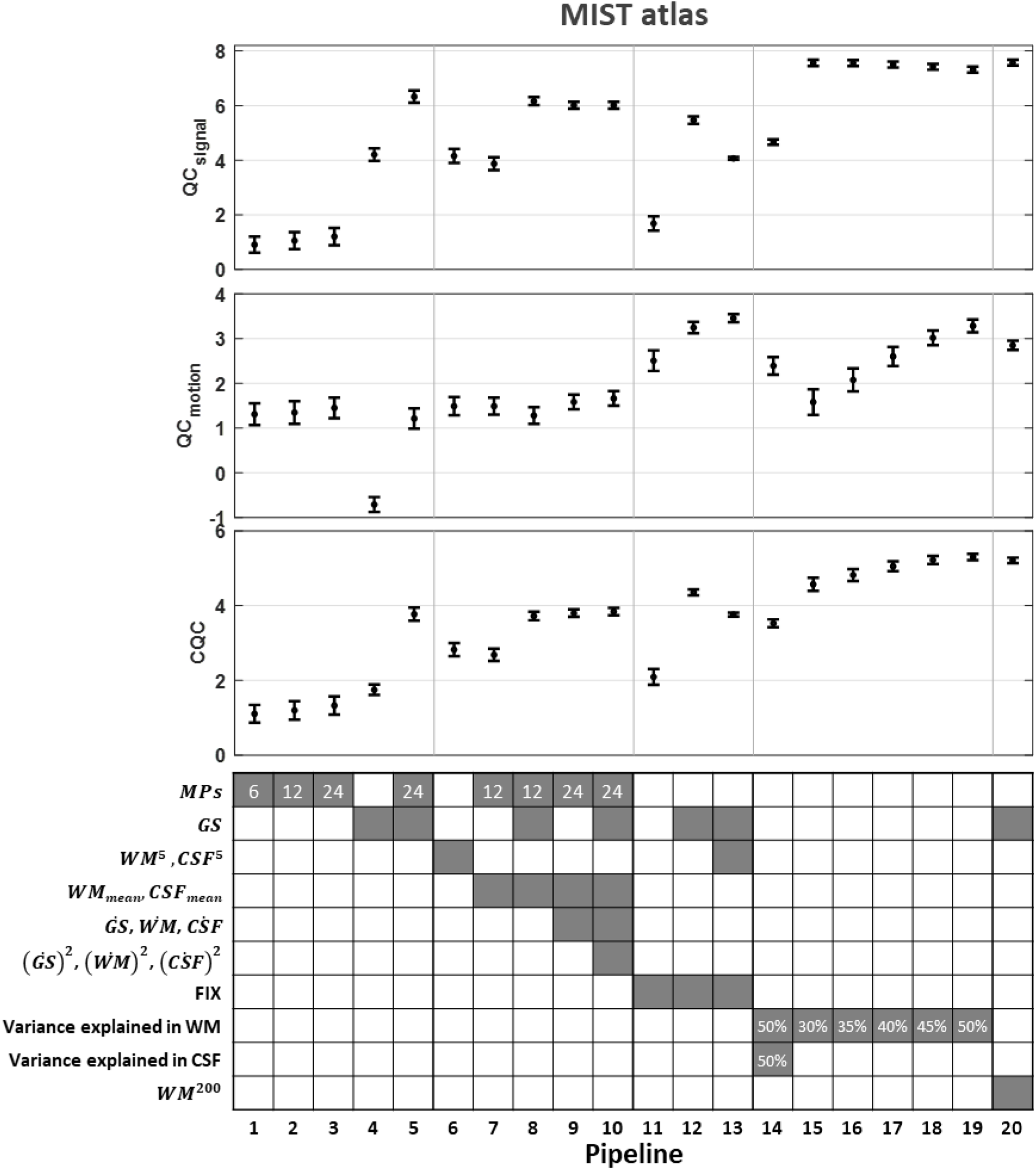
Evaluation of data-driven NCTs (MIST atlas). Twenty different data-driven pipelines were examined, as listed in Table 1. Similar to the Gordon atlas, among all pipelines, pipelines that included GSR and WM denoising yielded the highest scores in *QC*_signal_, *QC*_motion_ and *CQC* (i.e., pipelines 17-20). The FIX denoising (pipeline 12) yielded a smaller improvement in *QC*_signal_ compared to the Gordon and Seitzman atlases.

**Suppl. Fig. 14.**
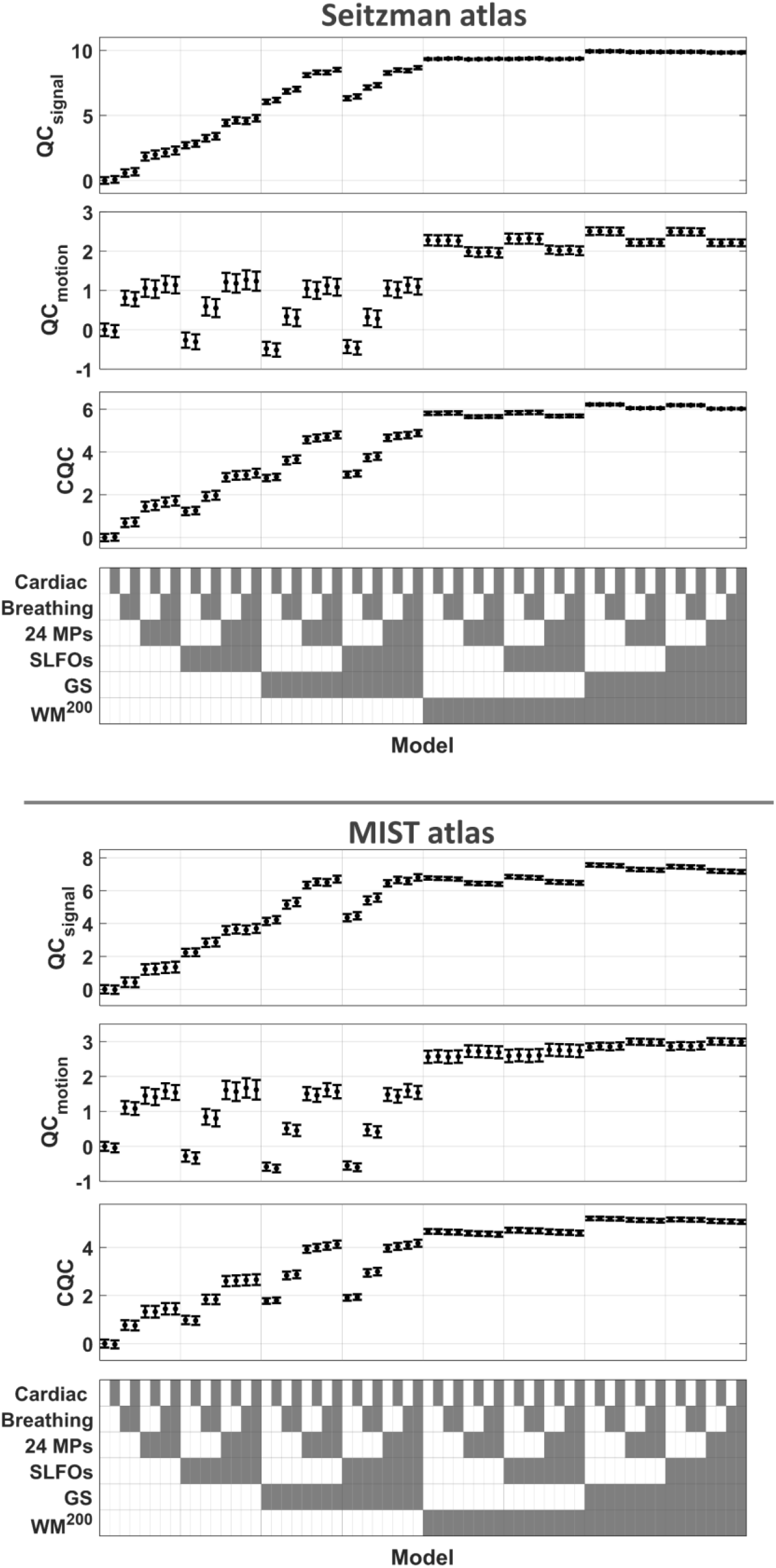
Evaluation of model-based NCTs using the fMRI data in the Seitzman (top) and MIST (bottom) parcel space. As with the data in the Gordon parcel space (Fig. 9), when the model-based regressors related to SLFOs, head motion and breathing motion were used without tissue-based regressors, the data quality as assessed with the three QC metrics *QC*_signal_, *QC*_motion_ and *CQC*, was improved. However, when the set of nuisance regressors included the GS and the 200 components from WM, none of the model-based regressors was found to provide any additional improvement on the data quality.

**Suppl. Fig. 15.**
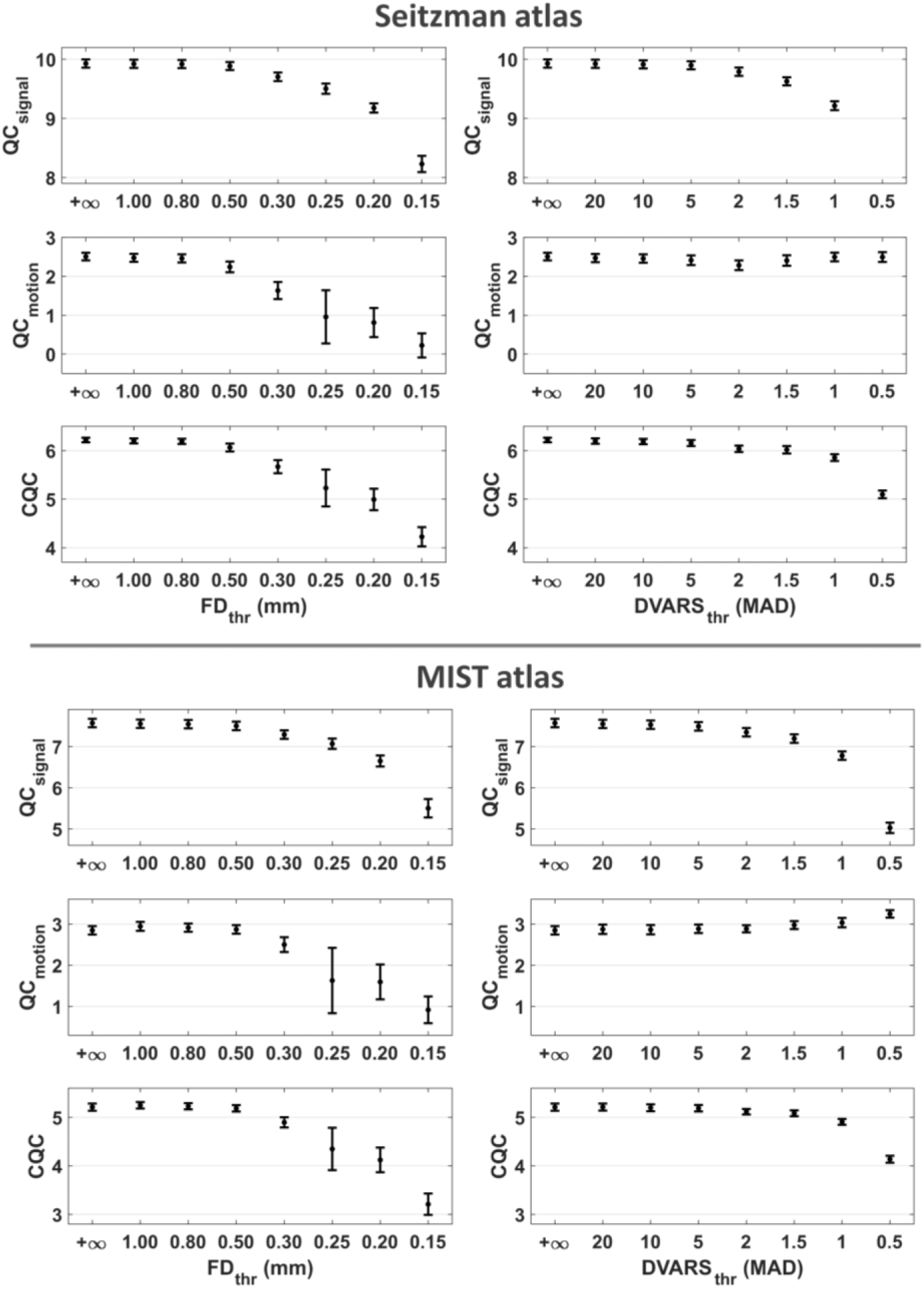
Effect of scrubbing in data quality for different threshold values on fMRI data in the Seitzman (top) and MIST (bottom) parcel space. When the data were preprocessed with the set of regressors 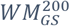, scrubbing before the removal of the regressors did not provide any improvement in the combined summarized QC metric *CQC*. In contrast, thresholds of *FD*_thr_ below 0.50 mm led to a significant decrease of the *CQC* score. In the case of the *DVARS*, the *CQC* score was decreased when the threshold was below 1.5 MAD. However, typically, higher values of *DVARS*thr are used in the literature.

**Suppl. Fig. 16.**
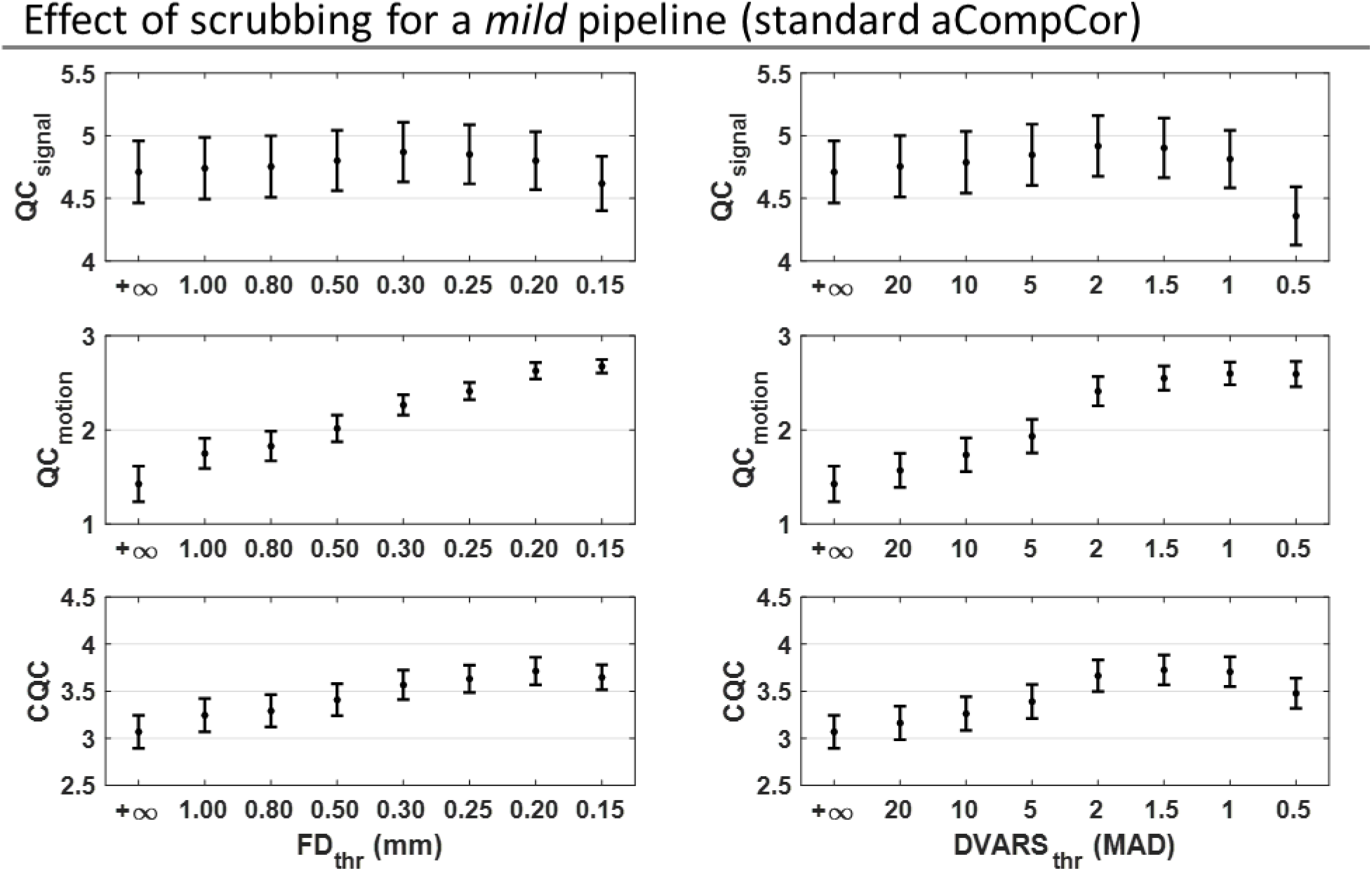
Effect of scrubbing for a mild pipeline that consists of removing five WM and five CSF regressors (Gordon atlas). When the data were preprocessed with the standard aCompCor approach (i.e. removal of five WM and CSF regressors), scrubbing before the removal of the regressors provided an additional improvement in the combined summarized QC metric *CQC*. The largest improvement was observed for a threshold *FD*_thr_ equal to 0.20 mm, which is commonly used in the literature (Power et al., 2015) and for a threshold *DVARS*_thr_ equal to 1.5 MAD.

**Suppl. Fig. 17.**
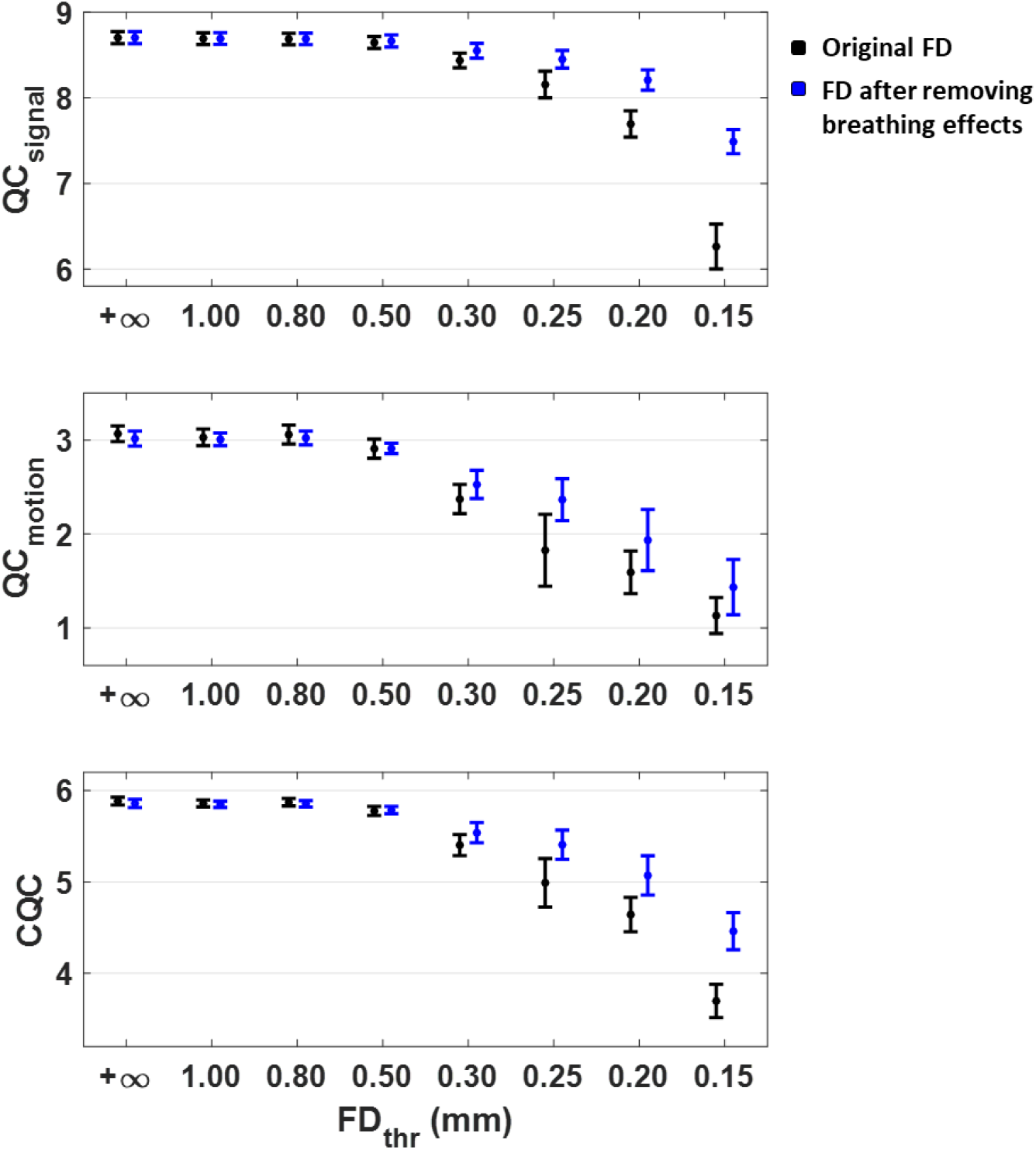
Effects of scrubbing for an *FD* trace that is free of breathing motion effects (Gordon atlas). To remove the effects of breathing motion from the *FD* trace, the six motion realignment parameters that are used in the estimation of *FD* were first filtered out for breathing-related fluctuations using 3rd order RETROICOR. When the data were preprocessed using 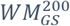, scrubbing resulted in lower QC metric values, regardless of whether the influence of breathing in *FD* traces was removed or not. Removing the influence of breathing motion from *FD* traces was found to attenuate the effects of scrubbing, which is likely due to the smaller number of volumes marked as motion-contaminated for a given value of threshold *FD*_thr_.

**Suppl. Fig. 18.**
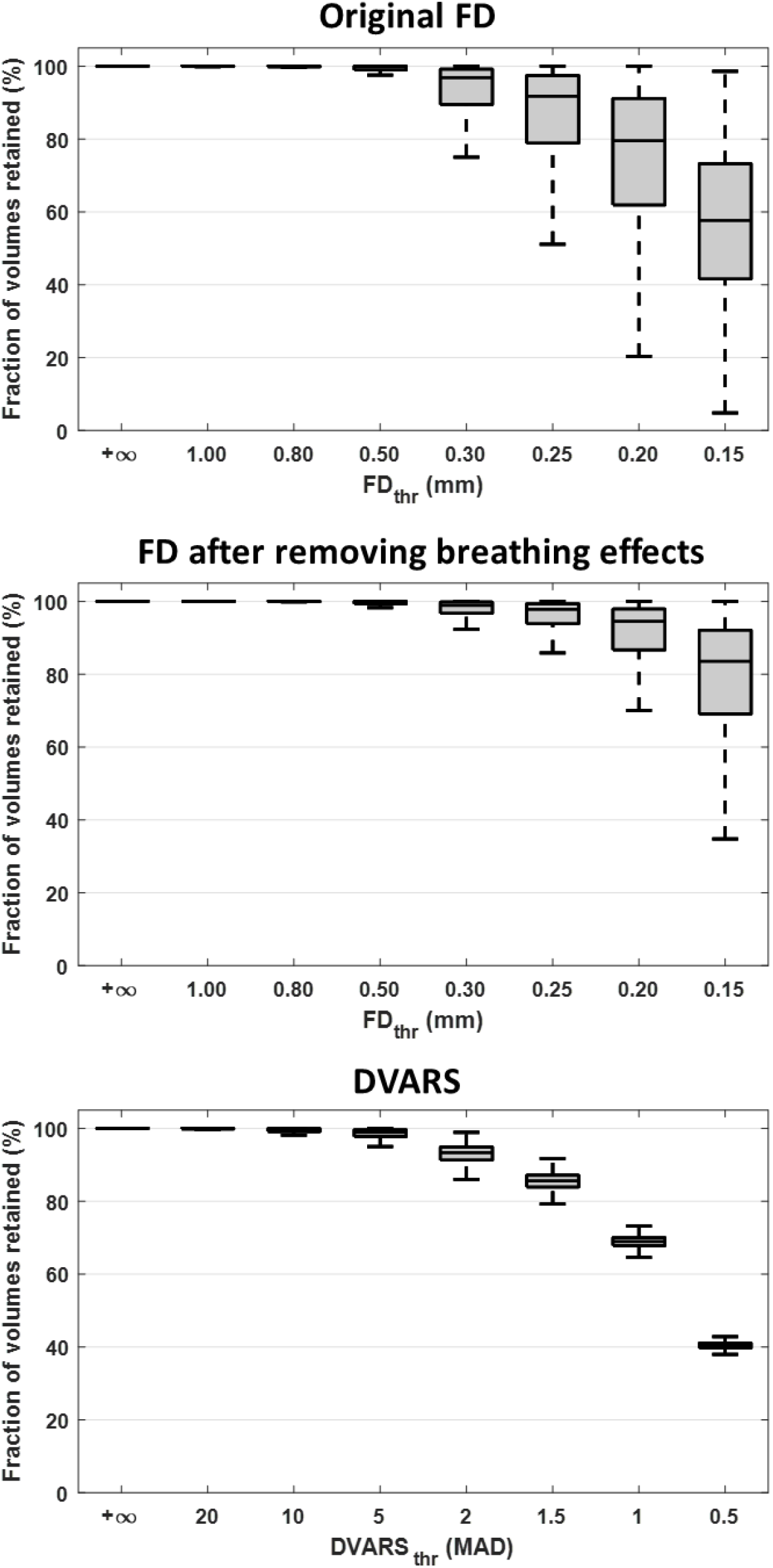
Fraction of volumes retained after scrubbing based on the original *FD* trace (top row), an *FD* trace free of breathing motion effects (middle row) and *DVARS* (bottom row). The bottom and top of each box correspond to the 25th and 75th percentiles of the sample distribution and the line in the box corresponds to the median (the outliers are not shown for ease of visualization).

**Suppl. Fig. 19.**
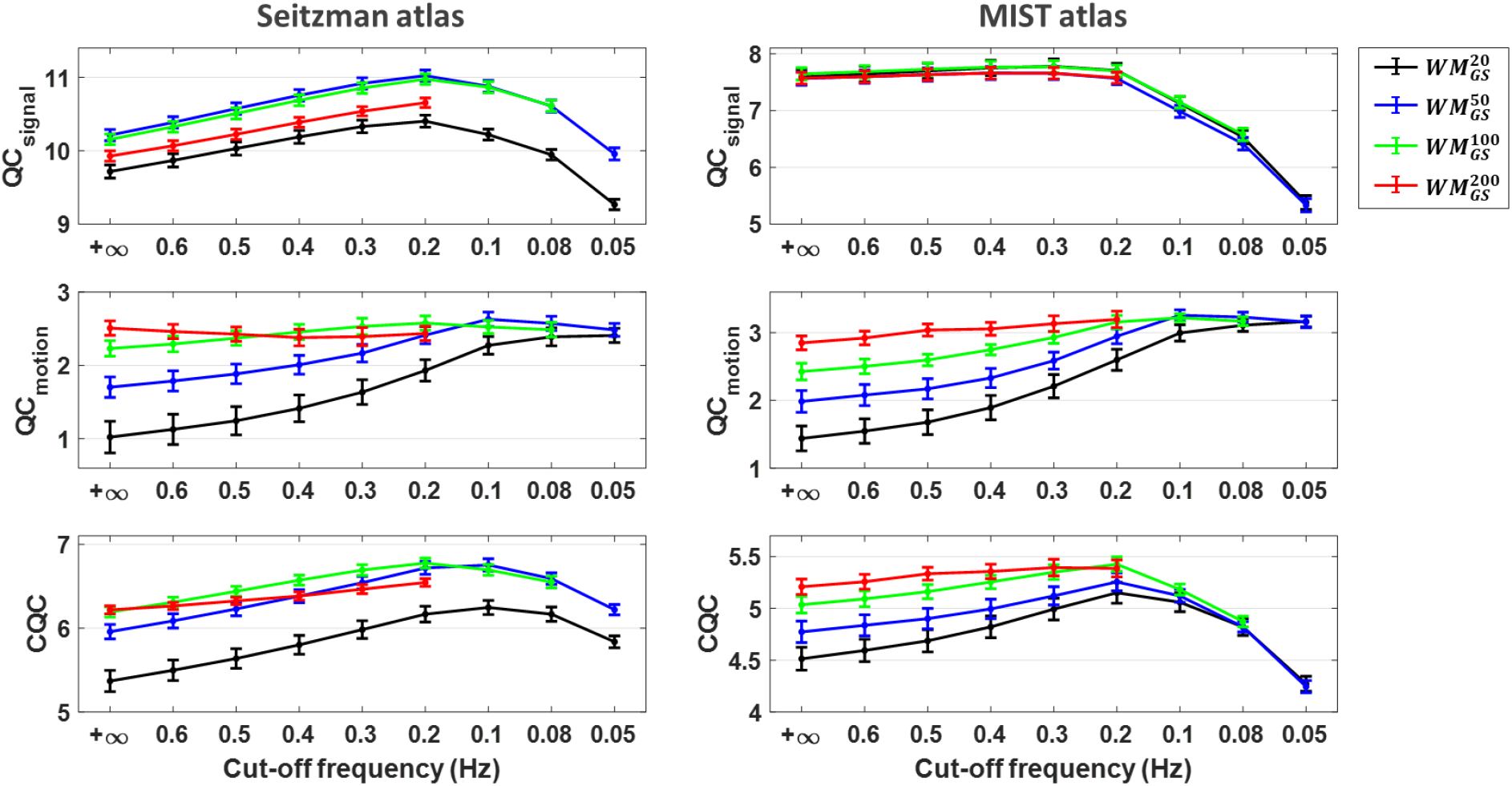
Effect of low-pass filtering in data quality for different cut-off frequencies on fMRI data in the Seitzman (left) and MIST (right) parcel space. For both atlases, the highest *CQC* score was achieved when low-pass filtering was done with a cut-off frequency of 0.2 Hz, and removal of nuisance regressors was done with 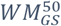 or 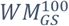. In the case of the Seitzman atlas, similar levels of *CQC* score were also achieved with 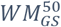 and 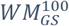 at a cut-off frequency of 0.1 Hz. For both atlases, mild variants of WM denoising (i.e. 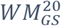 and 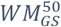) benefited significantly from low-pass filtering in terms of *QC*_motion_.

**Suppl. Fig. 20.**
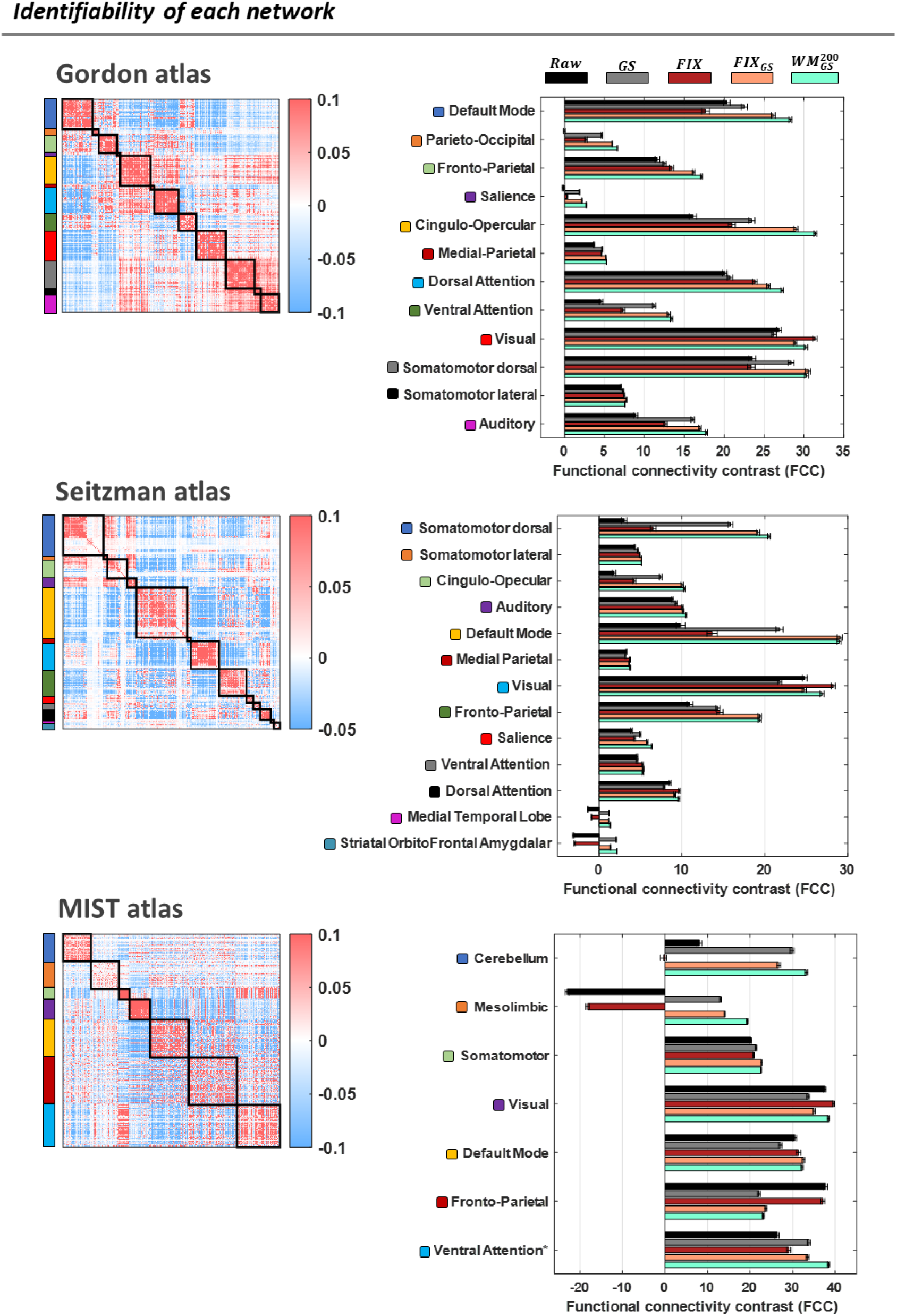
Identifiability of each network for the three examined functional atlases (Gordon, Seitzman and MIST). The *FCC* score of each network was defined as the *Z*-statistic of the Wilcoxon rank-sum test related to the null hypothesis that WNEs of the examined network and BNEs in the FC matrix are samples from continuous distributions with equal medians. In the case of the Gordon and Seitzman atlases, there was larger variability in *FCC* scores across networks rather than across pipelines, which might be due to the variability in the number of parcels of each network. In the majority of networks, pipelines *FIX*_*GS*_ and 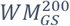 exhibited the highest *FCC* scores. * The last network in the MIST atlas, apart from the ventral attention network, consists also of the salience network, the basal ganglia and the thalamus.

**Suppl. Fig. 21.**
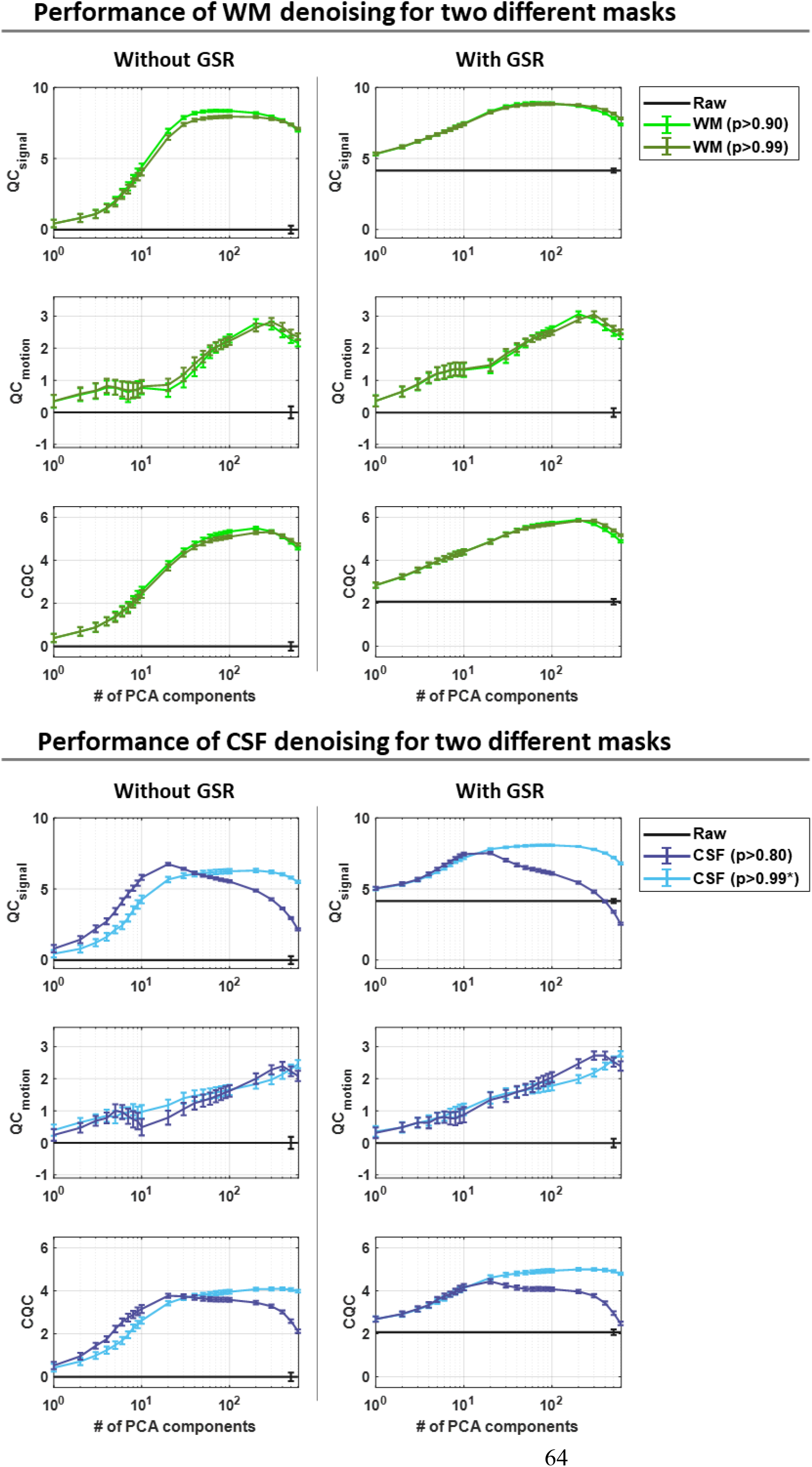
Performance of WM (top panel) and CSF (bottom panel) denoising for two different masks (Gordon atlas). For the case of WM denoising, using a stricter probability threshold for defining the mask of WM compartment (*p* > 0.99 instead of *p* > 0.90) did not have any significant impact on the performance of WM denoising. In contrast, for the case of CSF denoising with nuisance sets consisting of more than 20 regressors, a stricter probability threshold for defining the CSF mask (*p* > 0.99 instead of *p* > 0.90) led to substantially higher scores of *QC*_*signal*_. *For several scans, a probability threshold of *p* > 0.99 led to masks with an inadequate number of CSF voxels (<600) for performing principal component analysis (PCA), and thus, in those cases, the threshold was iteratively decreased (up to *p* < 0.90) until a sufficient number of CSF voxels was found.

**Suppl. Fig. 22.**
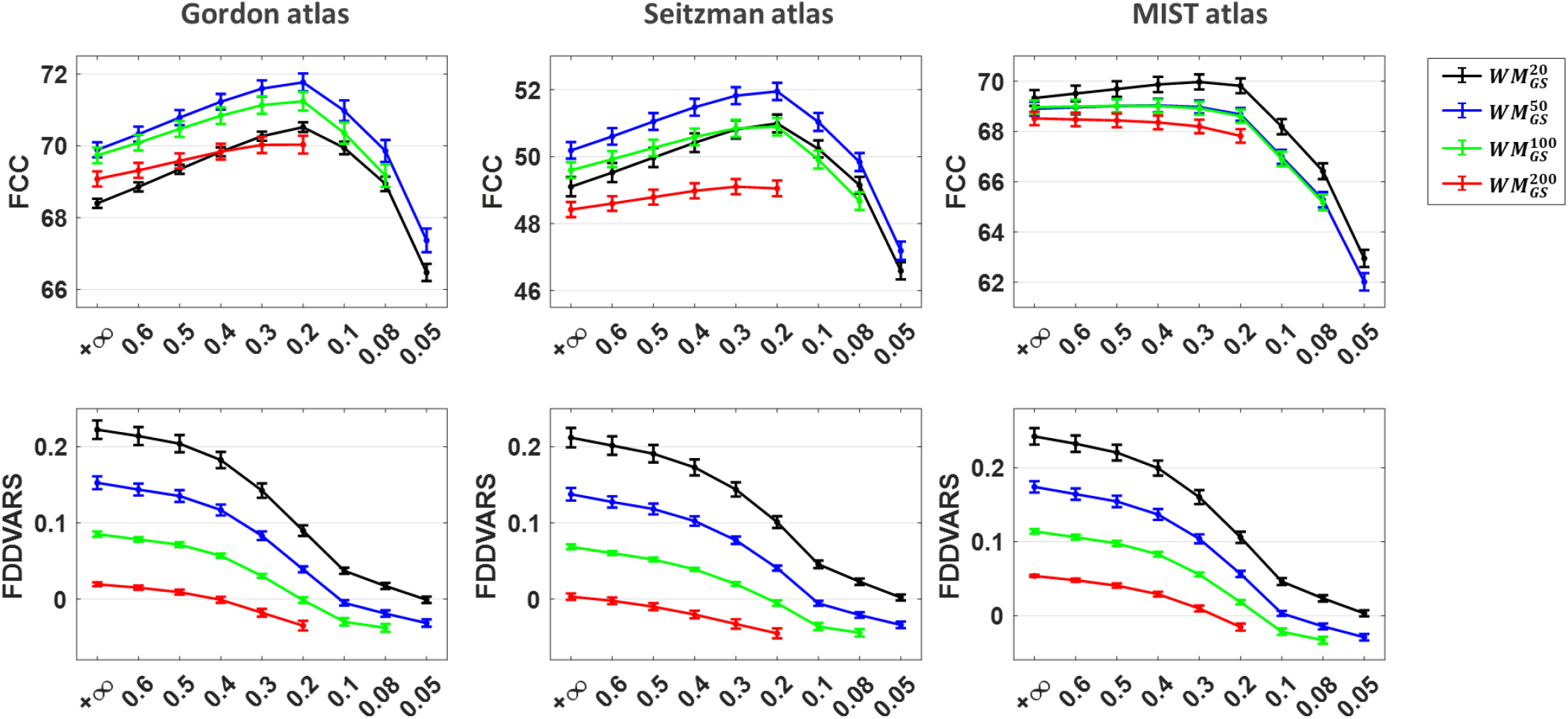
Effect of low-pass filtering in FCC and FDDVARS on fMRI data in the Gordon (left), Seitzman (middle) and MIST (right) parcel space. For the Gordon and Seitzman atlas, the highest *FCC* score was achieved with 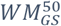 and a cut-off frequency of 0.2 Hz, whereas for the MIST atlas it was achieved with 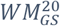 and a cut-off frequency of 0.3 Hz. For all three atlases, a cut-off frequency lower than 0.2 Hz led to a significant decrease in *FCC* score. In the case of *FDDVARS*, the pipeline that yielded the lowest absolute scores (i.e. lowest levels of motion) varied across the three atlases. However, for all atlases, 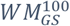 with low-pass filtering at 0.2 Hz led overall to scores very close to zero (i.e. low levels of motion artifacts).

